# A unifying theory explains seemingly contradicting biases in perceptual estimation

**DOI:** 10.1101/2022.12.12.519538

**Authors:** Michael Hahn, Xue-Xin Wei

## Abstract

Perceptual biases are widely regarded as a window into the computational principles underlying human perception. To understand these biases, previous work has proposed a number of conceptually different and even seemingly contradicting ingredients, including attraction to a Bayesian prior, repulsion from the prior due to efficient coding, and central tendency effects on a bounded range. We present a unifying Bayesian theory of biases in perceptual estimation. We theoretically demonstrate an additive decomposition of perceptual biases into attraction to a prior, repulsion away from regions with high encoding precision, and regression away from the boundary. The results reveal a simple and universal rule for predicting the direction of perceptual biases. Our theory accounts for, and leads to new understandings of biases in the perception of a variety of stimulus attributes, including orientation, color, and magnitude.

## Introduction

Human perceptual decision is often biased (1, 2, 3). For instance, a slightly tilted vertical bar is often perceived as more tilted than it really is (1). As key signatures of the underlying computational process, these biases are of tremendous interest to brain scientists, and have substantial societal implications (4, 5, 6, 7, 8, 9, 10) and clinical applications (11, 12, 13, 14). Over the past decades, Bayesian inference has emerged as a major theoretical framework for understanding perception, and more generally cognition (sometimes referred to as “Bayesian brain” hypothesis) (15, 16, 17, 18, 19, 20, 21, 22). The key tenet is that human perception and cognition reflect optimal combination of uncertain sensory input with prior expectations according to Bayes’ rule.

Previous studies have proposed several ingredients that may influence the bias of perceptual judgement. The first idea is that perception is biased toward prior expectation (23, 24, 25, 26, 27, 28, 29, 30, 16). In the Bayesian view, this can be formalized as a bias towards the peak of a prior distribution, *e*.*g*., the “slow speed prior” in motion (27, 31, 29), “light from above prior” in shape perception (24, 32, 28), and categorical priors in explaining biases in spatial perception (26, 33). A second idea is regression toward the mean: when the range of the stimulus is limited, the reported stimulus will be biased toward the center of the range (3, 34). Third, recent work considering Bayesian models constrained by efficient coding found that perception may be biased away from the prior expectation (35, 36). This idea explains why orientation perception exhibits biases towards oblique directions (37, 38, 39) even though these are least frequent in natural scene statistics (40, 41). Fourth, the particular form of loss function used by the observers may influence the estimation bias in perceptual tasks (42, 34).

So far, there is very limited understanding of how these disparate ingredients together determine perceptual biases, *e*.*g*., when should one expect an attractive (or repulsive) perceptual bias? This makes it challenging to interpret many previous results. Here we develop a theory that unifies all the aforementioned ingredients. Our analytical results provide an additive decomposition of perceptual biases under a wide class of loss functions, and arbitrary combinations of encoding models and decoding priors. The theory leads to a simple yet widely applicable rule for judging the direction of perceptual biases. This theory not only correctly accounts for prominent perceptual bias patterns reported in experimental data, but also provides a unified understanding of seemingly contradicting perceptual biases.

### Bayesian observer model with arbitrary prior and encoding

We first describe a general Bayesian observer model (Figure 1a), which serves as the basic framework for our results. Suppose a given stimulus θ follows the prior distribution *p*_*prior*_(θ). We assume that sensory encoding satisfies the general encoding model (43, 44, 36, 45, 46, 47)

**Figure 1:**
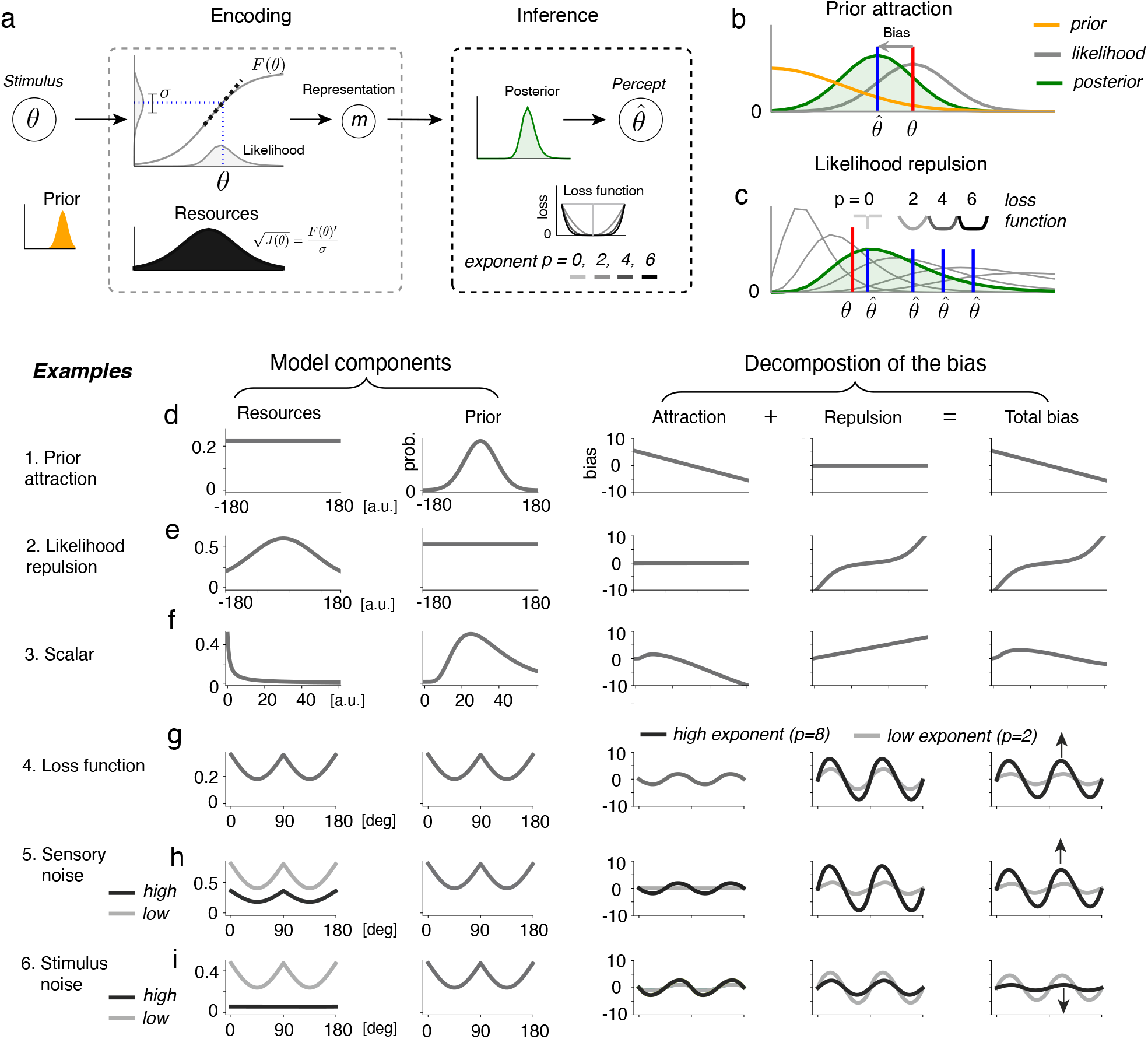
(a) The Bayesian observer model is specified by the prior, the allocation of coding resources, and the loss function. Observations are encoded by a nonlinear map, an abstraction of the neural representation, and subject to neural (sensory) noise. The precision of coding is described by the slope of the encoding function, which is proportional to the square root of the Fisher Information. Via Bayesian inference, an estimate is decoded as the minimizer of a loss function under the posterior. (b–c) The bias of the estimate consists of two components: (b) When the prior (orange) is nonuniform around a stimulus, the posterior (green) is shifted towards regions of high prior density compared to the likelihood (gray). The estimate (blue) is correspondingly drawn towards towards the peak of the prior. (c) When the encoding precision is nonuniform around a stimulus, both the likelihood and the posterior become asymmetric, with a long tail in regions with lower encoding precision. The encoding will on average be biased towards regions of low precision; the posterior will again have a long tail in those regions. On average across trials, the estimate is correspondingly drawn towards those regions. A higher exponent in the loss function (here *p* = 2, 4, 6) leads to higher tolerance of small errors, and increases this repulsive bias. (d–i) Examples (*p* = 2, unless otherwise indicated): (d) When the encoding is uniform, biases point towards the prior’s peak. (e) When the encoding is nonuniform, biases point towards regions with low encoding precision. (f) A scalar variable obeying Weber’s law, with a log-normal prior. Attraction and repulsion combine. (g) In circular stimuli, both the prior and the encoding may be periodic due to efficient coding for natural scene statistics (36, 50). Biases will point towards oblique directions when the encoding varies steeply, and towards the cardinals when it varies less. (g) The attractive component is independent of the loss function exponent *p*, whereas the repulsive component increases with the exponent. (h–i) Role of noise (*p* = 8, see SI Appendix, Figure S5 for other exponents): Increasing sensory noise (h) increases both attraction and repulsion. In contrast, external noise (i) applied to the stimulus increases attraction, but decreases repulsion.

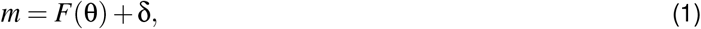

where the transfer function *F* maps the stimulus into a one-dimensional sensory space, an abstraction of the neural representation in the brain. δ is additive Gaussian noise in the sensory space. The transfer function *F* captures how coding resources are allocated. When it changes more quickly around a stimulus θ, more resources are allocated to stimuli around θ. Formally, this is described by the Fisher information, which is proportional to the square of the slope of the transfer function: 𝒥 = *F*′(θ)^2^*/*σ^2^, where σ ^2^ is the sensory noise variance. When *F* is proportional to the cumulative distribution of the prior, the system allocates the coding resources according to the principle of information maximization (48, 43, 49), a special case that was addressed in (36, 50). This model also subsumes models involving domain-specific encodings, such as logarithmic encodings of magnitude (51) and log-odds encodings of probabilities (52, 53). We will show that this general model (Eq. 1) allowing arbitrary combinations of prior and encoding enables us to obtain general insights regarding the factors determining the biases in perceptual estimation.

Given a sensory measurement *m*, the encoding model (Eq. 1) and the prior distribution together give rise to a posterior *P*(θ | *m*) over possible stimuli, from which a Bayesian point estimate 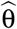 is decoded via minimization of a loss function (Eq. (5) in Materials and Methods). We consider the general family of *L*^*p*^ loss functions. In particular, *p* = 1 corresponds to the posterior median and *p* = 2 results in the posterior mean. The limit *p* → 0 leads to the posterior mode (MAP estimator). Intuitively, higher exponents lead to loss functions that are more tolerant of small errors.

#### Additive decomposition of estimation biases

We analytically derive the biases that correspond to different combinations of prior, encoding and loss function. We find that, for the model defined in Eq. (1), the bias of the estimate 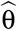 for *L*^*p*^ loss (*p* ≥ 1 an integer) can be written as:

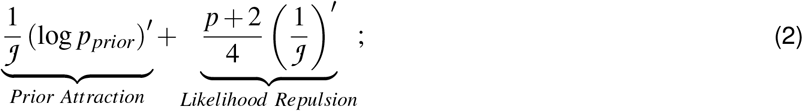

and, for the MAP estimator

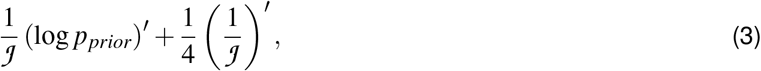

where 𝒥 is the Fisher information, which is a function of θ. These analytical results hold when noise is small (an assumption also made in, e.g. 29, 36, 54). Numerical simulations show that qualitative patterns predicted remain valid for larger noise (see SI Appendix, Figure S4). The main deviation at higher noise levels is that biases grow more slowly as noise increases, because high noise effectively ‘smooths out’ local variability in prior and encoding precision. We prove an analogous decomposition for an alternative model sometimes assumed in the literature (e.g. 29, 34, 55, 41), in which sensory noise is Gaussian in the stimulus space with stimulus-dependent variance (See SI Appendix, Section S3.5 for details).

Our results provide a general expression for the bias under arbitrary combination of prior distribution, resource allocation and loss function. The first term in the equations describes attraction to the prior: the bias is proportional to the slope of the logarithmic prior, increased in regions with low encoding precision, consistent with the general idea that reliance on the prior increases when sensory input is less certain (e.g. 27, 29, 54). The second term describes a repulsive bias away from regions of high encoding precision, scaling with a factor depending on the loss function. This additive decomposition into attractive and repulsive components clarifies how prior and likelihood repulsion (induced by encoding heterogeneity) jointly determine perceptual biases under our general model. Notably, the likelihood repulsion scales with the exponent of the loss function, so that repulsive biases are increased when the loss function is tolerant of small errors, whereas prior attraction is largely independent of the loss function.

The theory is illustrated using representative examples in Figure 1d–i. Traditional Bayesian models assume a nonuniform prior and a uniform encoding (Figure 1d); this gives rise to attraction towards the prior peak. When, on the other hand, the prior is uniform but the encoding precision is not uniform, a repulsive bias away from regions of high precision is predicted (56) (Figure 1e). In recent work, considerations of efficient coding have led to the proposal of models where the encoding precision is proportional to a power function of the prior (36, 50, 55, 57, 58). In contrast, our theory allows for situations where the prior and encoding are arbitrary and both nonuniform, so that both attractive and repulsive components will come together. Figure 1f considers the case of a scalar variable whose encoding follows Weber’s law, and subject to a unimodal prior. Here, the bias will point towards the prior peak, with strength varying with the stimulus.

#### Previous results as special cases

Special cases of Eq. (2–3) recover various previous analytical results (see SI Appendix, Section S3.1.2). In the special cases *p* = 0 and *p* = 2, we recover results from (54, 59). Some prior work has considered coding models where the Fisher information is matched to the square (36) or another power of the prior, which can be justified in terms of efficient coding. In this setting, our results lead to a general description of the bias in terms of the power *q* linking prior and encoding, not previously known beyond the special cases *q* = 2 (50) or *p* = 2 (54, 61). Recent work found that, in a broad range of experiments, the discrimination threshold and perceptual bias follow a lawful relationship such that the bias is proportional to the slope of the squared threshold, which can be accounted for by Bayesian inference based on efficient coding (50). Our results entail that this psychophysical law holds if and only if the threshold is proportional to a power of the prior (see SI Appendix, Section S3.4).

### A simple and universal rule for judging the direction of perceptual biases

Our theory leads to a simple and universal rule to predict the direction of the bias, which is one of our key results. We find that, when noise is small, the bias for a given *L*^*p*^(*p* > 0) loss function is predicted to have the same sign as:

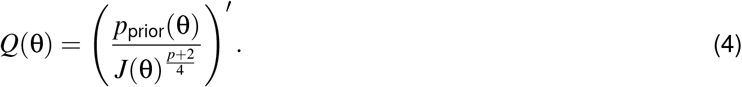

As the quantity in the denominator quantifies the encoding precision for each stimulus, we refer to this as the prior/precision ratio rule, or P/P ratio rule for simplicity.

Consider the case where the Bayesian estimate is taken to be the posterior mean (*L*^2^ loss). In this case, the ratio rule can be written as 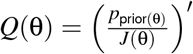. Assuming the neural code maximizes mutual information, previous work (43, 57, 36, 62) found 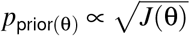, thus the bias has the same sign as 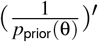, precisely the opposite of the direction of the prior belief. Thus, a repulsive bias is predicted (36).

More generally, the ratio rule shows precisely that the direction of the perceptual bias depends on how fast the prior distribution and the encoding precision change relative to each other (Figure 2): The component that changes faster than the other will dominate the biases. This ratio rule provides a new insight that may reconcile a range of previous experimental observations: One plausible explanation of seemingly contradicting observations of either attractive or repulsive biases is that the balance between the prior and precision may differ across different experimental scenarios – a hypothesis we will test later with experimental data.

**Figure 2:**
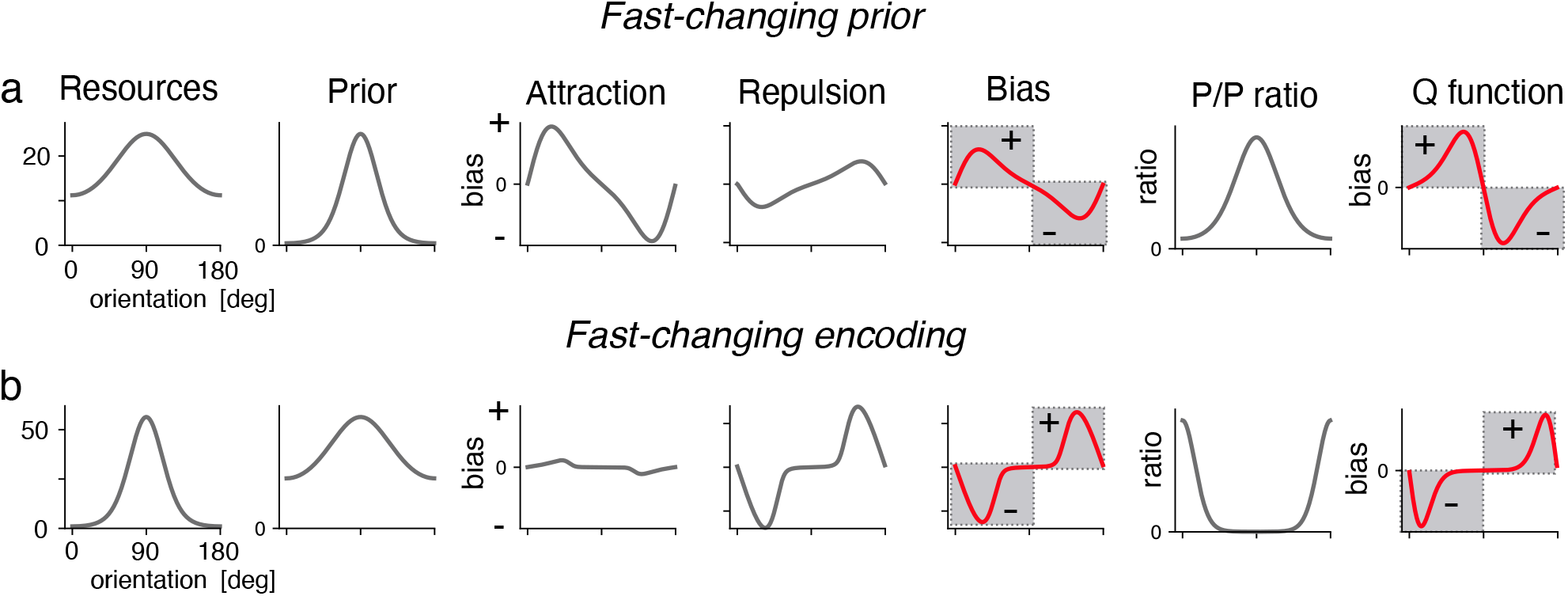
The P/P ratio rule determines the direction of the overall bias, when the prior (a) or the resource allocation (b) changes more quickly over the stimulus space. In addition to the model components and the bias components (*p* = 2), we plot the ratio between prior and Fisher information, and its derivative *Q*(θ) (Equation 4). As predicted by our theory, *Q*(θ) predicts the direction of the bias: Attraction dominates when the prior changes faster than the encoding (a), repulsion dominates when the resource allocation changes faster (b).

### Extensions of the theory

#### Stimulus boundary induces biases toward the interior

The results so far apply to stimuli in the interior of the stimulus space, away from any boundary. Effects of the boundary may become important in experiments that measure the perceptual biases using a relatively small bounded range (63) or ratings on a bounded scale (45). We consider two ways to model the effect of boundaries, and will compare these two schemes quantitatively using experimental data. First, boundaries may lead to a truncation of the posterior (34). We analytically derived the impact of the boundary on the bias in this case (see Theorem 3 in Materials and Methods): In the vicinity of the boundary, biases are dominated by a regression effect into the interior of the range; this effect disappears as one moves into the interior. Alternatively, when stimuli are limited to an interval, subjects might also develop a smoothly-changing prior, such as a Gaussian prior, as an approximation to the sharply-dropping prior at the edge (Figure 1f). For this case, the predicted bias is readily accounted for by Eqs. (2 &3). These considerations have implications for interpreting experimental data: Biases measured in experiments with bounded range will likely exhibit systematic biases toward the center of the range, in particular when noise is substantial. Hence, perceptual biases in experiments with bounded range may exhibit patterns quite different from those found with circular variables.

#### Stimulus noise modulates estimation biases

Experimental work has shown that stimulus (external) noise also influences perceptual biases (39, 41, 64). For instance, in perception of orientation, adding Gaussian stimulus noise biases responses toward the cardinal orientation (39, 41), whereas sensory noise increases biases toward the obliques (38). To understand these results, we extended the theory to address stimulus noise (see Theorem 2 in Materials and Methods). Our theory generalizes simulation results in (36), and provides a precise analytical characterization of how the stimulus noise affects the perceptual biases and how its effect differs from that of the internal noise. The comparison between the two types of noise is illustrated in Figure 1h–i. When adding stimulus noise, the representation becomes noisier and reliance on the prior increases. Thus, prior attraction increases with both sensory and stimulus noise. Simultaneously, and more subtly, stimulus noise reduces likelihood repulsion when *p >* 2, because it broadens the posterior and reduces its asymmetry. Thus, the theory predicts that, in settings where prior and encoding are aligned, increasing sensory noise makes biases more repulsive, while stimulus noise makes biases more attractive.

### Applications to data reveal both domain-specific and general insights

We applied our model to a number of datasets collected in previous experiments (65, 39, 38, 63, 66, 67), ranging from color and orientation perception to the estimation of numerosity. To fit our general Bayesian model to the data, we developed a general numerical fitting procedure that determines the model components, including prior, encoding, and noise magnitude, by maximizing the likelihood of trial-by-trial response data. Hence, the model is fitted to account for the full response distribution conditioned on the stimulus, taking both the perceptual bias and the response variability into account.

#### Application 1: Orientation estimation

Perception of orientation is generally biased, with some studies finding biases towards cardinal orientations (41), while others reported systematic biases toward the obliques (37, 38, 39, 68). Bayesian models have been applied to understand these observations (41, 39, 36). In particular, Bayesian models based on efficient coding have been shown to *qualitatively* reconcile these contradictory findings (36). However, it remains unclear whether this type of model could quantitatively explain the data, and if so, how important each of model component is.

We start by re-analyzing the data from de Gardelle et al. (38). In this experiment, stimulus noise was manipulated (via varying the length of stimulus presentation time) while observers were asked to reproduce the orientation of individual Gabor stimuli. Presupposing a well-established orientation prior based on natural image statistics (39, 41, 36), we fitted our model (defined by Eq. 1) to these data (Fig. 3a–c). The fitted model exhibits decreased coding precision around the oblique orientations, in agreement with the oblique effects long documented in experimental work (69, 70). As predicted by our theory, the measured biases pointed away from cardinal directions, and increased with sensory noise. Meanwhile, Tomassini et al. (39), Girshick et al. (41) measured biases in orientation perception at two levels of stimulus noise, finding that this oblique bias is reduced with larger stimulus noise. This pattern is well as predicted by our theory (model fits in Fig. 3f–h).

**Figure 3:**
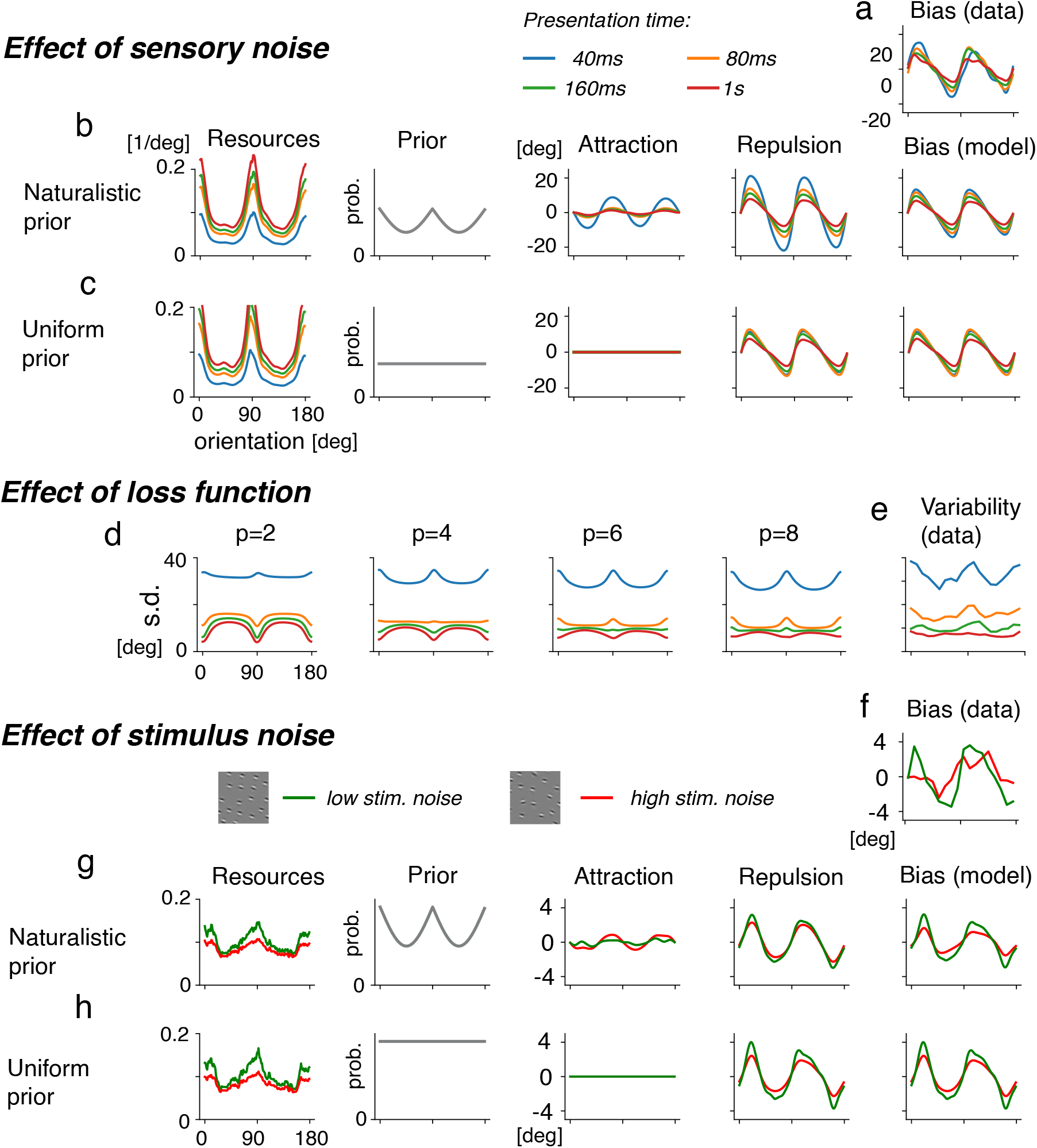
Bias in orientation estimation. (a–c) Bias in orientation perception (data collected by 38), with naturalistic or uniform prior. The naturalistic prior is based on natural image statistics, *p*(θ) ∝ 2 − | *sin*(θ) | (41, 36). We show results for the best-fitting exponent, *p* = 8. The bias is dominated by likelihood repulsion. Decreased exposure times correspond to increased sensory noise, leading to increased biases. The prior plays a small role; a uniform prior achieves approximately optimal fit (see SI Appendix, Figure S10). (d–e) While most loss functions can reproduce the bias (SI Appendix, Figure S15), small values of *p* exhibit poor fit to the observed variability. The data (e) are best accounted for using a loss function that is tolerant of small nonzero errors (e.g., *p* = 8). (f–h) Effect of increasing stimulus noise (data collected by (39); schematic sample stimuli are adapted from (39)). We fitted the model both with the prior based on natural image statistics (g) and a uniform prior (f), again at *p* = 8. In agreement with our theory, higher stimulus noise (red) decreases repulsion and increases attraction, leading to an overall decrease in bias magnitude.

By directly fitting to trial-by-trial data, these results reveal two novel insights that go beyond previous qualitative models of orientation perception (36). First, the inferred loss function exhibits a higher exponent (*p* = 8) than previously assumed. Estimators with lower exponent, as assumed in previous work (41, 36) provide poor fit (Fig. 3d); an observation recently also made by (71). The data are better described by higher exponents; our modeling thus indicates that, in these orientation estimation tasks, human observers are much less sensitive to small errors than previously thought. Second, the encoding heterogeneity plays an important role in accounting for both the oblique biases and the reduction of oblique biases when adding stimulus noise. When only sensory noise is present, inhomogeneity in encoding leads to an asymmetric posterior (Fig. 1c) accounting for the oblique biases. With increasing stimulus noise, our theory predicts both reduction of likelihood repulsion and increase of prior attraction (Fig. 1i), pulling biases towards the cardinals. Strikingly, in both datasets (38, 39), a model with a uniform prior and appropriate encoding and loss functions achieves at least as strong a model fit, and freely fitting a prior indeed leads to an approximately uniform fit (SI Appendix, Figure S12 and S23).

These results highlight the importance of loss function and encoding heterogeneity in orientation perception. The reported orientation biases can be readily explained by the interactions between (sensory and stimulus) noise and heterogeneity of encoding, without considering attraction to prior expectations. This small role of the prior is surprising, and might be related to the fact that stimulus distributions were uniform in these experiments. These findings represent a major departure from previous models which ascribed the impact of stimulus noise on the bias to an increased role of the prior (39, 41).

#### Application 2: Perceptual learning of motion direction

Next, we investigate how statistical learning of a short-term prior may influence the perception of an important circular stimulus variable, *i*.*e*., motion direction. Similar to orientation, motion perception exhibits oblique effects (72, 73) and biases away from the cardinal directions (74). Studies have used perceptual learning paradigms to study motion perception (75, 67). In these studies, subjects were asked to estimate the motion direction of a coherently moving point cloud. On each trial, the movement direction was drawn from a bimodal distribution whose peaks were separated by 64 degrees around a central direction that was randomly chosen for each subject. In order to determine whether the perceptual learning-induced effects could be explained by our theory, we reanalyzed data collected by Gekas et al. (67). Assuming, first, the oblique effect in encoding precision and, second, that the subjects learned the bimodal stimulus distribution, our theory predicts that biases will combine repulsion (away from the cardinal directions) and attraction (towards the short-term motion prior). We note that the pattern of bias and variability reported in (67) was quite complex, and the Bayesian models considered in (67), which assumed homogeneous encoding, showed qualitative mismatch with the data. We hypothesize that by considering both encoding heterogeneity and short-term prior, a clearer explanation would emerge.

Because the experimentally induced stimulus distribution varies between the subjects in this experiment, we fitted a mixed-effects version of our model that includes subject-specific adjustments to encoding and prior (see Materials and Methods). The results provide strong support for our prediction. For subjects where the prior was centered close to a cardinal direction, prior attraction and likelihood repulsion consistently pointed in the same direction, leading to a large bias (Figure 4b). For subjects where the distribution of motion direction was centered close to the oblique, the components partially pointed in opposite directions and the magnitude of the bias was reduced (Figure 4d). The recovered resource allocation is comparable among the subjects who were presented with different motion statistics during the task, consistently peaking at cardinal directions (Figure 4a). In contrast, the recovered priors differ systematically and reflect the experimentally-defined motion statistics (see also SI Appendix, Figure S28). Quantitative model fit strongly improved over a model assuming a uniform encoding (as assumed by (75, 67), difference in negative log-likelihood [NLL]: > 50). Overall, these results show that (i) both encoding heterogeneity and short-term priors in this perceptual learning task are important for understanding the reported errors in motion direction; (ii) learning short-term statistics leads to a major change in the subject’s prior, but less so in their encoding.

**Figure 4:**
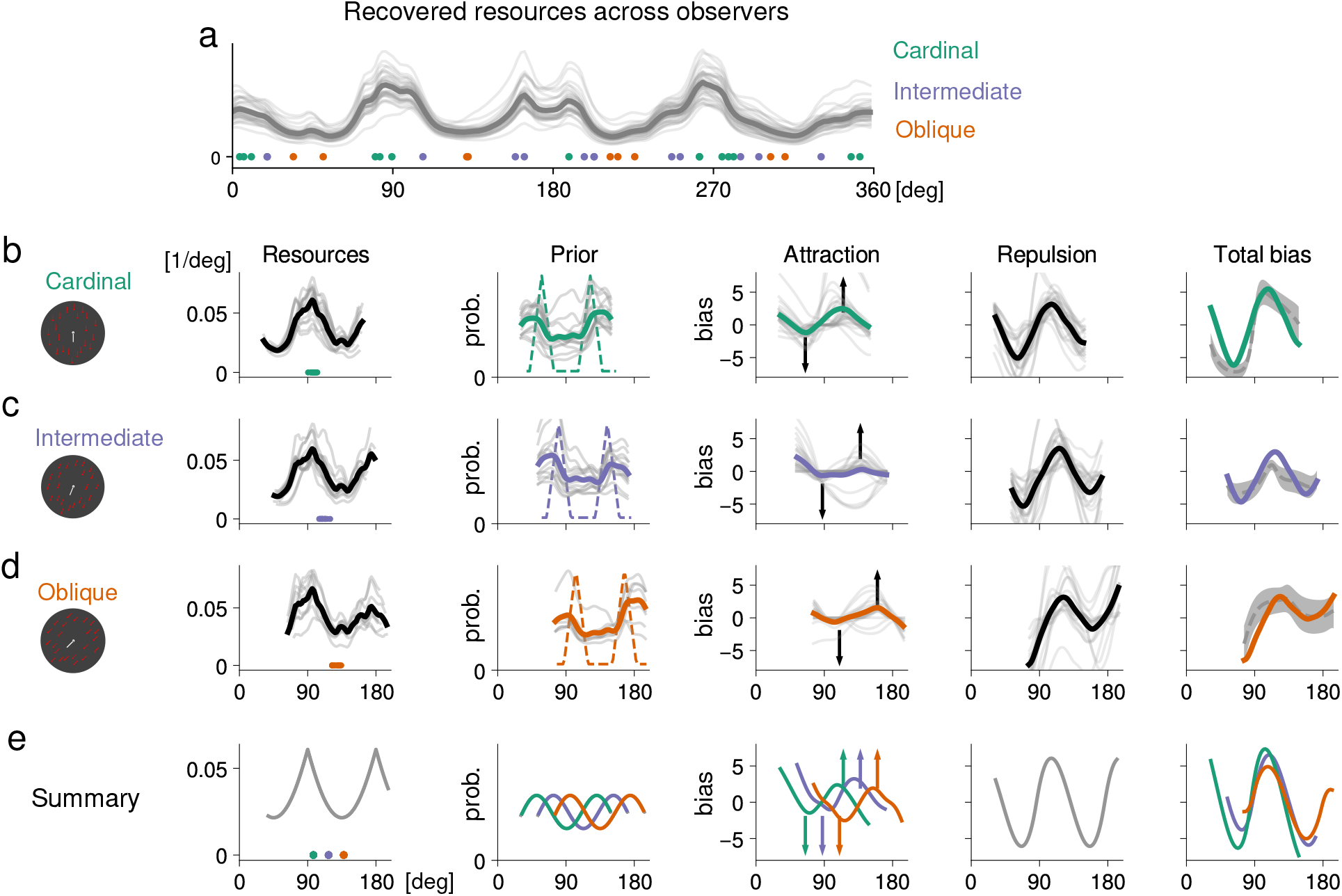
Effect of perceptual learning on motion perception. Gekas et al. (67) exposed 36 subjects to moving point clouds, whose movement directions were sampled from a bimodal distribution centered at different angles for different subjects. Sample stimuli are schematic; contrast is increased here for visibility. We fitted our model with per-subject adjustments to encoding and prior. (a) Across subjects, a resource allocation peaking at cardinal angles is identified. Dots indicate the central directions for each of the subjects, colored depending on whether they are close to the cardinal or oblique directions, or in between. (b–d) We separately consider subjects based on the three-way grouping from (a). In order to compare biases across subjects, we transformed the coordinates for each subject so that the central direction was in [90^◦^, 135^◦^]. We show results fitted for each subject (faint), and averaged (solid). The fitted resource allocation reflects increased coding precision at cardinal directions; the fitted prior (solid) approximately reflects the bimodal stimulus distribution (dashed). Measured biases (dashed=human with standard errors over subjects, solid=model) reflect the impact of both encoding inhomogeneity and subject-specific priors. The two components reinforce each other for subjects in the cardinal group, while biases are reduced in the oblique group. (e) Idealized pattern across the three groups of subjects. Resources and likelihood repulsion are approximately the same across subjects. Different priors give rise to different prior attraction biases; the overall bias is reinforced for the cardinal group and reduced for the oblique group.

#### Application 3: Regression towards the mean in magnitude perception

Perhaps the most pervasive and long-documented bias in perceptual estimation is the central tendency effect: Humans exposed to magnitudes from a bounded range produce estimates biased towards the center of the range (3, 76, 77). Recent studies also found that internal noise (78, 64, 79, 80) and stimulus noise (64) increase the regression effect. Bayesian models (34, 79, 51, 81, 66) have been proposed to account for these effects. To model a limited stimulus range, some of the semodels assumed a flat prior supported on the relevant range (34), and others assuming a smooth varying prior that peaks in the interval (51, 82). Which mechanism better accounts for the data remains unclear. In addition, these kinds of tasks prominently exhibit a scaling law (variability increases with the mean; or equivalently, a constant Weber fraction), raising the question of how much the encoding heterogeneity contributes to the reported biases.

To dissect these factors, we first fit the data collected by Xiang et al. (66) using our framework. In this task, 300 subjects estimated the number of dots on a screen; within each block, dots were uniformly distributed in a range of width 30. In model fitting, we constrained the encoding precision to be consistent with Weber’s Law. Comparing models based on two different priors considered in previous studies (flat as in (34), normal in sensory space as in (51)), we find that the latter gives a much better fit (Fig. 5) than a flat prior. When allowing the prior to freely vary for each interval, we recover a model consistent with Gaussian noise in sensory space, with similar quantitative model fit (see SI Appendix, Section S30). As the encoding satisfies Weber’s law, the theory also correctly predicts that the central tendency effect will be larger for blocks with larger mean numerosity, as observed in (66). Examining the fitted models (Figure 5), we find that the encoding heterogeneity leads to a systematic bias towards larger magnitude, yet most of the bias is accounted for by prior attraction. This is expected from the theory, because the prior on this bounded range varies much faster than the encoding precision.

**Figure 5:**
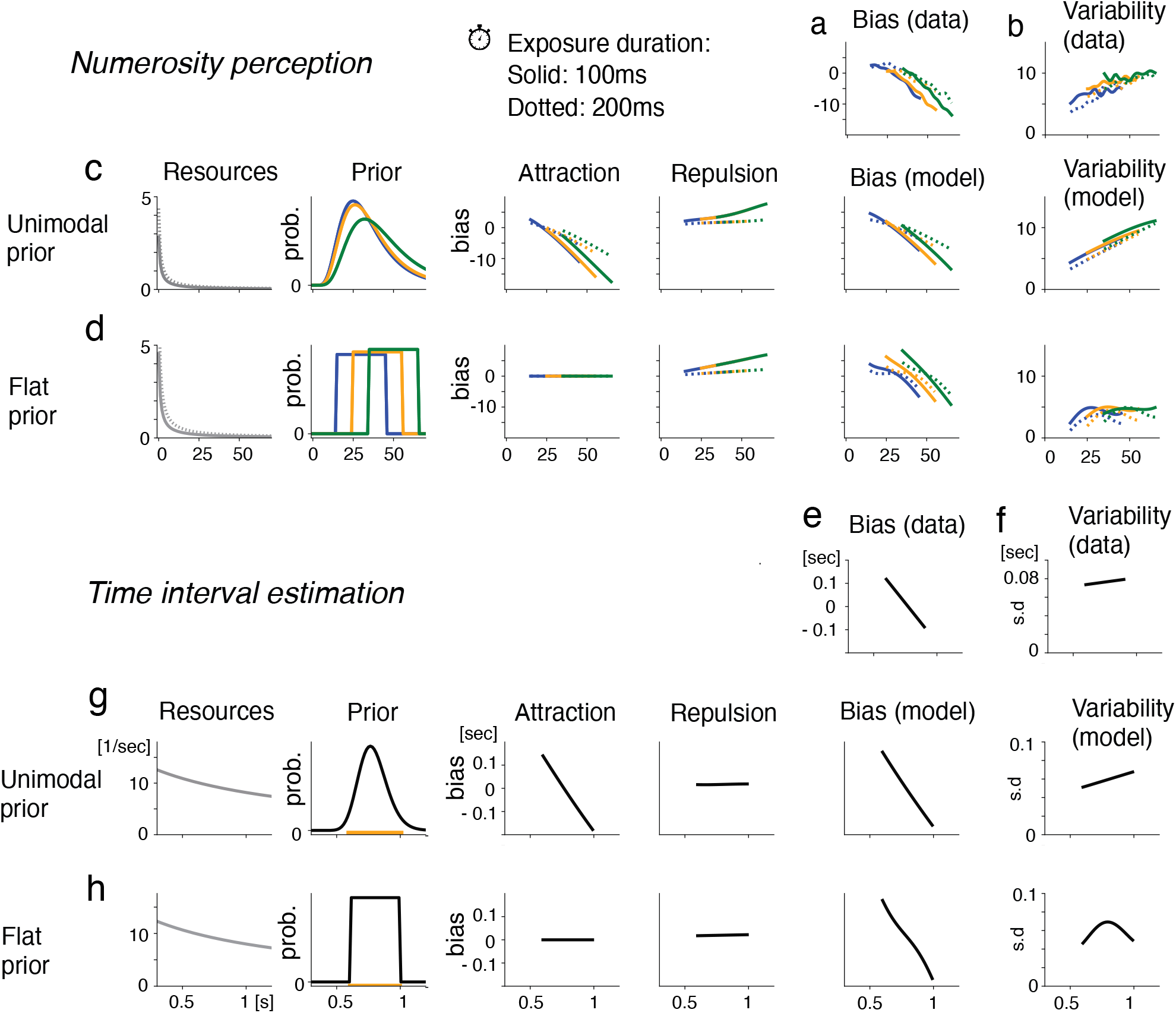
Central tendency effect in numerosity perception (a–d) and time interval estimation (e–h). (a–b) Xiang et al. (66) exposed subjects to point clouds with varying numbers of points, within a range that varied between blocks. We show fits for three of the 21 ranges considered (see SI Appendix, Figures S30–S34 for the other ranges), either with a unimodal prior that is Gaussian in sensory space (c), or a flat prior reflecting the true stimulus statistics (d), as assumed in some prior models (34, 63). We show fits at *p* = 0, results at other exponents are similar (SI Appendix, Figures S31–S34). In each range, biases towards the center of the range are observed. Increasing exposure duration (dotted) decreases sensory noise and hence biases. The central tendency effect is accounted for by prior attraction under the unimodal prior, and by boundary effects under the flat prior. A flat prior incorrectly predicts the variability to peak in the center. The unimodal prior achieves near-optimal fit (see SI Appendix, Figure S30). (e–f) Remington et al. (63) asked subjects to reproduce the time interval between two flashes on a screen. Stimuli were uniformly distributed within the interval indicated by the orange bar. We show fits at *p* = 2 (g–h), results at other exponents are similar (SI Appendix, Figure S35). As in (a–d), a flat prior incorrectly predicts the variability to peak in the center. Freely fitting the prior leads to a similar unimodal fit (see SI Appendix, Figure S36).

To verify robustness of these findings, we replicated these analyses using data from a time interval estimation task (63), and obtained largely similar results (Figure 5 e–h and SI Appendix, Figure S35 and S36). In previous modeling of biases in time interval estimation, Jazayeri and Shadlen (34) found that the squared error loss (*p* = 2) led to a much better fit compared to the maximum likelihood estimate or the MAP estimator (*p* = 0) in a time interval estimation task. We replicated these results on the dataset from (63) for the flat prior as assumed by (34) (SI Appendix, Figure S39). However, with the unimodal priors considered above, both estimators led to similar fit (and better fit compared to that of the flat prior), and model fit did not vary much with the exponent (SI Appendix, Figure S35). These observations are consistent with our theory, as the model fits suggest that the biases in these data are dominated by prior attraction, which is largely independent of the loss function according to Eq. (2).

These results provide new understanding of a classical effect. They suggest that the central tendency effect observed in estimation of scalar variables is well accounted for by a log-normal prior adapting to the stimulus range. The observers did not appear to learn a flat prior perfectly reflecting short-term statistics. While the encoding heterogeneity leads to a small bias towards larger values, this component is small compared to the prior attraction. While an abrupt drop-off of the prior does not provide a good model of the central tendency effect, we note that such an effect may still be observed when stimuli are close to a strict boundary of the range of allowed responses. See SI Appendix, Section S1.2, for a relevant example of subjective value ratings on a bounded scale (45).

#### Application 4: Categories in color perception

Experimental work has reported systematic biases in the perception of color hue (e.g. 65, 83), which has been explained in terms of representations involving discrete color categories (e.g. 65, 83). In the Bayesian framework, categories can be conveniently formalized as priors peaking at the most typical elements of a category (80), and the observed color bias may be explained by a bias toward these exemplars. Through re-analysis of previous data, a radically different explanation emerged, as described below.

We fitted our model to the trial-by-trial responses collected in Bae et al. (65) (Figure 6a). We find that the model explains the color biases as a combination of likelihood repulsion and prior attraction: Resource allocation is periodic across the color wheel with four peaks, creating an overall periodic bias pattern. The fitted prior peaks in the yellow-green range; attraction to this prior peak accounts for a negative bias in the blue range. Qualitative fit was similar across exponents *p* ≥ 0 (SI Appendix, Figure S41).

**Figure 6:**
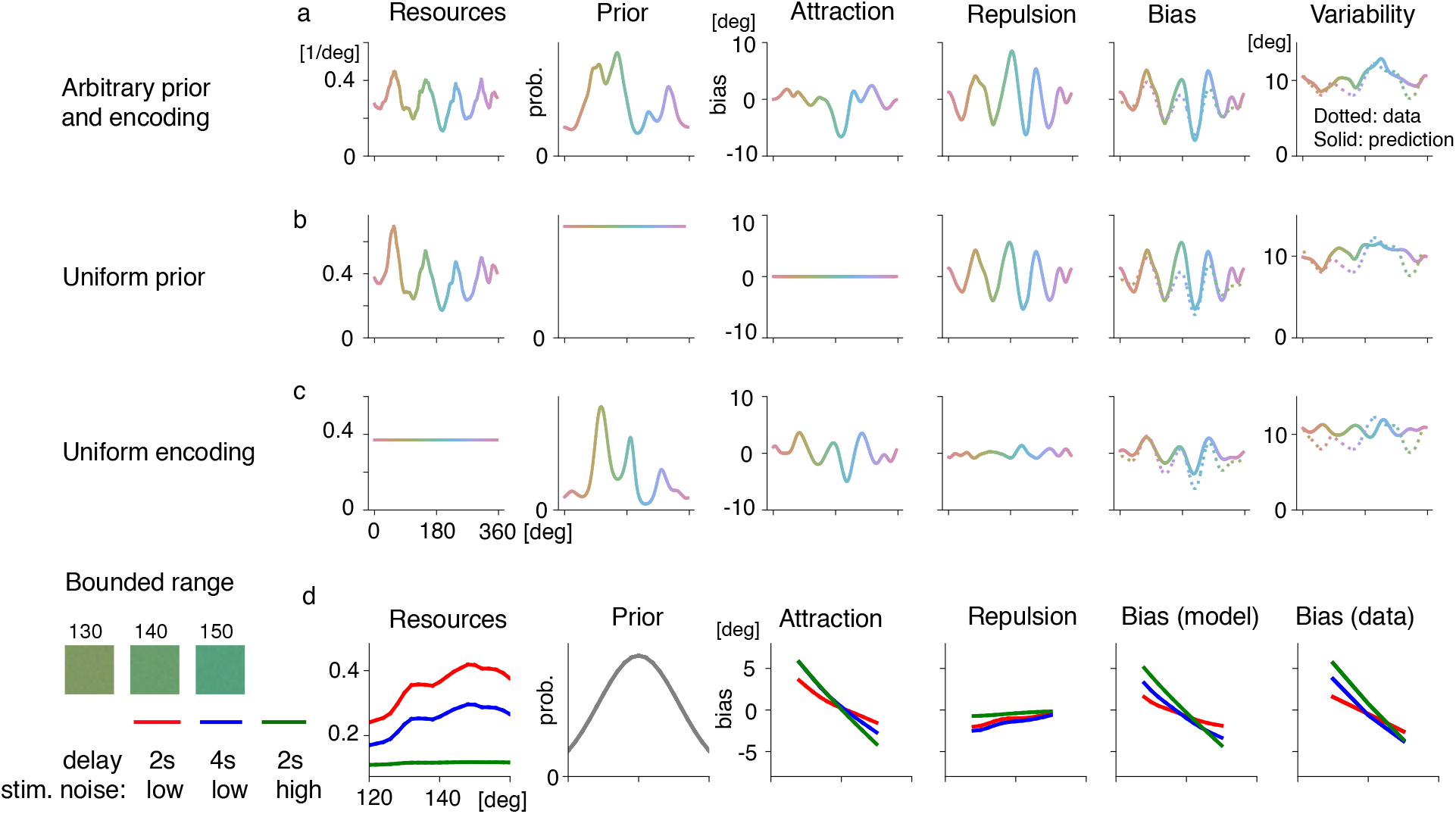
Bias in color hue estimation. (a,b,c) Bae et al. (65) asked subjects to reproduce a color hue on a color wheel (undelayed estimation, N=7). (a) The fitting procedure identifies a periodic pattern of resource allocation, and a prior peaking in the yellow range. We show fit at *p* = 4; results are similar across different exponents (SI Appendix, Figure S41) and for three further subjects where response was delayed (SI Appendix, Figure S45). The observed bias pattern (dotted) is mostly accounted for by likelihood repulsion; prior attraction accounts for relatively negative biases in the green-blue range. (b) A model with a uniform prior can qualitatively reproduce bias and variability, though fit is slightly inferior. (b) A model relying entirely on a prior can match the bias, but does not reproduce the variability pattern. (d) Olkkonen et al. presented subjects with stimuli taken from a small section on the color wheel. Encoding resources are from the fit in (A); a unimodal prior is assumed based on model fits to other datasets with small stimulus ranges (Figure 5). Parameters were chosen to reproduce reported biases and thresholds, with the same loss function (*p* = 4) as in (A). The resulting central tendency effect (red) is increased when increasing the delay after presentation (increasing internal noise, blue) or adding external noise to stimuli (green).

To better understand the relative importance of the prior and the encoding, we fitted two additional models. A model ablating the prior (Figure 6b) predicts a similar overall bias pattern, although it cannot account for the trend towards negative biases in the blue region. A model ablating the nonuniformity in encoding (Figure 6c) can account for the pattern of the bias, but provides a rather poor characterization of variability of the response; and correspondingly has inferior cross-validated likelihood (see SI Appendix, Figure S40). Together, these results suggest that while both prior and encoding heterogeneity are needed for fully accounting for the pattern of the color biases, the latter accounts for a larger fraction of the biases. We found similar results when analyzing data from 3 further subjects where exposure was followed by a delay (see SI Appendix, Figure S45).

Our modeling can be thought as an instantiation and re-evaluation of prominent Bayesian models of categorical perception (80). Our results show that a model building on a categorical prior is not favored by the color hue data. To the extent that the observed biases are related to color categories, such categories would determine biases via inhomogenous encoding, not the prior belief.

Our theory further predicts that attractive biases will become more relevant when color hues are restricted to a small interval. This is supported by data from Olkkonen et al. (64), who presented subjects with stimuli taken from a small section on the same color wheel (Figure 6d). Biases point into the interior of the range and increase not only with sensory noise, but also with external noise (64). This result is compatible with an interpretation as attraction to a prior, overriding the repulsive effect on a small range where the prior varies rapidly.

### General insights emerging from the analyses

Figure 7 summarizes some of main results from the analyses of these datasets. Importantly, in all cases, the results can be well captured by our P/P ratio rule (Figure 7a). Combining the results from different applications reveals major differences between circular variables and scalar variables. For circular variables, we find that encoding heterogeneity is often key in driving the observed biases, and that prior attraction can play a surprisingly small role. This stands in marked contrast to traditional models emphasizing attraction to prior expectation. We summarize the role of attractive and repulsive biases across the datasets considered in this work in Figure 7b, confirming that repulsive biases play a larger role in accounting for the overall biases in the four circular datasets, and a smaller role in the two scalar datasets. Consistent with our theory, the loss function plays an important role in accounting for perceptual data based on circular variables, and much less so for that of scalar variables (Figure 7c). Taken together, these results clarify a large and disparate literature, and reveal novel insights into the factors dominating the perceptual biases under different experimental conditions.

**Figure 7:**
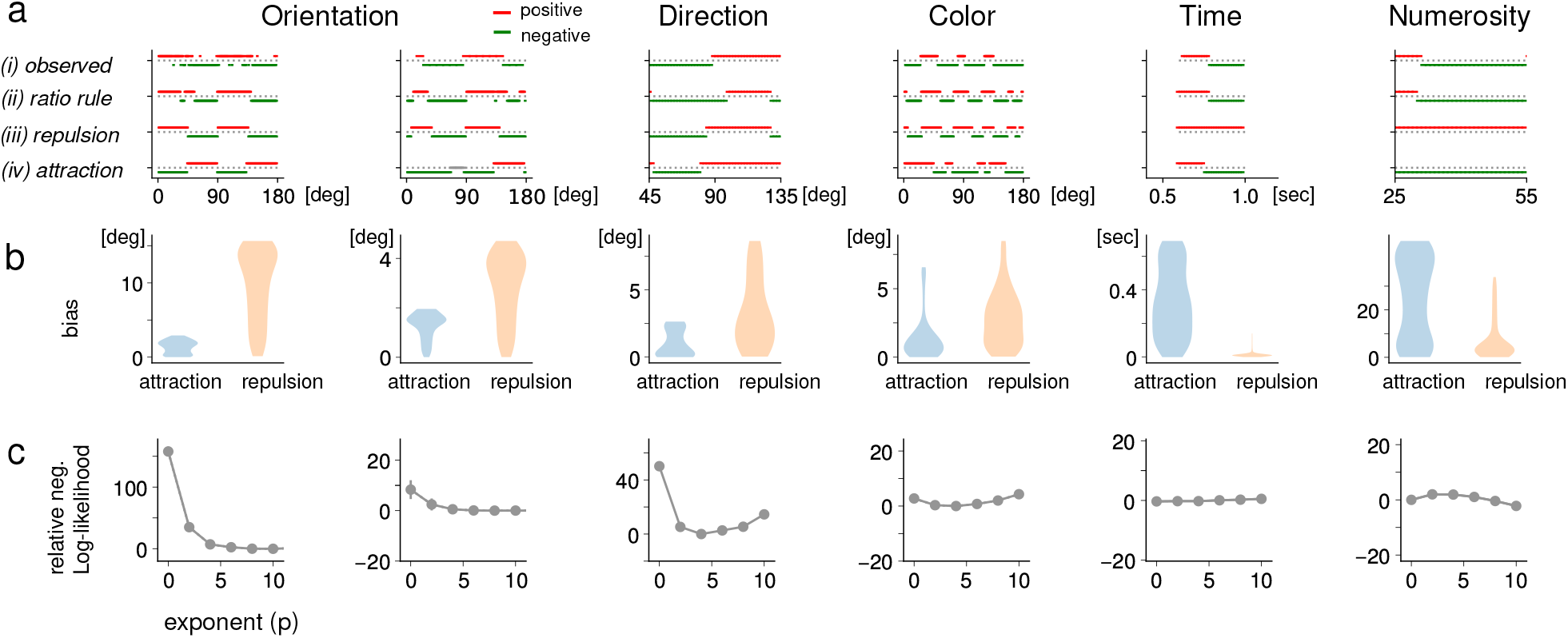
Relative role of attractive and repulsive biases, and their relation to the loss function, for the six datasets fitted in Figures 3–6. We plot data from the Cardinal group in the Direction dataset, and from one interval in the Numerosity dataset. (a) P/P Ratio Rule: For each stimulus, we plot the sign of (i) the observed bias, (ii) the prediction made by the P/P Ratio Rule, (iii) likelihood repulsion, (iv) prior attraction. The P/P Ratio Rule accurately predicts the sign of the observed bias. Repulsive biases play a larger role in accounting for the overall biases in the four circular datasets, and a smaller role in the two scalar datasets. (b) Magnitudes of likelihood repuslsion (blue) and prior attraction (orange). We show violin plots describing the distribution of absolute value bias magnitudes (lowest=0) across the stimulus space. Repulsion plays a dominant role in the four circular datasets, and only a very small role in the two scalar datasets. (c) Model fit (y-axis), measured by cross-validated negative log-likelihood (lower is better), relative to the reported model, by loss function exponent (x-axis). Error bars indicate standard errors across 10 folds, except for Direction and Magnitude, where only one fold was used. For the orientation and direction datasets, different loss functions provide substantially different model fit. For the two datasets with interval stimulus spaces, the loss function has little impact on fit, because biases are dominated by prior attraction.

## Discussion

Traditional Bayesian perspectives emphasize perceptual biases due to prior expectation (16, 84, 25, 29, 41). In contrast, recent work has discovered that Bayesian models based on efficient coding predict biases away from the prior expectation, which had been thought to be “anti-Bayesian” (35, 36). Our unifying theory closes the major gap between these divergent perspectives, and covers experimental data accounted for by neither of those prior perspectives individually. Our analytical results describe how biases are determined by the interaction among prior, encoding, loss function, and noise characteristics. A corollary is a simple and universal P/P ratio rule predicting the direction of the perceptual biases based on the scales at which prior and encoding vary over the stimulus space. We expect these results to have implications in neuroscience, psychology, and neuroeconomics (9) and decision making, where estimation of stimulus properties plays an essential role.

Through a combination of theory and analysis of experimental data across multiple modalities, our results demonstrate that perceptual biases are determined crucially not only by the prior, but also by the encoding, the stimulus range, and the loss function. When the stimulus has no boundary (circular variables) or the boundary is far away, the heterogeneity of encoding often dominates the estimation bias. Attraction to a prior, commonly considered a hallmark of Bayesian models of perception and cognition, may play a surprisingly subordinate role. For perceptual judgments within a bounded range, the boundary plays an important role. This is primarily due to large variation of the prior distribution induced by the boundary, consistent with the P/P ratio rule. Adjudicating between previous models, we found that a smoothly varying prior (e.g. 51) accounts for the datasets for time interval (63) and numerosity (66) estimation better than a sharply dropping prior (e.g. 34) exactly matching the experimental stimulus statistics.

In determining the individual elements in the Bayesian model, we have leveraged two complementary approaches: (i) constraining the component a priori using normative theory of neural coding (*e*.*g*., efficient coding) or well established behavioral results (*e*.*g*., Weber’s law) ; (ii) fitting directly to the the data. Previous work has typically assumed uniform encoding (85), simple parametric priors (29), or constraining the encoding according via efficient coding (36). Our procedure generalizes previous analysis procedures to nonparametrically fit prior and encoding. In practice, flexibly leveraging normative considerations and constraints from the data are likely to provide the most insights on the underlying factors that determine the response distribution of the observer in a given behavior task.

The current work could be extended in several ways. By considering an *L*^*p*^ family of loss function, our results generalize previous models, yet do not fully take into account the detailed shape of, possibly even asymmetric, loss functions (86, 87). Future work on larger datasets with more relaxed assumptions about the loss function may allow more precise identification of the loss function. Our results are also limited in that they only address the one-dimensional case. Recent experimental work reveals biases in the multi-dimensional case (47, 88), and generalizing our results to multi-dimensional cases is an important problem for future work. In addition, we only consider models which the decoder has full access to the encoding. For certain adaptation or contextual effects, model mis-specification (89) may have be taken into account.

One outstanding question in neuroscience is how the prior belief and encoding adapt to the statistics of the environment. Recent studies proposed that encoding and the prior should be both link to the environmental statistics, with the adaption of the prior and encoding can be formulated as statistics learning and efficient coding respectively. This seems to be a reasonable hypothesis in a stationary environment, and is supported by some experimental data (41, 36, 50). Yet how the prior and encoding dynamically evolve to reflect the changing input statistics remains an open question. One hypothesis is that the encoding may be adapted at a slower timescale than the prior (46). Our results suggest that the short-term stimulus statistics in an experiment may substantially modify the prior, and only to a lesser extent the encoding precision. For instance, in perception of motion direction, the encoding precision showed little change when observers were exposed to different motion statistics, while the prior seem to approximately reflect these short-term statistics.

Our results inform neural implementations of Bayesian inference (90, 91, 92, 93, 94, 95, 96), for which questions remain wide open. How does the neural system encode a particular prior (57, 36, 95)? How does it rapidly adjust prior belief when input is changing (97, 98, 46)? And how can it implement inference for a particular loss function in a neural circuit? Our results provide guidance for addressing these questions at the level of neural circuits. Any reasonable neural implementations of Bayesian inference should produce behavioral response patterns consistent with our theory. Furthermore, the recovered prior, encoding and loss function impose specific constrains on the neural implementation. By providing a comprehensive characterization of behavioral biases and variability, our results provide important constraints and a rigorous framework for testing a major hypothesis (*i*.*e*., *the Bayesian hypothesis*) about the computations in the brain.

## Materials and Methods

### Model Assumptions

We assume that the stimulus space 𝒳 is a contiguous one-dimensional manifold (the real line, an interval, or a circle). The sensory space 𝒴 has the same topology as the stimulus space. The sensory space conceptually represents the one-dimensional submanifold of the space of neural activations spanned by the mean encoding of the stimuli in 𝒳. We assume that *F* : 𝒳 *→* 𝒴 is bijective and differentiable, such that its slope (and thus the Fisher information of the encoding) is nowhere zero. Thus, no two stimuli are mapped to the same mean output. We assume that the prior is nowhere zero. Further, we make basic regularity assumptions: *F*′(θ), *F*^−1^(θ), and log *p*_*prior*_(θ) are twice continuously differentiable, and both they and their first and second derivatives cannot grow super-polynomially in θ. We relax these assumptions for Theorem 3 (Boundary Effects) by allowing the prior to discontinuously become zero at a boundary θ_*Max*_. These assumptions are satisfied for wide classes of models across the literature (see SI Appendix, Section S2.1.1). Under the model (1), for a given measurement *x*, the likelihood function is the distribution of *F*^−1^(*m*); it can be computed as 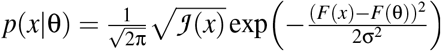.

Given an *L*^*p*^ loss function and an observed sensory encoding *m* ∈ 𝒴, the Bayes estimator 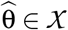 is defined as

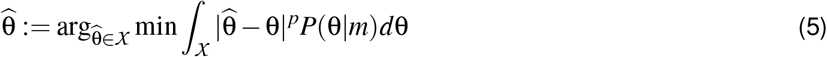

The limit *p* → 0 results in the MAP estimator. Given a true stimulus θ, the bias of an estimator 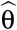 is

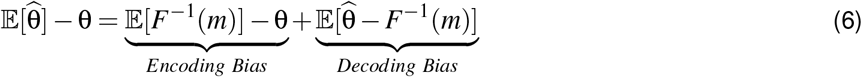

where the expectation runs over the noisy sensory encoding *m* defined in (1). Encoding bias always reflects likelihood repulsion and is independent of the loss function and the prior, whereas decoding bias includes both attractive and repulsive components depending on prior and loss function (SI Appendix, Section S3.1).

### Analytical Results

Our analytical results are stated in three theorems and a simple Prior/Precision ratio rule for judging the direction of the bias. Theorem 1 is the main theorem, addressing the biases under the model assumed in Eq. (1). Theorem 2 extends the theory to cases where both sensory and stimulus noise are present. Theorem 3 extends the theory to address biases close to a boundary. The proofs of the theorems are given in SI Appendix, Sections S3.1–S3.3. The Prior/Precision ratio rule is a straightforward consequence of Theorem 1 (see below).

### Main theory

#### Theorem 1

*Assume that* θ *is a point in the interior of the stimulus space. For the model defined in Eq. (1), the bias of the estimate* 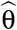 *for L*^*p*^ *loss (p* ≥ 1 *an integer) with arbitrary prior, for observed stimulus* θ, *can be written as*

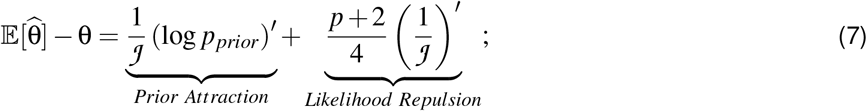

*and, for the MAP estimator (p* = 0),

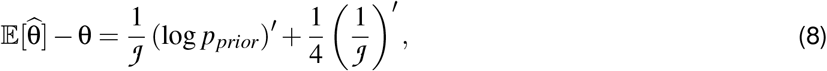

*both up to approximation error* 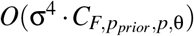, *when* σ *is sufficiently small, where the error contains constants depending on F, p*_*prior*,_, *p*, θ, *but not* σ. *Here* 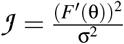 *is the Fisher information; both 𝒥 and p*_*prior*_ *are functions of* θ.

The error term 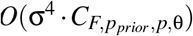 indicates that the error scales, on the one hand, with σ^4^, and on the other hand, with quantities that depend on the exponent and on the smoothness of encoding and prior. When σ < 1, *O*(σ^4^) has a smaller order of magnitude than 1*/J* ; hence, the approximation error is small relative to the size of the bias when noise is sufficiently small (se SI Appendix, Figure S4). The theorem is proven in SI Appendix, Section S3.1.

#### Prior/Precision ratio rule

To derive the Prior/Precision Ratio Rule, the bias (7) can be written as

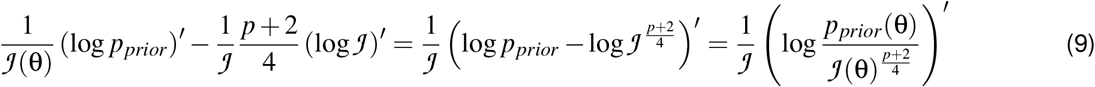

which has the same sign as *Q*(θ).

### Extensions of the theory

We extend the theory to address the impact of stimulus noise and the stimulus boundary.

#### Extension 1: Stimulus noise modulates the biases

Theorem 2 considers the impact of stimulus (external) noise, assuming that observers take both internal and external noise correctly into consideration. Formally, we extend the model (1) to

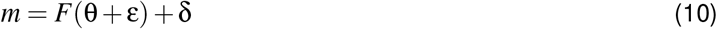

where ε is Gaussian stimulus noise with variance τ^2^.

##### Theorem 2

*Consider the model in Eq. (10). Assume that* θ *is a point in the interior of the stimulus space. Let* σ^2^ *be the variance of sensory noise and* 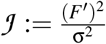 *the Fisher information of the sensory encoding. Let* τ^2^ *be the variance of stimulus noise. Then, for even integers p* > 0, *the bias of the estimate* 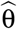 *for L*^*p*^ *loss is*

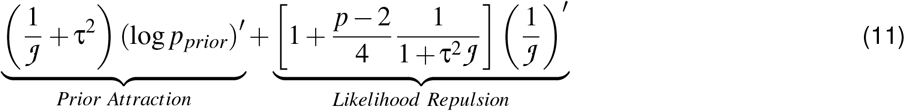

*up to approximation error of order* 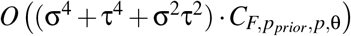 *when* σ, τ *are sufficiently small*.

The special case τ^2^ = 0 recovers the corresponding expression in Theorem 1. In the prior attraction term, stimulus noise leads to a straightforward increase in bias. In the likelihood repulsion term, stimulus noise decreases repulsion when *p* > 2. This formal result, proven in SI Appendix Section S3.2, covers even exponents *p* > 0. See SI Appendix, Figure S5 for other exponents.

#### Extension 2: Boundary-induced biases

In order to understand boundary effects, we consider the case where the prior is truncated at some point θ_*Max*_ close to the point θ, formally, *p*_*prior*_(*x*) ≡ 0 when *x* > θ_*Max*_. Let 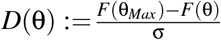 the distance of θ from the boundary in sensory space, normalized by noise SDs. Then, we have the following result, proven in SI Appendix, Section S3.3:

##### Theorem 3

*Assume the model in Eq. (1), and assume p*_*prior*_(*x*) ≡ 0 *if and only if x* > θ_*Max*_. *Assume* θ *<* θ_*Max*_. *For some positive constants C*_1,*p*,θ,*F*,σ_, *C*_2,*p,D*_, *C*_3,*p,D*_ *the bias (for even integers p >* 0*) is given as*

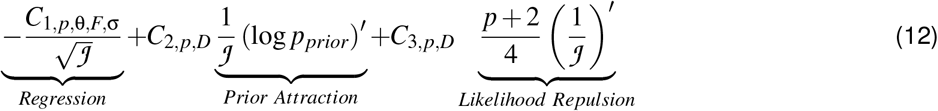

*up to approximation error of order* 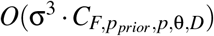 *when* σ *is sufficiently small, where the following conditions hold: C*_1,*p*,θ,*F*,σ_ = Θ(1) *as* σ → 0; lim_*D*→∞_ *C*_1,*p*,θ,*F*,σ_ = 0, lim_*D*→∞_ *C*_2*/*3,*p,D*_ = 1.

The coefficient *C*_1,…_ increases with the exponent *p* (see SI Appendix, Figure S7), so that higher exponents lead to a stronger regression effect. We note that, while the boundary-induced regression effect bears some conceptual similarity to attraction to a prior, there are two key differences: First, the boundary effect increases with higher exponents, unlike prior attraction. Second, the boundary effect scales with 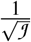, whereas prior attraction scales with 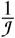 ; the former is larger when noise is small, so that the boundary effect dominates repulsion and attraction effects close to the boundary (SI Appendix, Figure S6). This correctly accounts for biases in subjective ratings on a bounded scale (SI Appendix, Section S1.2).

#### Fitting Procedure

We developed a general fitting procedure that can represent encoding and prior without restrictive parametric assumptions often made in prior work. The stimulus space is discretized as θ_1_, …, θ_*N*_; resource allocation and prior are parameterized by the values 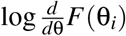 and log *p*_*prior*_(θ_*i*_), for each *i* = 1, …, *N*. Stimuli θ are mapped to the closest grid point θ_*i*_ and encoded as *F*(θ_*i*_). The encoding likelihood is computed on the discrete grid by computing the Gaussian density on each *F*(θ _*j*_) and renormalizing. An analogous approach is used for stimulus noise, when it is present. We then numerically compute the posterior by exact Bayesian inference in the discretized stimulus and sensory space. The estimator 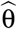 is computed using Newton’s method in the full continuous stimulus space (not just the grid), representing the integral in [5] by a sum over the discretized stimulus space.

The observed responses are assumed to result from the estimator by adding zero-mean motor noise with a stimulus-independent width. The motor distribution is represented on the full continuous stimulus space, except in datasets where the stimulus space is discrete (numerosity perception). All noise types (sensory, stimulus, motor) are Gaussian for interval stimuli and von Mises for circular stimuli. We assume that responses reflect a mixture of estimator plus motor noise and a uniform distribution, weighted with a fitted guessing rate. Motor noise impacts the variability of the observed response, but not its bias. Guessing decreases the bias by a constant factor because random guessing is unbiased; we elide this in stating our theoretical results for simplicity.

The size of the grid is 180 for circular stimuli, 200 for time interval perception, and equal to the number of discrete numbers involved (151) in numerosity perception.

Parameters are fitted to maximize trial-by-trial log-likelihood using gradient descent (see SI Appendix, Section S2.1 for details). We employ standard automated differentiation methods implemented in PyTorch (99), differentiating through [5] using the implicit function theorem (SI Appendix, Section S2.1.2). Besides resource allocation and prior, further parameters are magnitudes of sensory, stimulus, and motor noise (stimulus magnitude assumed to be zero except where explicitly manipulated), and a guessing rate. In experiments with conditions involving different magnitudes of noise, parameters other than the relevant noise magnitudes were shared across conditions.

Overfitting is mitigated by a regularization term penalizing the average squared difference between logarithmic values of FI (prior) in adjacent cells, with a weight determined for each dataset through cross-validation in preliminary simulations (SI Appendix, Section S2.1.1).

We report results where data from all subjects was pooled, except for the data in Figure 4, where we instead fitted subject-specific adjustments for prior and encoding, and freely fitted all noise parameters for each subject: Analogous to standard parametric mixed-effects models, subject-specific adjustments were subjected to a mean-zero prior favoring smooth adjustments (see SI Appendix, Section S2.1.3). We parameterized the shared components of the prior and encoding relative to the central direction (prior) or the quadrant containing the central direction (encoding), and fitted the model using a standard variational approximation to the intractable likelihood (see SI Appendix, Section S2.1.3). A parameterization with subject-specific adjustments is in principle also applicable to other datasets, though fitting by-subject adjustments is computationally more expensive. See SI Appendix, Section S45 for an application to the model in Figure 6A.

##### Implementation of Loss Function

For circular stimuli (orientation, motion direction, and color), we considered two versions of the *L*^*p*^ loss in implementing (5): The first one centers the circular coordinates [− 180^◦^, 180^◦^] at *F*^−1^(*m*), and applies the ordinary absolute distance. This makes (5) a convex optimization problem. The second one, inspired by the density of the von Mises distribution, uses the loss function ℓ(*x, y*) := (1 − cos(*x* − *y*))^*p/*2^. In the low-noise regime, both are equivalent (ℓ (*x, y*) ∼ |*x* − *y*|^*p*^); hence, our theoretical results apply equally to these and other similar loss functions. We report the implementation with the cosine-based loss as it has better quantitative model fit in orientation perception, but conclusions hold for both loss functions (see SI Appendix, Figure S10–S11). Another natural distance, the geodesic (arc-length) distance, is not everywhere differentiable, rendering it impractical in the context of gradient-based fitting procedures.

##### Evaluation and Visualization

In order to estimate model fit, we randomly partitioned the trials from each subject into 10 folds. We then conducted cross-validation (100), fitting the model for maximum likelihood on 9 folds (pooled across subjects) and recorded the held-out likelihood of the remaining fold. We report the averaged log-likelihood across the 10 folds. For the numerosity dataset (N=300) and the motion direction dataset (involving by-subject parameters), we only evaluated on one fold due to computational cost.

When both encoding and prior are nonuniform, we separately estimated the prior attraction as the decoding bias of the MAP estimator, and the likelihood repulsion as the difference between the overall bias and the MAP estimator’s decoding bias. For circular stimuli, bias and variability are reported using circular statistics.

#### Datasets

Orientation (38): 49 subjects reproduced the orientation of a Gabor patch by adjusting a blue strip. Stimuli were uniformly distributed. However, on some of trials, the stimulus had orientation 0, 45, 90, 135 degrees; we excluded these trials. On each trial, exposure time was varied among 20, 40, 80, 160, 1000 ms; no stimulus was shown on 10% of trials. We excluded the latter set of trials. In Figure 3, we omit the 20ms condition for visualization purposes as responses are too variable to be reliably estimated with Kernel smoothing; however, these data were included in model fitting.

Orientation (39): Five subjects aligned two comparison dots with the average orientation of an array of Gabor patches. Within each array, the orientations were sampled from a Gaussian distribution with an SD of 2 degrees (low stimulus noise) or 14 degrees (high stimulus noise). Exposure duration was varied between trials; results are averaged across these in Figure 3.

Perceptual learning of motion direction (67): 36 subjects used a response bar to indicate the motion direction of a cloud of points. Subjects were exposed to two distributions distinguished by colors; one distribution was bimodal, the second one was flat or trimodal for 18 subjects each. We report results including all trials associated with the bimodal distribution. Following (67), we excluded the initial 200 trials of each session. We excluded trials where no stimulus was shown (zero contrast). There were two contrast levels, modeled as separate levels of sensory noise.

Numerosity (66): 300 subjects estimated the number of dots on a screen (min 15, max 65). Within each block, all stimuli were uniformly distributed among an interval of width 30. At the beginning of each block, the prior was made salient by informing the subject about the mean number in the block. We assumed a separate prior for each interval, shared across subjects.

Time intervals (63): A total of 15 subjects estimated and reproduced time intervals (Ready-Set-Go Task) in Experiments 1 and 2. There were multiple contexts differing in whether subjects were asked to reproduce the time interval or its 0.75 or 1.5-fold; we only considered data from sessions in the identity condition.

Color (65): Seven subjects reproduced color hues by clicking on on a color wheel in CIELAB space. Stimuli were equally spaced on the wheel. Three further subjects performed the task with a delay of 900ms before the response; see SI Appendix, Figure S45.

Central tendency effect in color hue (64). Unlike the other datasets, this used a 2AFC design; the model was not fitted to trial-by-trial data. We included results reported for Experiments 2 and 3; red corresponds to the average of the identical conditions 2s in Experiment 2 and None in Experiment 3; blue corresponds to 4s in Experiment 2; green corresponds to High in Experiment 3. Following some previous work (e.g. 29, 41), we assumed that subjects first decode 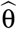 and then compare it to the reference, as the basis for the response in the 2AFC task.

## Acknowledgement

This research uses data from a number of previous published studies. We would like to thank Gi-Yeul Bae, Vincent de Gardelle, Nikos Gekas, Joshua Solomon, Rafael Polania, and Christian Ruff for sharing their data with us, as well as the authors of several other studies for making their data publicly available. We thank Alan Stocker and Michael Woodford for helpful discussions.

