## Supplementary material for "A unifying theory explains seemingly contradicting biases in perceptual estimation": SI Appendix

Michael Hahn
Xue-Xin Wei

December 9, 2022

### S1 Simulations

#### S1.1 Illustrating Theorems 1–3

In this section, we provide simulations illustrating the implications of Theorems 1–3 (see Main Text, Equations (2–3) and theorems in Materials and Methods).

1. *(Additive Decomposition)* Theorem 1 describes an additive decomposition of biases into attraction and repulsion; repulsion but not attraction depends on the loss function. We illustrate this in Figure S1, where we show the bias for all combinations of a set of encoding resource allocations with a set of priors.

2. *(Sensory Noise and Loss Function)* Theorem 1 implies that sensory noise increases both repulsion and attraction. It further states that repulsion – but not attraction – scales linearly (factor $\frac{p+2}{2}$) with the exponent $p \geq 1$. This functional form does not hold for $p = 0$, where the factor is $\frac{1}{4}$ instead of expected $\frac{1}{2}$. Figure S2 exemplifies this at the example of orientation estimation. Figure S3 shows the scaling of bias with sensory noise and with loss function exponents, including non-integer exponents. Results transfer to non-integer exponents; furthermore, there is a nonlinearity close to 0, corresponding to the difference in functional form.

3. *(Behavior at Large Sensory Noise)* Theorem 1 describes the bias in the limit when noise is small. In Figure S4, we examine the behavior of the bias as sensory noise increases, for the same point as selected in Figure S3. For small noise, biases scale quadratically with $\sigma$ as predicted by the analytical approximation. For large noise, biases grow more slowly and saturate, reflecting averaging out of local variation.

4. *(Role of Stimulus Noise)* While sensory noise increases both components of bias according to Theorem 1, Theorem 2 shows that the action of stimulus noise is different for attractive and repulsive components: Stimulus noise always increases prior attraction. It leaves likelihood repulsion unchanged when $p = 2$, but decreases it when $p > 2$. These analytical results are verified by the simulation in Figure S5.

5. *(Boundary Effects)* Theorem 3 describes how abrupt truncation of the prior leads to both a regression effect into the interior, and modulation of attractive and repulsive components close to the boundary. This is illustrated by simulations in Figure S6. Whereas prior attraction is independent of the loss function, the boundary effect increases with the loss function exponent (Figure S7).

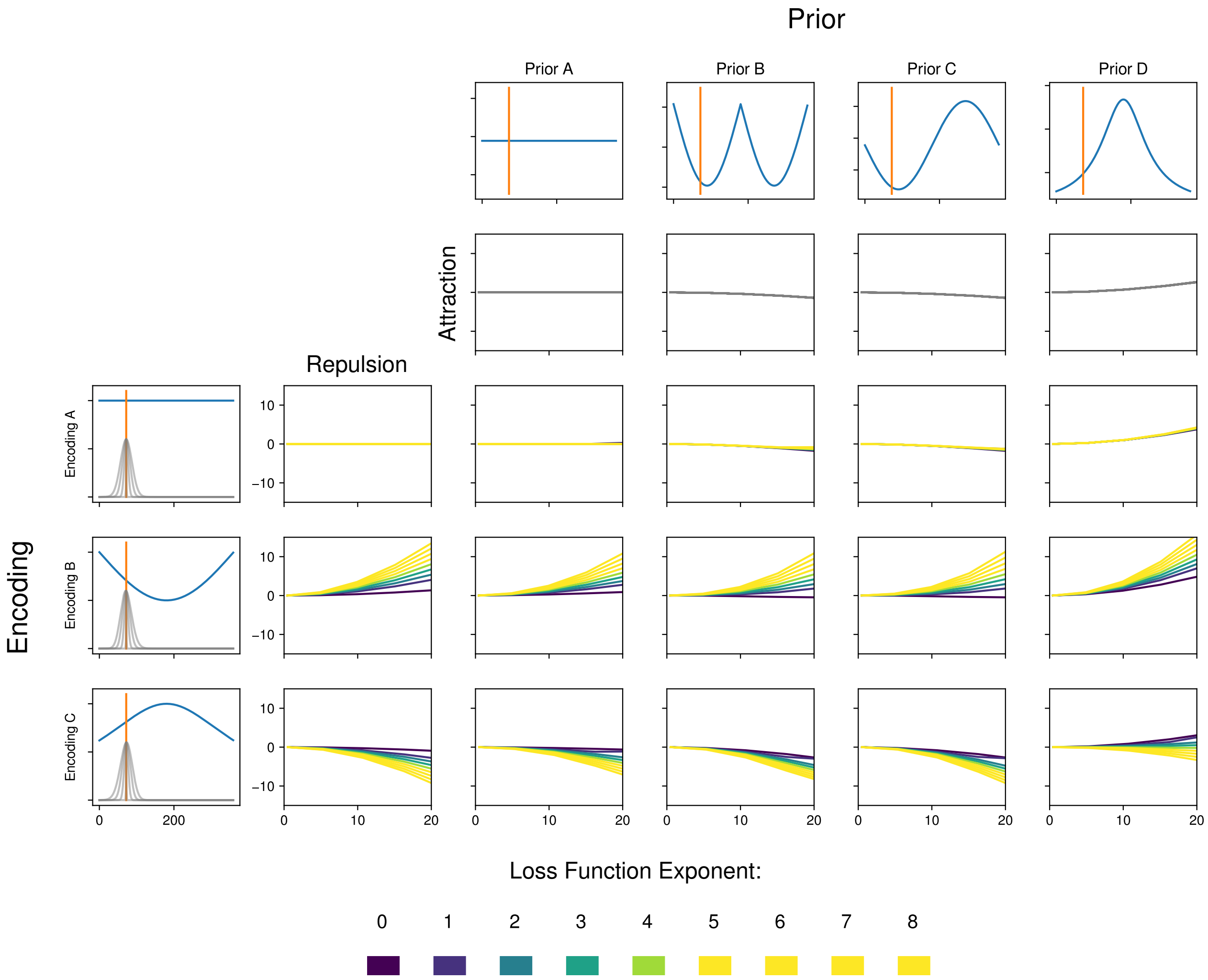

Figure S1: Additive effects of prior attraction and likelihood repulsion on biases at a stimulus θ, as a function of encoding resource allocation, prior, and sensory SD (quantified as the SD of the likelihood). All biases are numerically simulated. **Rows** represent three different sample encoding resource allocations ($\sqrt{\mathcal{S}} = F'(\theta)$ is plotted in the left columns): one is uniform, one is bimodal, one is unimodal. The position of the stimulus θ in stimulus space is indicated by the orange line. In order to make biases comparable across the three encoding resource allocations, we chose θ so that $\mathcal{S}(\theta)$ is approximately constant across the three encodings. For each encoding, we plot the corresponding likelihood at the different levels of sensory noise. The resource allocations give rise to different likelihood repulsion biases at θ, plotted in the second column, as a function of sensory SD σ (expressed as the SD of the likelihood projected into the stimulus space $\mathcal{X}$). **Columns** indicate four different priors (plotted in the first row), with corresponding prior attraction biases at θ (plotted in the second row, again as a function of likelihood SD). The remaining cells indicate the simulated overall bias determined by encoding and prior. As predicted by Theorem 1, each bias arises as the sum of the attractive and repulsive biases corresponding to its row and column. Prior attraction is independent of the loss function, whereas the strength of likelihood repulsion increases with $p$.

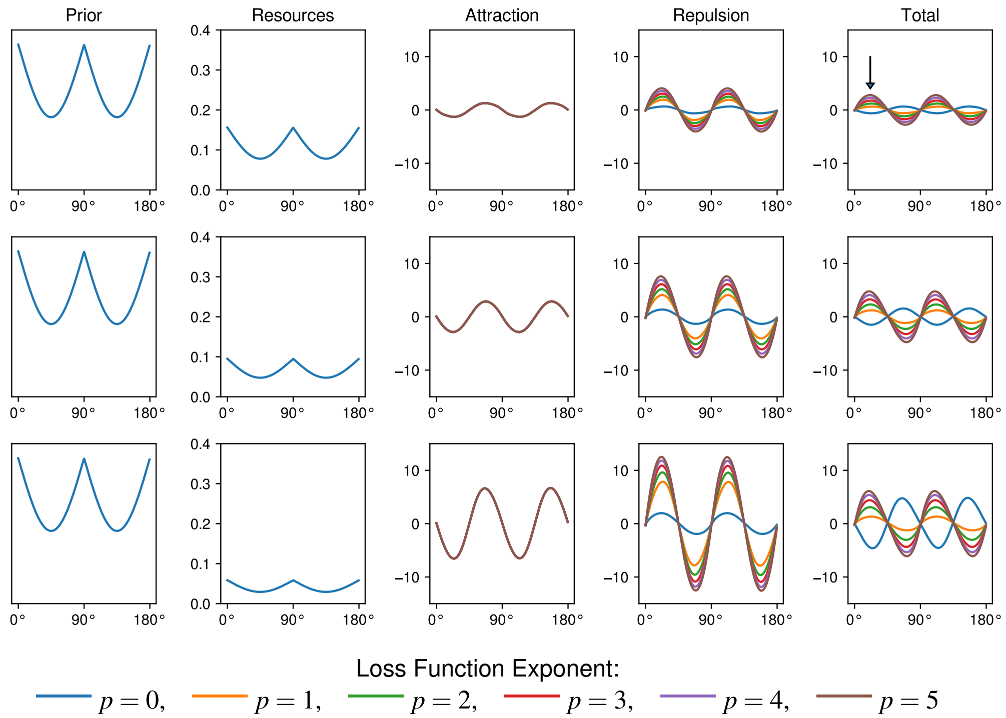

Figure S2: Role of sensory noise magnitude and loss function exponent, at the example of orientation estimation. For a model of orientation perception with matched prior and encoding precision (compare Main Text, Figure 1D 5–7), we show simulated biases for three levels of sensory noise (rows) and loss function exponents from 0 to 5 (colors). As predicted by Theorem 1, sensory noise increases attractive and repulsive components. Loss function exponent increases repulsive components, but not prior attraction. In this case, the overall bias is attractive when $p = 0$ and repulsive when $p \geq 1$. The arrow refers to the stimulus analyzed further in Figures S3.

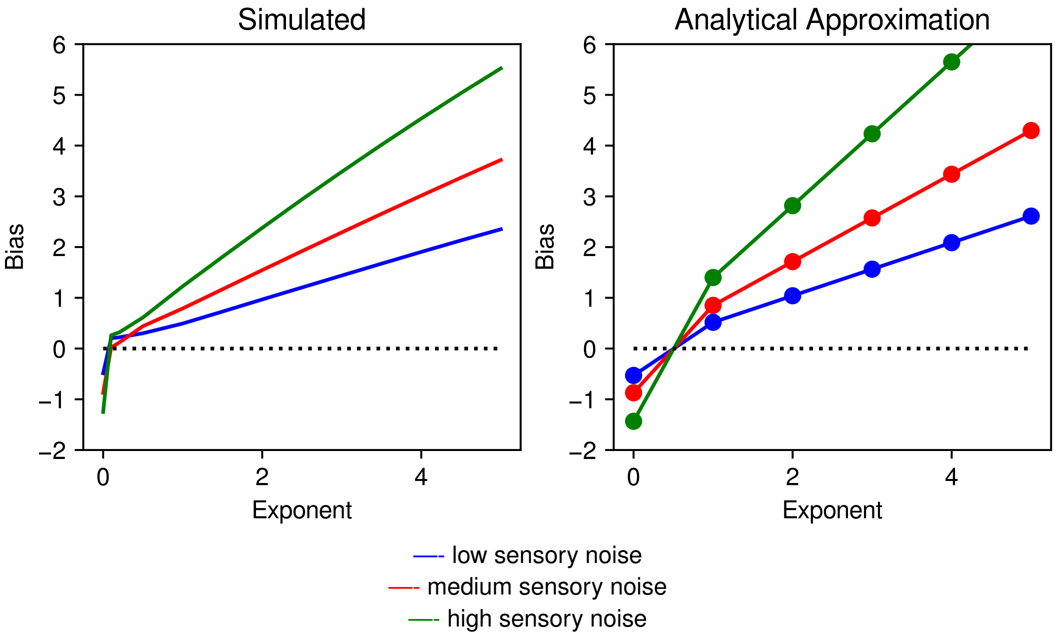

Figure S3: Role of sensory noise magnitude and loss function exponent, at the example of orientation estimation. Bias (at the 22.5° peak of the repulsive bias in orientation estimation, indicated by the arrows in Figure S2) as a function of the loss function exponent $P$, for three levels of sensory noise. On the left, we show simulated bias including fractional exponents. On the right, we show analytical approximations for integer exponents. As predicted by Theorem 1, the bias is a monotonic function of the exponent, and approximately linear when noise and exponent are small; for larger noise and exponent, the finite size of the stimulus space leads to sublinear growth (as seen in the green curve on the left). The monotonic relationship between exponent and bias, shown for integer exponents in Theorem 1, smoothly extends to fractional exponents. There is a very rapid jump as $P \to 0$; in the analytical expression, this is reflected in a special expression for the MAP estimator's bias (Equation [3] in main paper). The bias is attractive when $P$ is 0 or very close to 0, and repulsive otherwise.

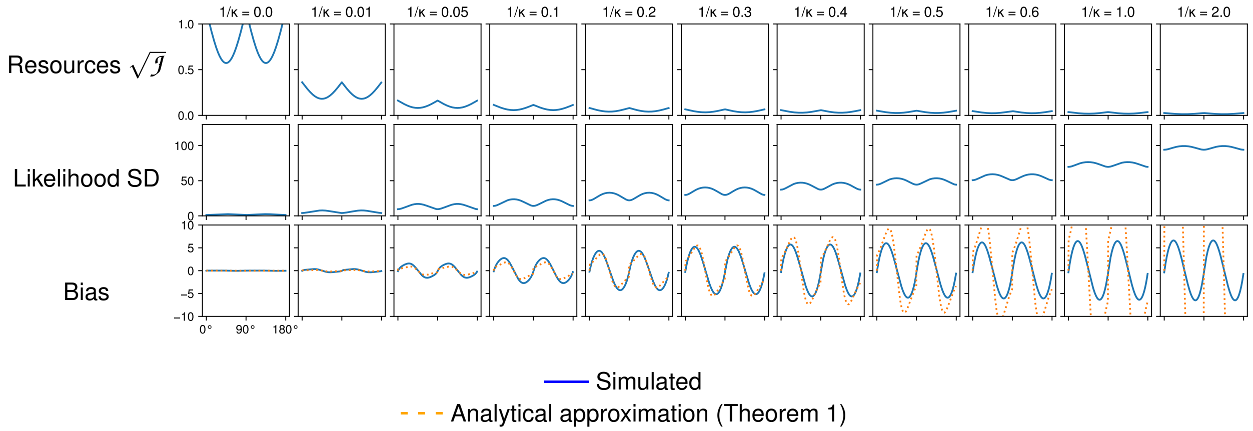

Figure S4: Scaling of bias in the example of Figure S2 as a function of the sensory noise magnitude $\sigma$ at $p = 2$. We describe noise magnitudes in terms of $\frac{1}{\kappa}$ ($\kappa$ the parameter of the von Mises distribution), which corresponds to the variance of the likelihood in $\mathcal{Y} = [0, 2\pi]$ in the low-noise limit. **First row:** $\sqrt{\mathcal{J}}$. **Second row:** SD of the likelihood transformed back into the stimulus space $\mathcal{X} = [0^\circ, 180^\circ]$. **Third row:** Bias of the estimator at $p = 2$, simulated (solid) and predicted by Theorem 1 (dotted). For small noise, biases scale quadratically with the noise magnitude, as $\frac{1}{\mathcal{J}} = \frac{\sigma^2}{\mathcal{S}} \propto \sigma^2$. For larger noise magnitude, biases grow more slowly and saturate. Hence, when noise is very large (likelihood SD $\gtrapprox 40^\circ$), biases are smaller than predicted by Theorem 1, though with the same qualitative behavior.

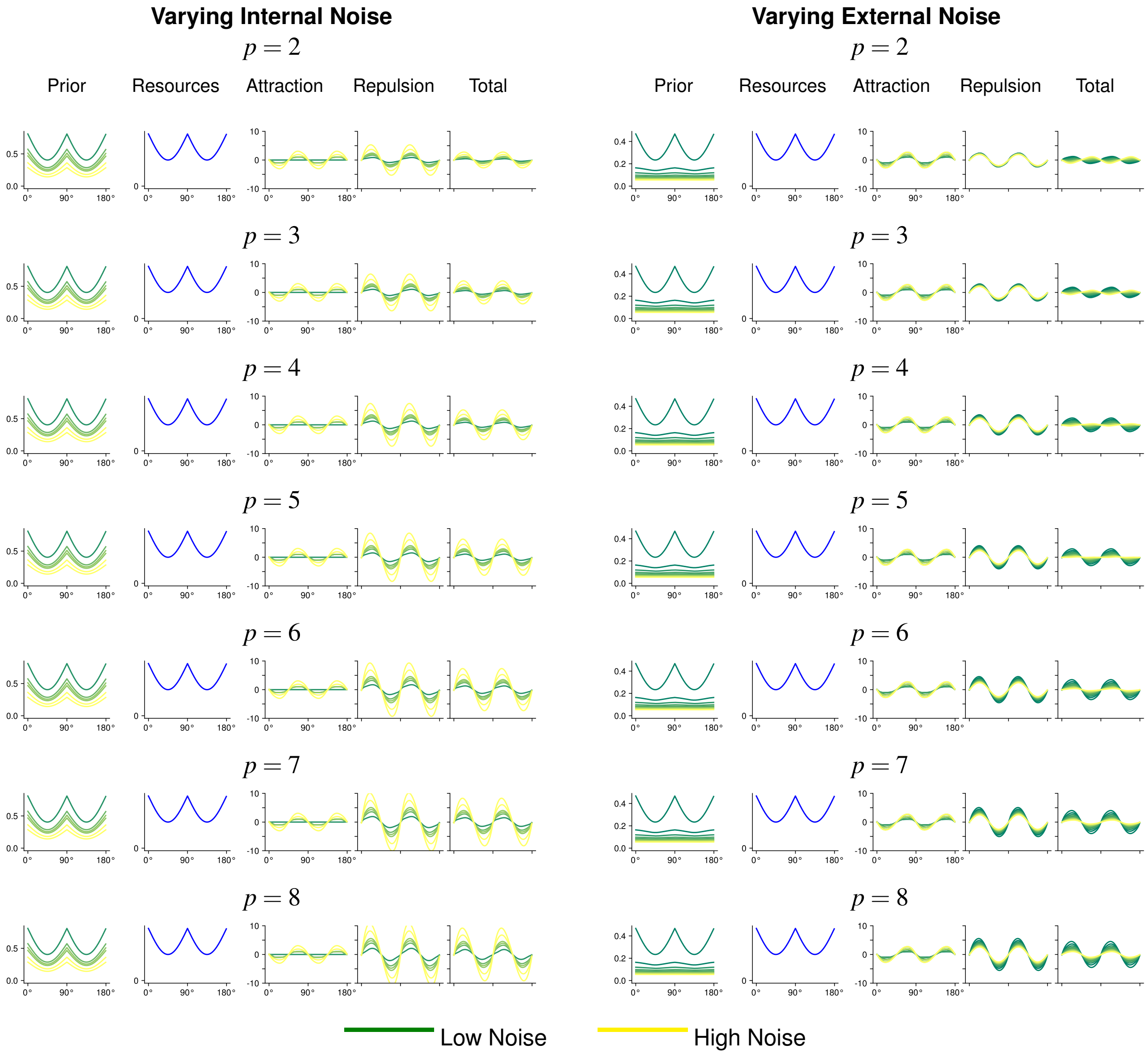

Figure S5: Effect of internal and external noise on bias. Simulated bias in orientation perception, assuming prior and encoding matched to natural image statistics. **Left:** Varying sensory noise at zero stimulus noise. Increasing internal noise (yellow) increases both attraction and repulsion, independently of the loss function, as predicted by Theorems 1 and 2. **Right:** Varying stimulus noise while keeping sensory noise fixed. Increasing stimulus noise (yellow) increases prior attraction independently of the loss function. While it leaves likelihood repulsion approximately unchanged at $p = 2$, it decreases the repulsive component for high exponents, as predicted by Theorem 2.

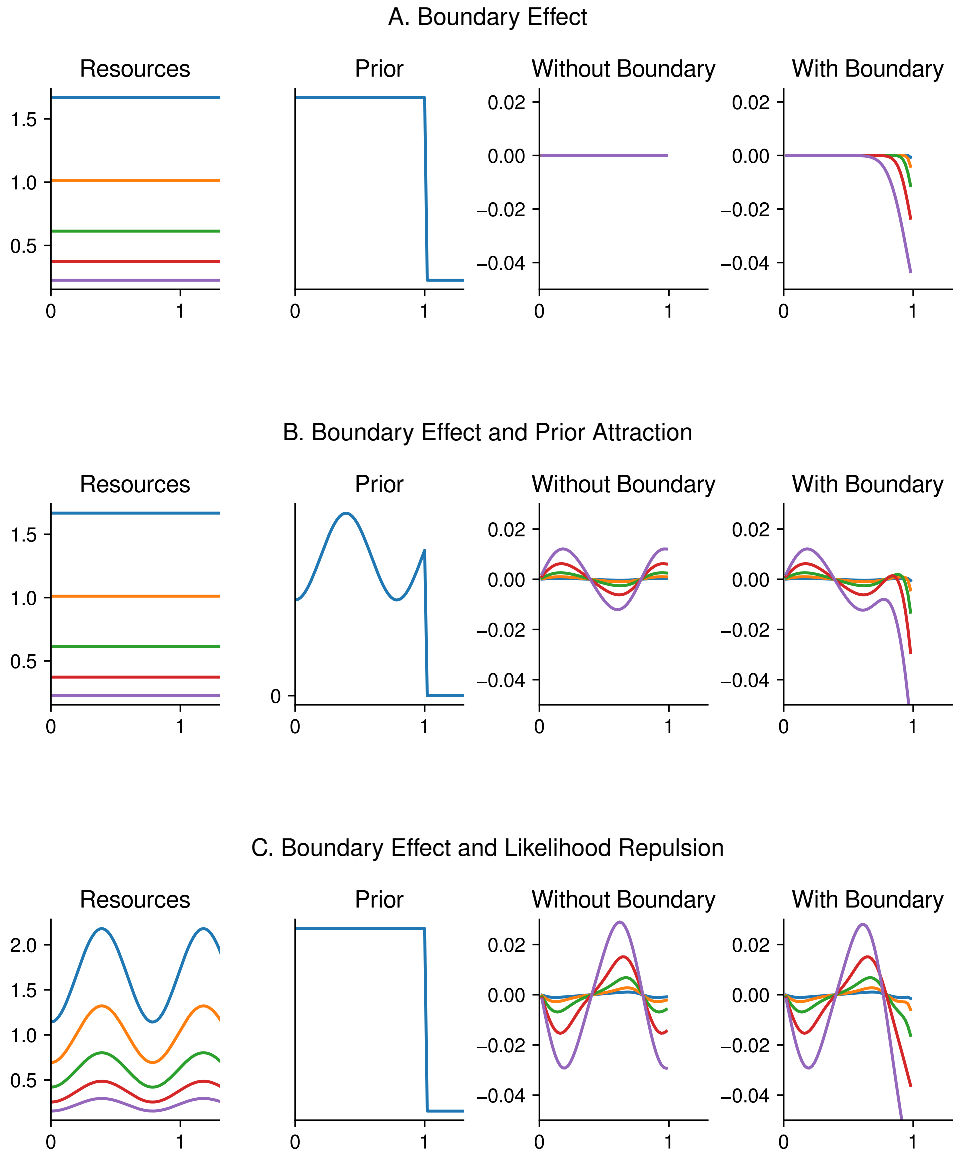

Figure S6: Boundary effects (simulated, at $p = 2$): Within a large stimulus space $\mathcal{X} = [-10, 10]$ with uniform or periodic prior and resource allocation (here, only showing $[0, 1.2]$), we consider the effect of truncating the prior at $\theta_{Max} = 1$: A. In the vicinity of the boundary, a regression effect into the interior is observed. As sensory noise increases, both the magnitude of the bias and the area affected increase. B. Truncating a nontrivial prior at the boundary leads to a combination of prior attraction in the interior and a regression effect close to the boundary; the regression effect prevails close to the boundary. C. The same happens to likelihood repulsion.

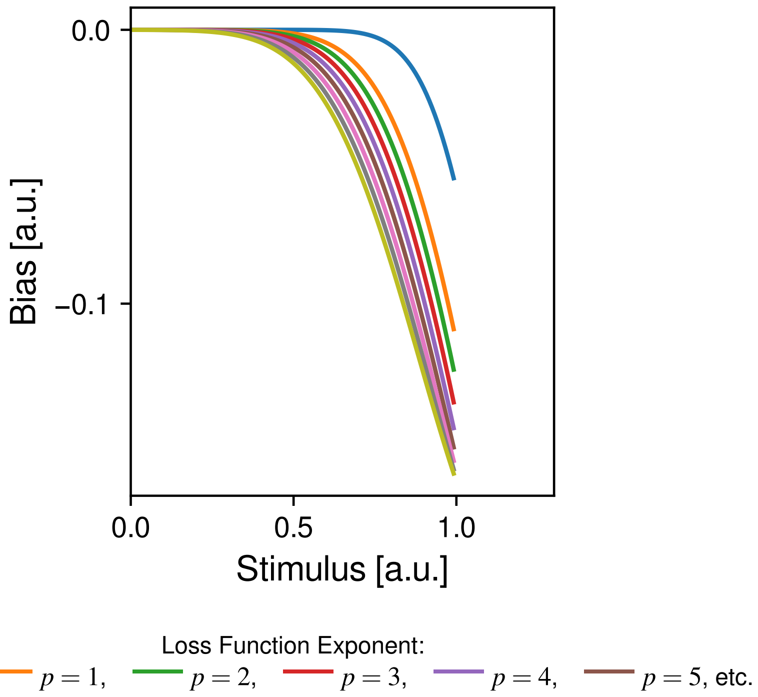

Figure S7: Effect of loss function on the boundary effect. We show the bias resulting at the boundary in Figure S6A as a function of the exponent. In the analytical expressions, this quantity corresponds to Equation (110); it reflects a local average of the coefficient $H_{1,p,D}$ plotted in Figure S46 for positive even exponents. As predicted theoretically, the magnitude of bias increases with the loss function exponent. This explains simulation results in Jazayeri and Shadlen [15, Figure 5], where the boundary effect was much more pronounced at $p = 2$ (their *Bayesian Least Squares* [BLS]) than at $p = 0$ (MAP). This dependence on the loss function is caused by the abrupt discontinuous truncation of the prior, qualitatively different from ordinary prior attraction to a smooth prior (as described by Theorem 1), which is approximately independent of the loss function. Jazayeri and Shadlen [15] concluded that $p = 2$ provided a better model of their data; however, our results show that this task is better accounted for by a smooth unimodal prior, with fit largely independent of the loss function (Section S2.3.3.3).

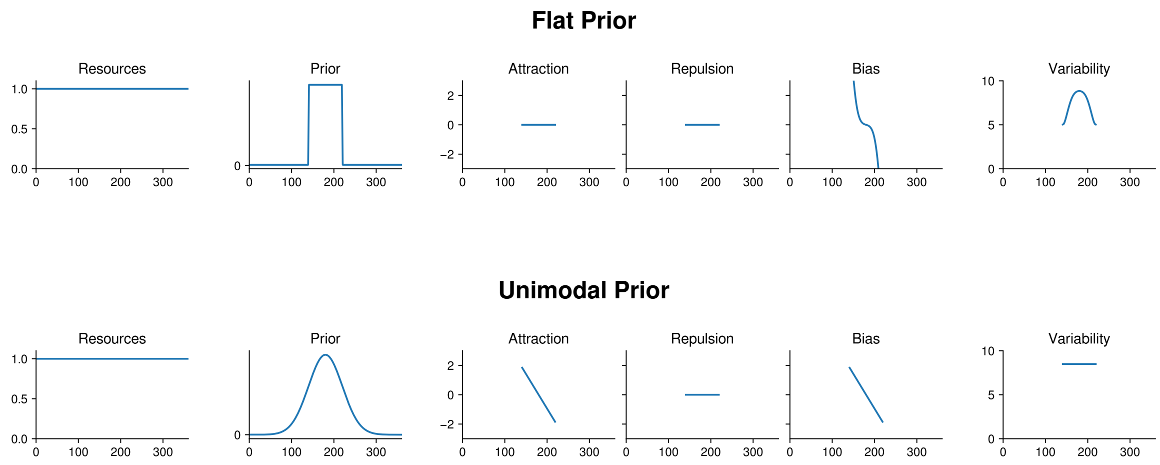

Figure S8: Comparison between flat and unimodal prior. Both kinds of prior can give rise to central tendency effect, either as a boundary effect (flat prior, [e.g. 15]) or prior attraction (unimodal prior, [e.g. 21]). An important difference is visible in the variability: The unimodal prior may not impact the variability; the flat prior leads to reduced variability at the boundary.

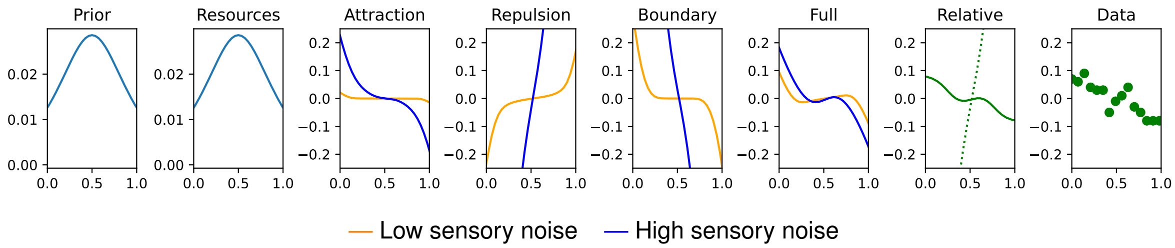

Figure S9: Bias in a bounded scale of subjective value ratings: Polanía et al. [22] measured the relative bias between short and long exposure times. The prior and resource allocation were parameterized following Polanía et al. [22]. Numerical parameters were fitted to match reported summary statistics. In the "Relative" facet, the dotted curve indicates the corresponding prediction without boundary effects. The predicted relative bias (solid) shows a repulsive bias away from the center when close to the center, and a bias towards it further out. This bias towards the center cannot be explained as prior attraction, because attraction would have to also dominate in the center if it dominated the repulsion away from it. Rather, the pattern is correctly accounted for in terms of the regression away from the boundary, similar to Figure S6C: Likelihood repulsion accounts for biases in the interior, whereas the regression effect dominates closer to the boundary. The dotted line indicates relative bias in the absence of a boundary; this would be repulsive everywhere.

#### S1.2 Distinguishing Prior Attraction from Boundary Effects

Here, we consider an example where regression to the mean reflects boundary effects, rather than attraction to a smooth prior. Polanía et al. [22] measured the relative bias in subjective value judgments on a bounded scale when sensory noise was high or low, finding an outwards bias around the center, and an inwards bias elsewhere. While directly fitting the encoding model on this data is infeasible as the ground truth values (i.e., subjects' subjective values) were not measured, we simulated our model based on the encoding and prior determined in independent experiments by Polanía et al. [22]. Results are shown in Figure S9. Importantly, even though the prior peaks in the middle of the range, the regression effect cannot be explained as prior attraction here: If prior attraction were responsible for the regression to the mean, biases should be attractive towards the center throughout the entire range. The fact that the regression effect only appears along the boundary shows that boundary effects are instead responsible. While prior attraction shows a broadly similar qualitative pattern as regression to the mean, the simultaneous apperance of attraction on the boundary and repulsion in the center is absent when disregarding the boundary effect.

### S2 Model Implementation and Results

#### S2.1 Fitting Procedure

##### S2.1.1 Model Specification and Implementation

**Model Specification** Here, we recapitulate all elements of the model for reference. The parameter set $\Theta$ of the model is given by:

1. The stimulus space, which is an interval or the circle.
2. The sensory space, which has the same topology as the stimulus space. Without loss of generality, its length can be assumed to be 1 (for intervals) or $2\pi$ (for the circle).[1]
3. $p > 0$, the loss function exponent
4. $p_{prior}(\theta)$, a function on the stimulus space
5. $F(\theta)$, a function mapping from the stimulus space to the sensory space.
6. $\sigma^2$, the variance of sensory (internal) noise
7. $\tau^2$, the variance of stimulus (external) noise
8. $\rho^2$, the variance of motor noise
9. $\kappa$, the guessing rate

When noise is von Mises (for circular stimulus spaces), $\sigma^2, \tau^2, \rho^2$ are identified as the reciprocal of the "inverse temperature" parameter $\kappa$ of the von Mises distribution; this equals the variance in the small-noise regime.

When there are multiple noise conditions (e.g., different exposure durations) within an experiment, there correspondingly are multiple noise parameters corresponding to the conditions (e.g., one value of $\sigma^2$ per exposure duration), while all other parameters are kept fixed. Details for each dataset are provided in Section S2.3.

If $\Theta$ is the full parameter set of the model, then the generative model generating a response $\theta'$ given a stimulus $\theta$ is as follows:

1. Stochastic encoding:
$$m = F(\theta + \varepsilon) + \delta \tag{1}$$
where $\varepsilon \sim N(0, \tau^2)$, $\delta \sim N(0, \sigma^2)$.
2. Bayes estimator:
$$\widehat{\theta}_{[m,\Theta]} := \arg_{\widehat{\theta} \in X} \min \int |\widehat{\theta} - \theta|^p P(\theta|m) d\theta \tag{2}$$
where the posterior $P(\theta|m)$ is determined by the model parameters $\Theta$, specifically the encoding likelihood and the prior.
3. Response: With probability $\kappa$, the response is uniform on the stimulus space; with probability $(1-\kappa)$ it is $\widehat{\theta}_\ell + \varepsilon_{Motor}$ where $\varepsilon_{Motor} \sim N(0, \rho^2)$ (or analogously von Mises, if the space is circular).

Given model parameters $\Theta$, the log-likelihood of a single trial with response $\theta'$ and true stimulus $\theta$ is thus given as

$$\log p(\theta'|\theta;\Theta) = \log \left\{ \frac{\kappa}{\theta_{Max} - \theta_{Min}} + (1-\kappa) \cdot \int\int P_{Normal}(\theta'|\widehat{\theta}_{[m=F(\theta+\varepsilon)+\delta,\Theta]}, \rho^2) P_{Normal}(\varepsilon|0,\tau^2) P_{Normal}(\delta|0,\sigma^2) d\varepsilon d\delta \right\} \tag{3}$$

[1] Rescaling it simply results in rescaling of $F$ and $\sigma$. This has no effect on $\mathcal{J}$, or any other observed quantities.

The overall negative log-likelihood is the sum over all datapoints $(\theta_{True,i}, \theta'_i)$ in the dataset:

$$-\sum_{i=1}^{K} \log p(\theta'_i|\theta_{True,i};\Theta) \tag{4}$$

This definition of the likelihood pools data from all subjects, irrespective of systematic differences between subjects. The model can also be fitted with subject-specific effects to account for such subject differences; see Section S2.1.3 for this version.

**Model Implementation** The encoding and prior are encoded by assigining one value of $F'$, $p_{prior}$ to each cell in the discretized grid. Internally, in order to ensure nonnegativity and normalization, these are parameterized using the softmax transformation:

$$p_{prior}(\theta) = \frac{\exp(\beta(\theta))}{\sum_{\theta'} \exp(\beta(\theta'))} \tag{5}$$

for $\theta$ on the grid, and similarly for $F'(\theta)$; here, $\beta(\theta_i)$ ($i = 1, \ldots, N$) are free real-valued parameters of the model. The transfer function is given as

$$F(\theta) = \int_{\theta_{Min}}^{\theta} F'(\theta')d\theta' \tag{6}$$

where $\theta_{Min}$ is the lower end of the stimulus space; it is implemented as the cumulative sum of the discretized $F'$. [2]

The input stimulus $\theta$ is mapped to the closest element of the discretized grid, and is then encoded as the corresponding entry of the discretized $F$. The likelihood is projected onto the discretized grid; sensory noise results in another point on the grid. The posterior is computed on the discrete grid using exact Bayesian inference. The estimator $\widehat{\theta}$ is computed by loss function minimization over the full continuous range of the stimulus space, not just the discretized grid. For this, we use Newton's method.[3]

In keeping with the parameterization of encoding and prior, the sensory noise variance is encoded as a fraction of the size of the sensory space using the inverse-logit transform:

$$\sigma^2 = \frac{1}{1 + \exp(-\gamma)} \tag{7}$$

where $\gamma$ is again a free real-valued parameter. The same parameterization is applied for the guessing rate $\kappa \in [0, 1]$. We parameterize stimulus and motor noise variance using their logarithms.

In order to mitigate and prevent overfitting, we added a regularization term to (4):

$$\frac{\lambda}{N} \sum_{i,j \text{ neighbors}} (\alpha(\theta_i) - \alpha(\theta_j))^2 \tag{8}$$

and analogously for $\beta$, where $i$ and $j$ are neighbors if $j = i + 1$ or (in the circular case), $i = N$, $j = 1$. The weight $\lambda$ was determined for each dataset using cross-validation in preliminary experiments.

We report biases using the mean of the estimate $\widehat{\theta}$ across the encodings $m$ on the discretized grid, taking the ordinary mean for interval stimuli and the circular mean for circular stimuli. The reported variability is $\sqrt{\mathrm{SD}(\widehat{\theta})^2 + \rho^2}$, where again $\mathrm{SD}(\widehat{\theta})$ is computed using circular statistics when the stimulus space is circular[4].

[2]When $\mathcal{X}$ – and equivalently $\mathcal{Y}$ – is circular, say $[0, 2\pi]$, $\theta_{Min}$ can be taken as 0, and the output $F(\theta) \in [0, 2\pi]$ is interpreted as a point on the circle.

[3]When $p > 1$, we apply Newton's method to minimize the $L^p$ loss; for $p = 0$, we use it to maximize the posterior. When the second derivative in loss minimization (similarly for posterior maximization) has the wrong sign in some iteration (negative in the case of minimization), we take a gradient descent step rather than a Newton step; this avoids divergence or convergence to a *maximum* of the loss when the loss is not convex.

[4]The circular SD is given as $\sqrt{-2\ln(\overline{R})}$, where $\overline{R}$ is the Eucliden distance between the origin and the vector mean of the angles.

###### S2.1.2 Fitting Procedure

**Parameter Fitting** We fitted the model separately for each loss function exponent $p$. We compute derivatives of the log-likelihood (3) w.r.t. the model parameters using automated differentiation as implemented in PyTorch [20] and apply gradient descent to optimize the model parameters. On circular data, we found the optimization to be robust to different learning rate scheduling schemes; we typically used vanilla GD with learning rate decayed by 70% whenever loss did not increase over 500 steps. For interval data, we found the optimization scheme to matter at the boundary, where fewer observations are available. We found that Sign-GD [e.g. 2] with momentum provided rapid convergence compared to other common schemes. Simplex-based (e.g., Nelder-Mead) or Quasi-Newton (e.g., BFGS) methods did not offer improved convergence compared to simple gradient descent schemes.

**Gradient Computation for $L^p$ Estimator ($p > 0$)** The only component of the generative model (1–3) that is not straightforwardly differentiable using standard automated differentiation methods is the estimator $\widehat{\theta}$, which is a function of the posterior $P$ (and thus of the model parameters $\Theta$). Differentiating through Newton's method is inefficient and prone to accumulation of numerical errors; we thus chose a different route. Writing the loss as $L(\theta, P_1, \ldots, P_N)$ and $M := \partial_\theta L$, (where $N$ is the number of discretization points), the estimator's gradient w.r.t. the posterior is, using the implicit function theorem [26]:

$$\frac{\partial\widehat{\theta}}{\partial P_i} = -\left(\frac{\partial M}{\partial\theta}\right)^{-1} \cdot \frac{\partial M}{\partial P_i} \tag{9}$$

evaluated at $\theta = \widehat{\theta}$. For the ordinary distance $|x-y|^p$ ($p > 0$), we thus have [5]

$$\frac{\partial\widehat{\theta}}{\partial P_i} = -\frac{\operatorname{sign}(\theta - x_i)|\theta - x_i|^{p-1}}{(p-1)\cdot\sum_{i=1}^N P_i|\theta - x_i|^{p-2}} \tag{10}$$

This is fully applicable when $p > 1$; for $0 < p < 1$, it is applicable whenever $\theta$ does not equal any of the grid points. For the cosine loss $\ell(x,y) = (1-\cos(x-y))^{p/2}$, the result is[6]

$$\frac{\partial\widehat{\theta}}{\partial P_i} = -\frac{(1-\cos(\theta - x_i))^{\frac{p}{2}-1}\sin(\theta - x_i)}{\sum_{j=1}^N P_j\left[\frac{p-2}{2}(1-\cos(\theta - x_j))^{\frac{p}{2}-2}\sin(\theta - x_j)^2 + (1-\cos(\theta - x_j))^{\frac{p}{2}-1}\cos(\theta - x_j)\right]} \tag{13}$$

which is applicable for $p = 2, 4, 5, 6, \ldots$.[7]

**Gradient Computation for MAP Estimator** On a discrete grid, the MAP estimator is always a grid point and thus piecewise constant (hence, its gradient is zero or undefined) as a functional of the posterior. In order to obtain meaningful gradients for the MAP estimator, it is thus necessary to consider the continuous limit. In general, for an

[5] $M = \partial_\theta L = p\cdot\sum_{i=1}^N P_i \operatorname{sign}(\theta - x_i)|\theta - x_i|^{p-1}$; hence, $\frac{\partial M}{\partial\theta} = p\cdot(p-1)\cdot\sum_{i=1}^N P_i|\theta - x_i|^{p-2}$ and $\frac{\partial M}{\partial P_i} = p\cdot\operatorname{sign}(\theta - x_i)|\theta - x_i|^{p-1}$.

[6] Note

$$\frac{d}{dx}\ell(x,y) = \frac{p}{2}(1-\cos(x-y))^{\frac{p}{2}-1}\sin(x-y) \tag{11}$$

and

$$\frac{d^2}{dx^2}\ell(x,y) = \frac{p}{2}\frac{p-2}{2}(1-\cos(x-y))^{\frac{p}{2}-2}\sin(x-y)^2 + \frac{p}{2}(1-\cos(x-y))^{\frac{p}{2}-1}\cos(x-y) \tag{12}$$

[7] At $p = 3$, the denominator is unbounded when $\theta = x_j$, and the estimator is not differentiable. The denominator is finite when $p \geq 4$. At $p = 2$, the quantity simplifies to

$$\frac{\partial\widehat{\theta}}{\partial P_i} = -\frac{\sin(\theta - x_i)}{\sum_{j=1}^N \cos(\theta - x_j)}. \tag{14}$$

arbitrary smooth posterior $P$, the functional derivative of the MAP estimator is the tempered distribution[8]

$$\frac{\delta\widehat{\theta}_{MAP}}{\delta P} = \frac{\delta'(x-\widehat{\theta})}{P''(\widehat{\theta})} \tag{15}$$

where $\frac{\delta}{\delta P}$ on the left-hand-side indicates the functional derivative, and $\delta'$ on the right-hand side is the distributional derivative of the Dirac distribution. In order to evaluate this on the discretized grid, we interpolate $P_1,\dots,P_N$ using a kernel: $f(P,\theta) := \sum_{i=1}^N g_i(\theta)P_i$ where $g_i \geq 0$, $\sum_i g_i(x) = 1$. Then by (15)[9] or by the implicit function theorem applied to $\frac{\mathrm{d}}{\mathrm{d}\theta}f(P,\theta) = 0$, the gradient $\frac{\partial\widehat{\theta}_{MAP}}{\partial P_i}$ is

$$\frac{d\widehat{\theta}_{MAP}}{dP_i} = -\frac{g_i'(\widehat{\theta})}{\sum_{i=1}^N g_i''(\widehat{\theta})P_i} \tag{16}$$

We chose the RBF kernel[10] and further added a smooth approximation of the function $\log 1_{\theta_{Min}\leq x\leq\theta_{Max}}$ to smoothly restrict $f$ to the stimulus space in those datasets where $\mathcal{X}$ has a boundary[11]. We use this smoothed version $f(\theta,P)$ both for computing the MAP estimator (using Newton's method) and its gradients (16).

##### S2.1.3 Parameterization with Subject-Specific Adjustments

As described in the main text, the motion direction dataset [11] has different experimental stimulus distributions for each subject, which may lead to subject-specific adaptation of encoding and prior. We thus analyzed that dataset using subject-specific adjustments to encoding and prior. Here, we describe the appropriate version of our fitting procedure. In standard parametric mixed-effects models extending linear regression [3], intercepts and slopes receive per-subject adjustments that are regularized towards zero using a Gaussian shrinkage prior. In our setting, adjustments to encoding and prior will be functions rather than individual numbers: As described in Section S2.1, our model implementation parameterizes $\sqrt{\mathcal{S}} := F'$ and $p_{prior}$ via the logit transformation. To account for subject-specific effects, we added these to the logit parameters:

$$\sqrt{J}(\theta|Subject_i) \propto \exp\Big(\alpha(\theta) + \alpha^{(i)}(\theta)\Big) \tag{17}$$

$$p_{prior}(\theta|Subject_i) \propto \exp\Big(\beta(\theta) + \beta^{(i)}(\theta)\Big) \tag{18}$$

Indeed, the literature on mixed-effects models contains models where the by-group adjustments are nonparametrically fitted functions [16]. The simplest adaptation of this approach to our setting is to assume an independent Gaussian prior for the adjustment to encoding or prior at each discretization point $\theta_i$. However, this approach does not encourage adjustments to be smooth, leading to substantial overfitting and potentially difficult-to-interpret discontinuous fits. We instead considered a prior that encourages smoothness:

$$P(\alpha^{(i)}|\mu=0;\sigma^2) = \frac{1}{Z_\sigma}\exp\left(-\frac{\|\frac{\mathrm{d}}{\mathrm{d}\theta}\alpha^{(i)}\|_2^2}{2\sigma^2}\right) \tag{19}$$

[8] Whereas (10) was derived by applying implicit differentation to the loss function, (15) can be derived by applying implicit differentiation to $P$, starting from the equation $P'(\theta_{MAP}) = 0$: Let $\theta_P := \operatorname{argmax} P$; then, at $\varepsilon = 0$ and for any test function $\phi$, we have $0 = \int \frac{\delta P'(\theta_P)}{\delta P}\phi dx = \frac{d}{d\varepsilon}(P+\varepsilon\phi)'(\theta_{P+\varepsilon\phi}) = \frac{d}{d\varepsilon}[P'(\theta_{P+\varepsilon\phi}) + \varepsilon\phi'(\theta_{P+\varepsilon\phi})] = P''(\theta)\frac{d}{d\varepsilon}\theta_{P+\varepsilon\phi} + \phi'(\theta_P) = P''(\theta_P)\int\frac{\delta\theta_P}{\delta P}\phi dx + \phi'(\theta_P) = \int\Big(P''(\theta_P)\frac{\delta\theta_P}{\delta P} - \delta'(x-\theta_P)\Big)\phi dx$. Setting the last term equal to zero for any test function $\phi$ yields the result. The resulting functional derivative is a tempered distribution, not an ordinary function; we note that, in contrast, for $p > 1$, the functional derivative is an ordinary function, analoguous to (10).

[9] On the finite-dimensional function space parameterized by $P_1,\dots,P_N$, the functional $\delta'(x-\theta)$ is given by the linear form $[-g_1'(\theta),\dots,-g_N(\theta]$.

[10] Defined by

$$g_i(x) = \frac{\exp\Big(-\frac{(x-x_i)^2}{2\sigma^2}\Big)}{\sum_{j=1}^N \exp\Big(-\frac{(x-x_j)^2}{2\sigma^2}\Big)}$$

with bandwidth $\sigma$ chosen to provide close fit to the discrete posterior.

[11] We chose $-\frac{1}{10}\exp(-(x-\theta_{Min})) - \frac{1}{10}\exp(x-\theta_{Max})$. This can be viewed as another kernel function $g_i$, and thus its contribution to the gradients can be absorbed into (16). While it would be possible to simply constrain $\widehat{\theta}$ to be in the stimulus space, such an approach would lead to incorrect gradients $\frac{\mathrm{d}}{\mathrm{d}P}\widehat{\theta}$ at the boundary.

where $\frac{1}{Z_\sigma}$ is the normalization constant. A penalty on the norm of $\alpha^{(i)}$ itself is not necessary, as adding any constant to $\alpha^{(i)}$ does not alter the resulting resource allocation (or prior). For interval data, a discretized version of (19) can be achieved by placing a Gaussian prior on the differences between values at successive points (analogously to the smootheness-based regularization term 8). However, for circular data, a naive application does not lead to a correctly-normalized probability distribution. To solve this, we parameterized the adjustment using its discrete Fourier transform[12], $\alpha^{(i)}(\theta) = \sum_{k=1}^{K} (w_k \sin(k\theta) + v_k \cos(k\theta))$; and utilized the fact that $\|\frac{d}{d\theta}\alpha^{(i)}\|_2^2 \propto \sum_{k=1}^{K} k^2 \left(|w_k|^2 + |v_k|^2\right)$ to compute (19) including the correct normalization:

$$P(\alpha^{(i)}|\mu = 0; \sigma^2) = \frac{\pi^K K!}{(2\sigma^2)^K} \exp\left(-\frac{\sum_{k=1}^{K} k^2 \left(|w_k|^2 + |v_k|^2\right)}{2\sigma^2}\right) \tag{20}$$

or equivalently

$$\log P(\alpha^{(i)}|\mu = 0; \sigma^2) = \log\left(\pi^K K!\right) - K \log\left(2\sigma^2\right) - \frac{\sum_{k=1}^{K} k^2 \left(|w_k|^2 + |v_k|^2\right)}{2\sigma^2} \tag{21}$$

as the precision matrix has determinant $\frac{\pi^{2K}(K!)^2}{\sigma^{4K}}$. We chose $K = 50$, providing ample space for fine-grained fits at the level allowed by the discretization. The likelihood then is

$$\int_{\mathbb{R}^{2K}} \int_{\mathbb{R}^{2K}} \cdots \int_{\mathbb{R}^{2K}} \int_{\mathbb{R}^{2K}} \exp(4) \prod_{i=1}^{\#Subjects} P(\alpha^{(i)}|\mu = 0, \sigma_\alpha^2) \prod_{i=1}^{\#Subjects} P(\beta^{(i)}|\mu = 0, \sigma_\beta^2) d\alpha^{(1)} d\beta^{(1)} \dots d\alpha^{(\#Subjects)} d\beta^{(\#Subjects)} \tag{22}$$

As this integral is intractable, we fitted parameters using ELBO variational inference [4, 17] and report negative log-likelihood of the fitted model at $\beta^{(i)}, \alpha^{(i)} \equiv 0$, i.e., evaluating only the fixed-effects structure.

#### S2.2 Plotted Quantities

For the human data, average responses and response SD are calculated for each subject at each $\theta_i$. The resulting curves are then smoothed using a Gaussian (for scalar stimuli) or von Mises (for circular stimuli) kernel.

We compute model predictions for variability as

$$\sqrt{\mathrm{SD}(\widehat{\theta})^2 + \rho^2} \tag{23}$$

where $\rho^2$ is the motor variance and where again $\mathrm{SD}(\widehat{\theta})$, the standard deviation of the estimate $\widehat{\theta}$ conditional on the stimulus $\theta$, is computed using circular statistics when the stimulus space is circular[13].

We compute the resource allocation $\sqrt{\mathcal{J}}$ as follows. We write $\mathrm{Vol}(\mathcal{X})$ for the size of $\mathcal{X}$; for an interval, it is $max(\mathcal{X}) - min(\mathcal{X})$; for orientation and direction perception (Main text, Figures 3–4), $\mathrm{Vol}(\mathcal{X}) = 180$. At $i = 1, ..., N-1$, write

$$V_i = F(\theta_{i+1}) - F(\theta_i) \tag{24}$$

so

$$F'(\theta_i) \approx \frac{F(\theta_{i+1}) - F(\theta_i)}{\theta_{i+1} - \theta_i} = \frac{V_i N}{\mathrm{Vol}(\mathcal{X})} \tag{25}$$

As $F$ is represented in the implementation as the cumulative sum of the discretization of $\sqrt{\mathcal{S}} = F'$, $F$ is internally represented in terms of $V_i$. Based on this, for an interval variable, we represent $\sqrt{\mathcal{J}(\theta_i)} = \frac{F'(\theta_i)}{\mathrm{SD}(m|\theta_i)}$ as [14]

$$\frac{V_i N}{\sigma \mathrm{Vol}(\mathcal{X})} \tag{29}$$

[12]The zeroth-order coefficient is not relevant, as adding any constant does not alter the resulting resource allocation (or prior).

[13]The circular SD is given as $\sqrt{-2\ln(\overline{R})}$, where $\overline{R}$ is the Eucliden distance between the origin and the vector mean of the angles.

[14]In the more general case where $\mathcal{Y}$ has length different from 1 and the Gaussian may be implemented with an extra factor for numerical stability, we have

$$\sqrt{\mathcal{J}(\theta_i)} = \frac{F'(\theta_i)}{\mathrm{SD}(m|\theta_i)} = \frac{V_i}{\sigma} \frac{N}{\mathrm{Vol}(\mathcal{X})} \frac{1}{\alpha} \tag{26}$$

For a circular variable, where the encoding follows a von Mises distribution with parameter $\kappa$, $\sqrt{\mathcal{J}(\theta_i)}$ can be represented analogously as [15]

$$\frac{V_i\sqrt{\kappa}N}{\mathrm{Vol}(\mathcal{X})} \tag{33}$$

when noise is small. In the presence of stimulus noise with variance $\tau^2$, this is extended to:

$$\sqrt{\mathcal{J}(\theta_i)} \approx \sqrt{\frac{1}{\tau^2 + \frac{1}{(\text{Equation 33})^2}}} \tag{34}$$

---

where

$$p(m|\theta_i) \propto \exp\left(-\frac{1}{2\sigma^2}\left|\frac{m - F(\theta_i)}{\alpha}\right|^2\right) \tag{27}$$

and thus

$$Var(m|\theta_i) \approx \sigma^2\alpha^2 \tag{28}$$

[15] In the more general case where $\mathcal{Y}$ may be indexed differently – e.g., as $[0,\pi]$ instead of $[0,2\pi]$ – this becones

$$\sqrt{\mathcal{J}(\theta_i)} = \frac{F'(\theta_i)}{\mathrm{SD}(m|\theta_i)} = V_i\sqrt{\kappa}\frac{N}{\mathrm{Vol}(\mathcal{X})}\frac{2\pi}{\mathrm{Vol}(\mathcal{Y})} \tag{30}$$

because

$$p(m|\theta_i) \propto \exp\left(\kappa \cdot \cos\left((m - F(\theta_i))\frac{2\pi}{\mathrm{Vol}(\mathcal{Y})}\right)\right) \tag{31}$$

and hence

$$Var(m|\theta_i) \approx \frac{1}{\kappa}\left(\frac{\mathrm{Vol}(\mathcal{Y})}{2\pi}\right)^2 \tag{32}$$

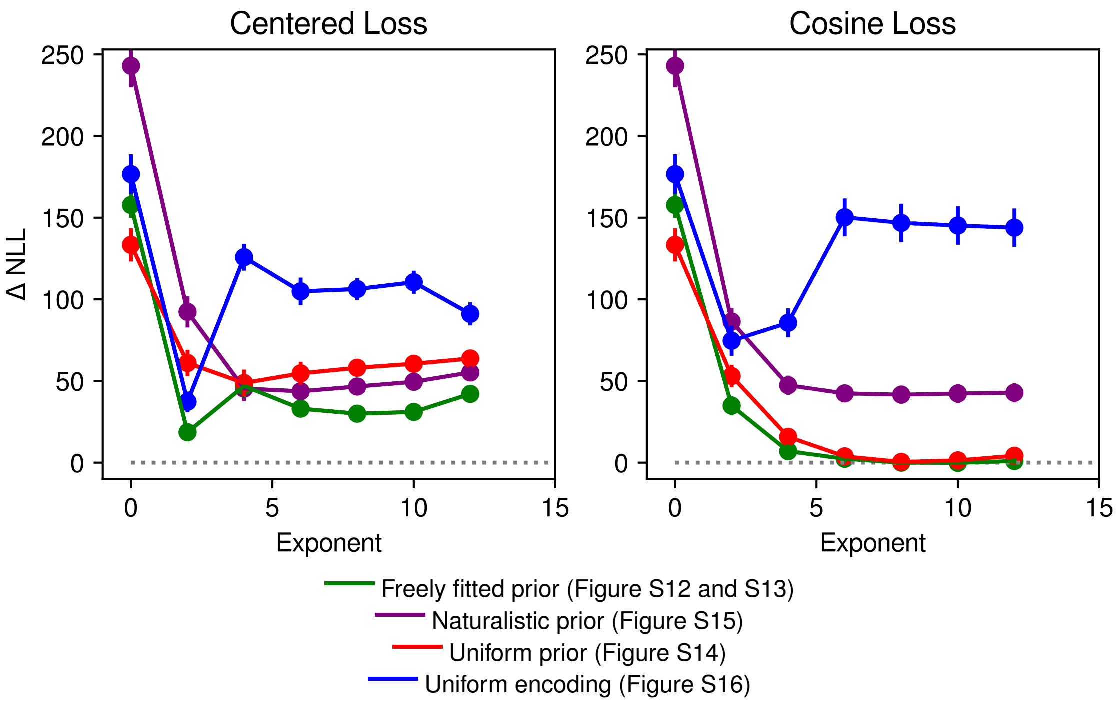

Figure S10: Orientation perception [data collected by 8]: Negative Log-Likelihood compared to best-fitting model as a function of the loss function exponent (lower is better). Dotted line indicates the best achieved fit. Left: Ordinary $L^p$ distance centered at $F^{-1}(m)$. Right: cosine-based distance. The cosine-based implementation of the loss function achieves lower Negative Log-Likelihood, in particular, a model with uniform (red) or freely fitted (green) prior. Note that $p = 2$ with freely fitted prior or uniform encoding can achieve relatively good Negative Log-Likelihood (though not optimal) in the left panel, but at the price of fitting an implausible prior peaking at oblique directions, at odds with both natural scene statistics or experiment-internal stimulus statistics (cf. Figure S13, $p = 2$). *Note: At $p = 0$, a uniform prior outperforms the freely fitted prior, suggesting slight overfitting in the latter. We did not observe such a phenomenon at other exponents.*

#### S2.3 Details and Results for Analyzed Datasets

##### S2.3.1 Orientation Perception

**S2.3.1.1 Orientation Perception (de Gardelle et al. [8])** Figure S10 shows model fit statistics, with references to further results.

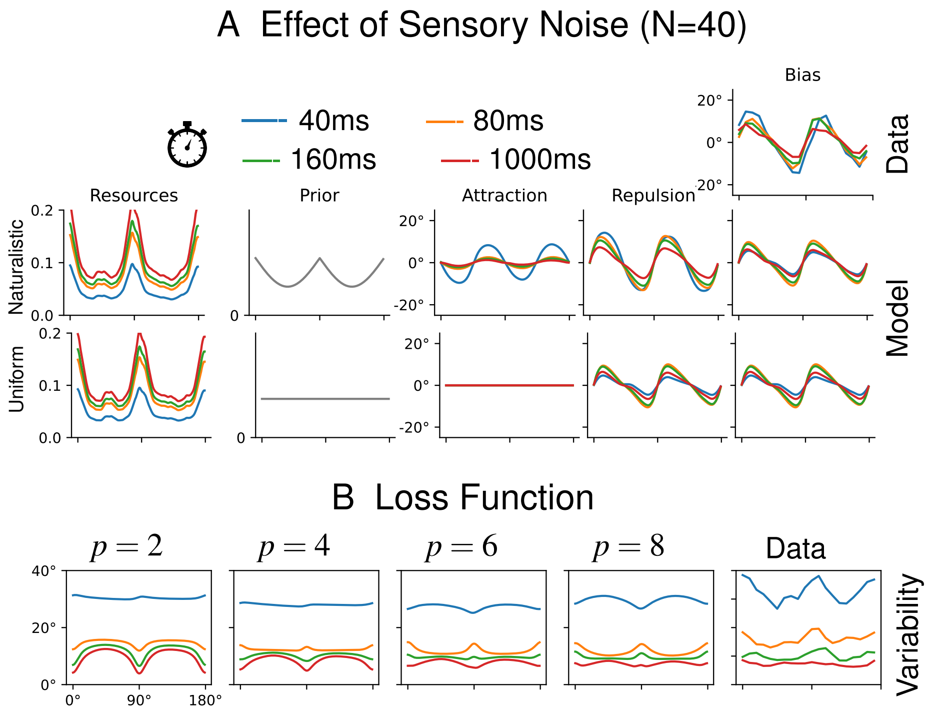

Figure S11: Orientation perception [data collected by 8] using the centered implementation of $L^p$ loss instead of the cosine-based implementation of loss function, corresponding to Figure 3A–B in the main text. Unlike the cosine-based implementation, this version cannot fit the variability well (B), accounting for its inferior quantitative fit compared to the cosine-based implementation (Figure S10).

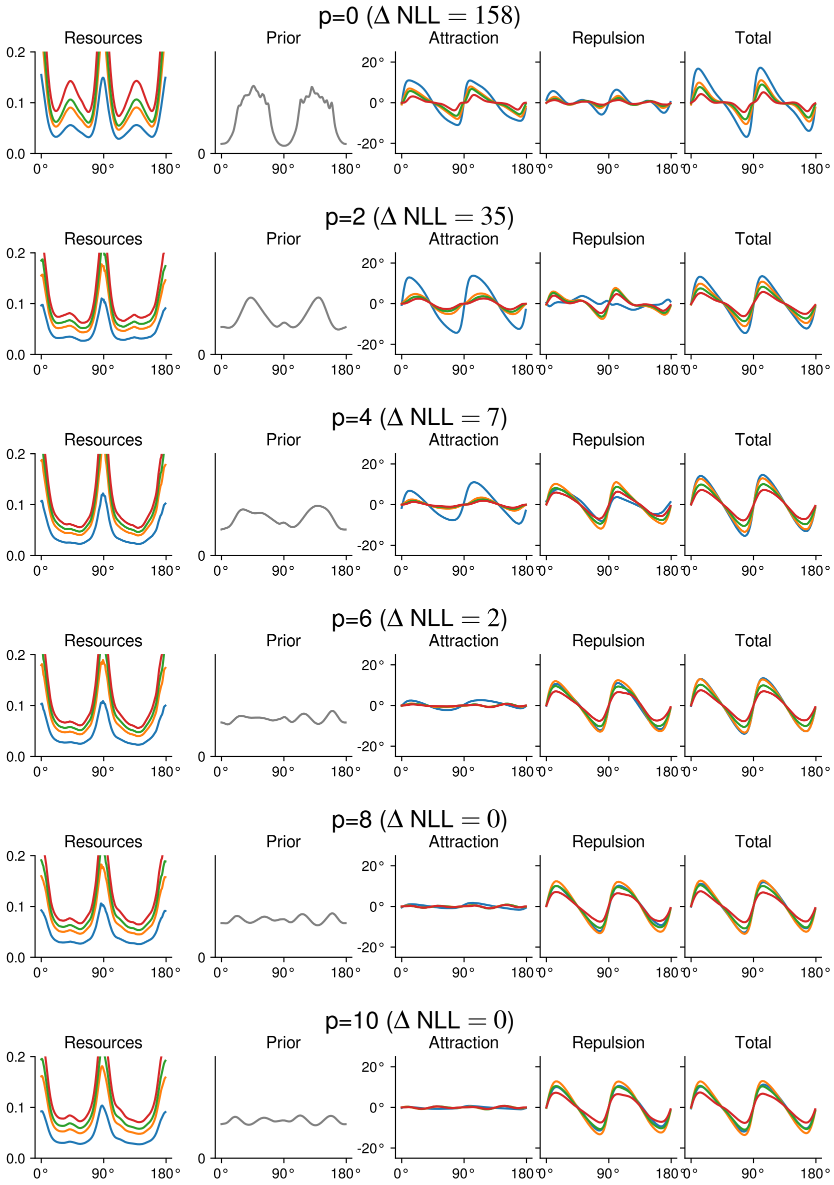

Figure S12: Orientation perception [data collected by 8]: Freely fitted prior (see Figure S12 for corresponding result with centered implementation of loss function). For each model, we show the increase in negative log-likelihood (NLL) compared to the best-fitting model reported in the main text; higher Δ NLL corresponds to poorer fit. For small exponents as assumed in prior work on orientation perception [12, 31], an implausible prior peaking in oblique directions results. For higher exponents, such as p=8, which achieve better quantitative model fit (see Figure S10), an approximately uniform prior is identified.

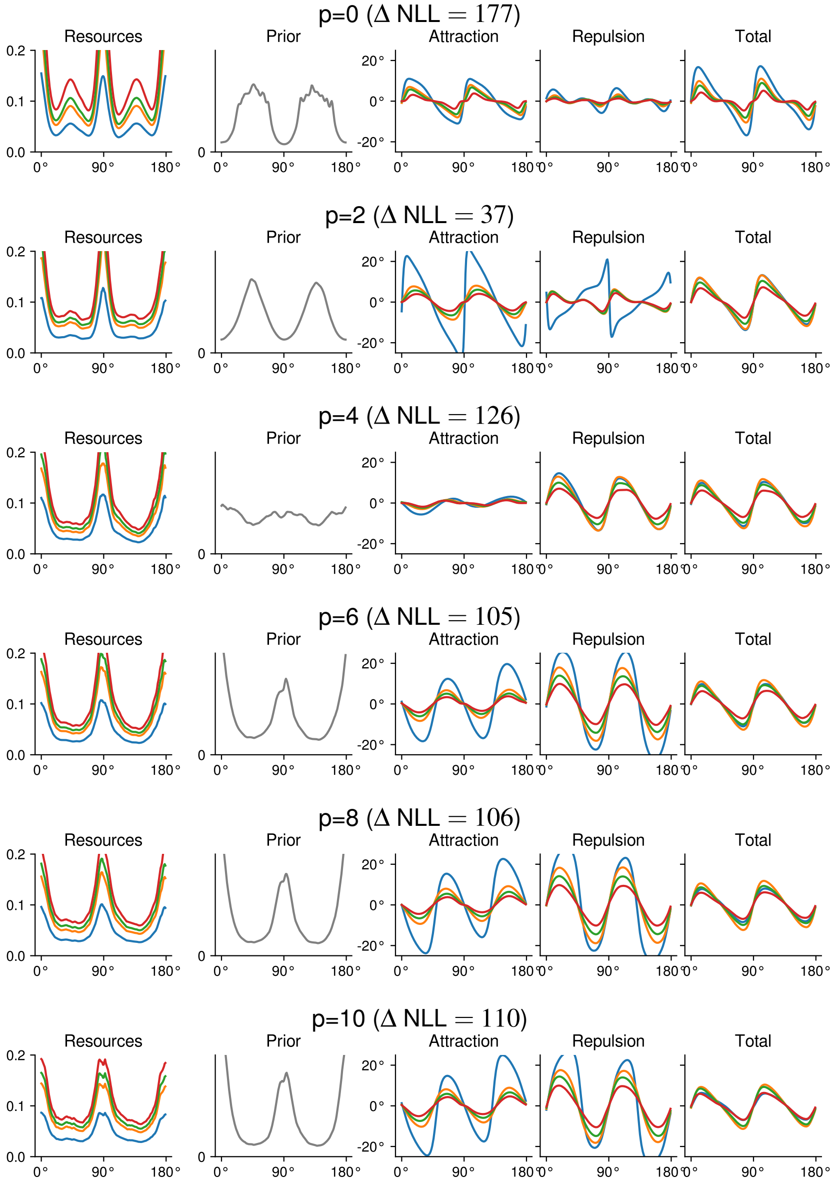

Figure S13: Orientation perception [data collected by 8]: Freely fitted prior, with centered implementation of loss function (see Figure S12 for corresponding result with cosine-based implementation). For small exponents as assumed in prior work on orientation perception [12, 31], an implausible prior peaking in oblique directions results. For higher exponents, the fitted prior peaks at cardinal directions in accordance with natural image statistics, though the best-fitting models show a uniform prior and use the cosine-based implementation (see Figure S12).

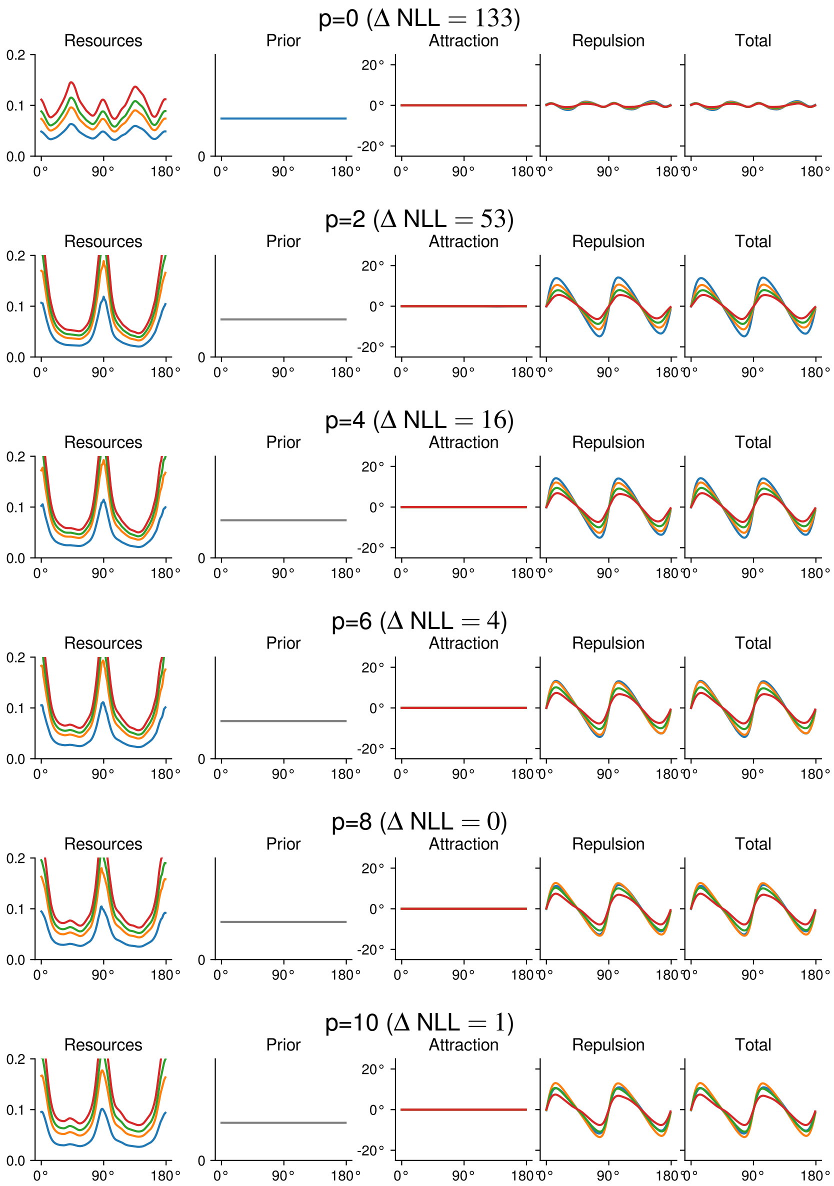

Figure S14: Orientation perception [data collected by 8]: Uniform prior, cosine-based implementation of $L^p$ loss.

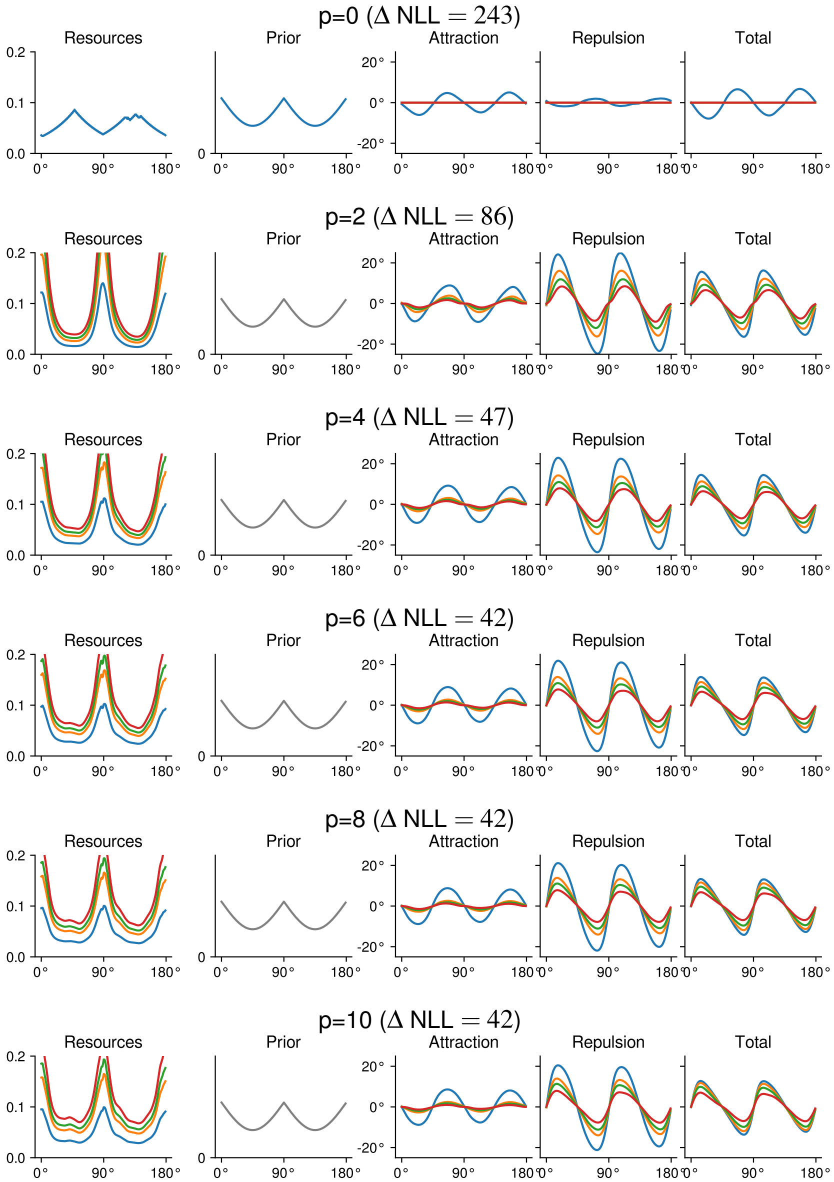

Figure S15: Orientation perception [data collected by 8]: prior based on natural image statistics, cosine-based implementation of $L^p$ loss. *Note: Fit at p=0 may look counterintuitive; we verified that a qualitatively equivalent fit was found even when initializing at the p=2 fit.*

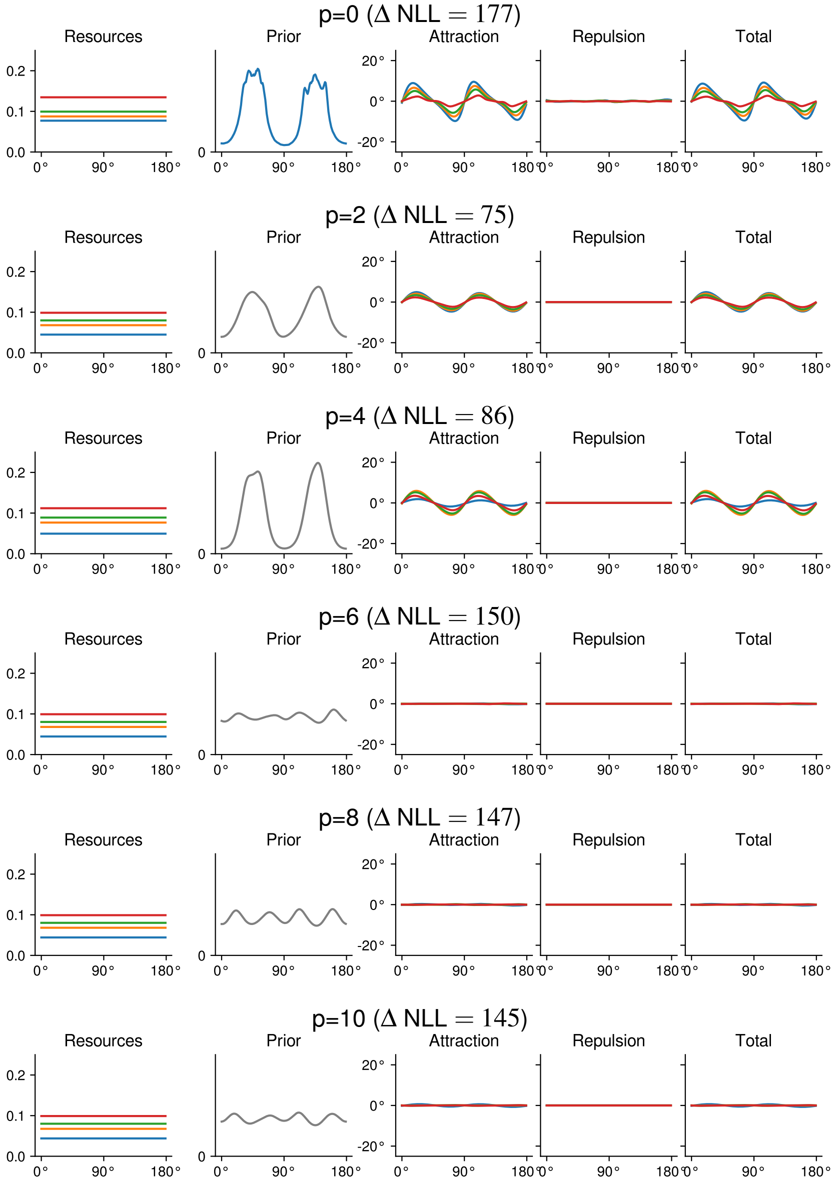

Figure S16: Orientation perception [data collected by 8]: uniform encoding, cosine-based implementation of $L^p$ loss. *Note: It may be surprising that almost zero bias is fitted at high exponents. We verified that the fitted flat prior is not due to the presence of regularization: The model simply has no way of accounting for both bias and variability if encoding is uniform.*

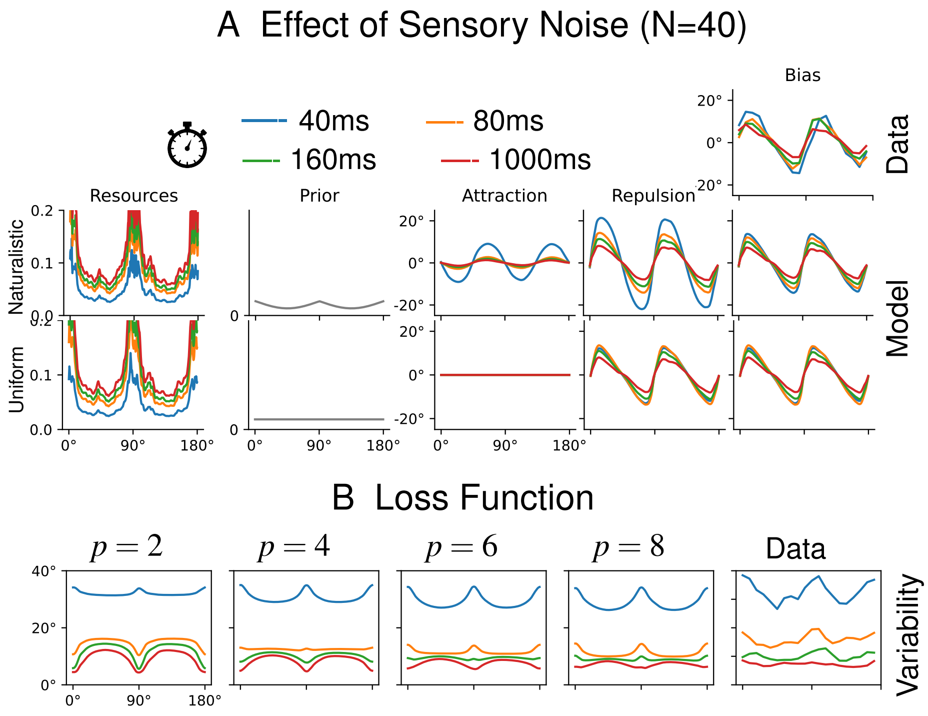

Figure S17: Orientation perception [data collected by 8]: Effect of the size of the discretization grid, at the example of orientation perception: Results corresponding to Main Text, Figure 3 A–B, with a discretized grid of size 720 instead of 180. Results closely match those reported in the Main Text.

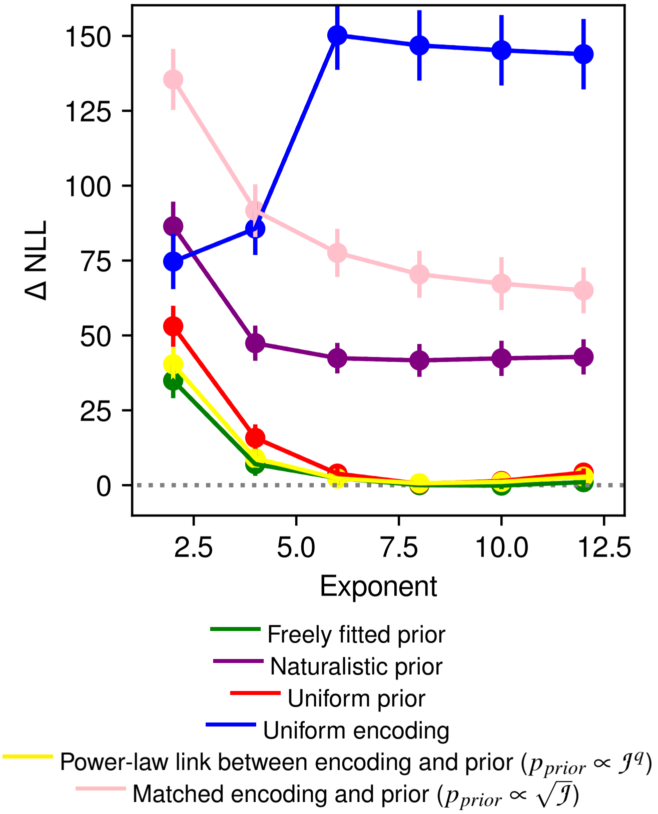

Figure S18: Orientation perception [data collected by 8]: Follow-up analyses for orientation perception (compare Figure S10), with cosine-based loss and restricting to $p > 0$. When all model components are freely fitted, the fitting procedure identifies encoding resource allocation peaking at the cardinals in agreement with prior work, and an approximately uniform prior, unlike assumed in previous models [12, 31]. However, this does not rule out that priors more in line with prior work, i.e., with a shape analogous to the encoding, could fit the data similarly well. To test this, we fitted two additional models: First, we considered a model where prior and encoding are directly matched (pink), which subsumes the prior and encoding assumed by [12, 31]; this one achieves poor quantitative fit. Second, we fitted a model where prior and encoding are linked by a power law [19]. This model achieves near-optimal fit (yellow); however, the resulting fit closely agrees with that obtained when both prior and encoding are fitted independently shown in Figure S12: $q$ is fitted to be *negative* at $p = 2, 4$, and approximately zero for higher exponents. In conclusion, the data are better explained by a model with a uniform prior than by models where both prior and encoding have similar shapes, peaking at the cardinals, assumed in prior work.

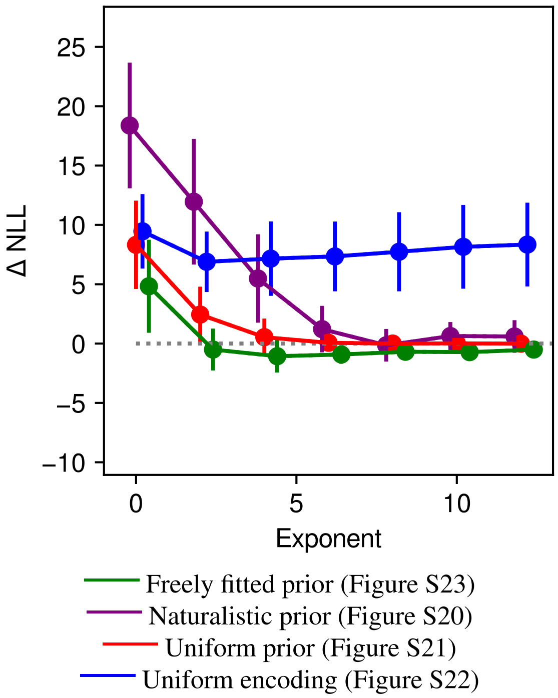

Figure S19: Orientation perception [data collected by 29]: negative Log-Likelihood relative to the model plotted in Main Text, Figure 2 as a function of the loss function exponent. We only considered the cosine-based implementation of $L^p$ loss, as it showed better fit in the larger dataset collected by de Gardelle et al. [8] (Figure S10).

###### S2.3.1.2 Stimulus Noise in Orientation Perception (Tomassini et al. [29])

**Details** Setup of the stimulus and sensory space is as for the data from de Gardelle et al. [8]. The dataset has three exposure durations; we fitted separate sensory noise variances to the three durations. In visualizations, we followed the original paper in pooling the three durations; there are not enough human observations per condition to fit a curve to the human biases. In accordance with the fit to the data from de Gardelle et al. [8], the resulting fits consistently assign higher sensory variance to lower exposure durations.

**Results** Table S19 shows model performance as a function of loss function.

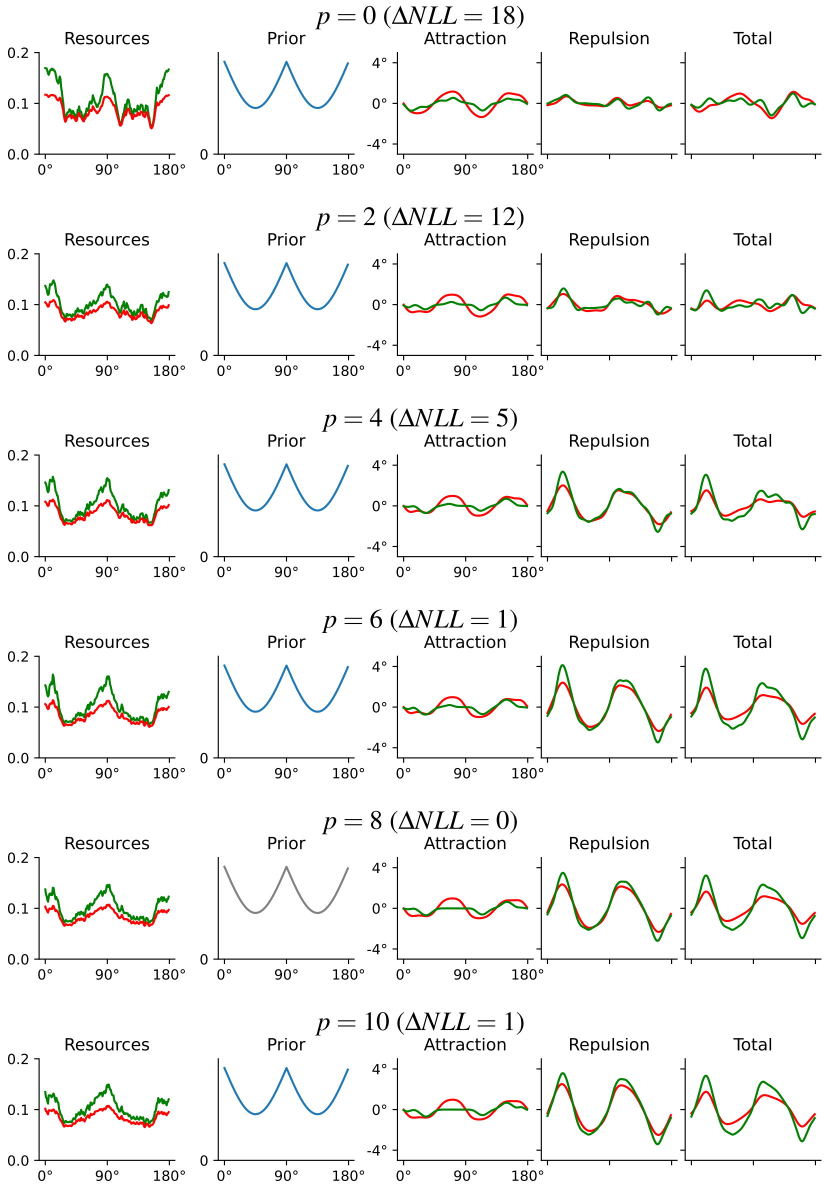

Figure S20: Orientation perception [data collected by 29]: prior based on natural image statistics

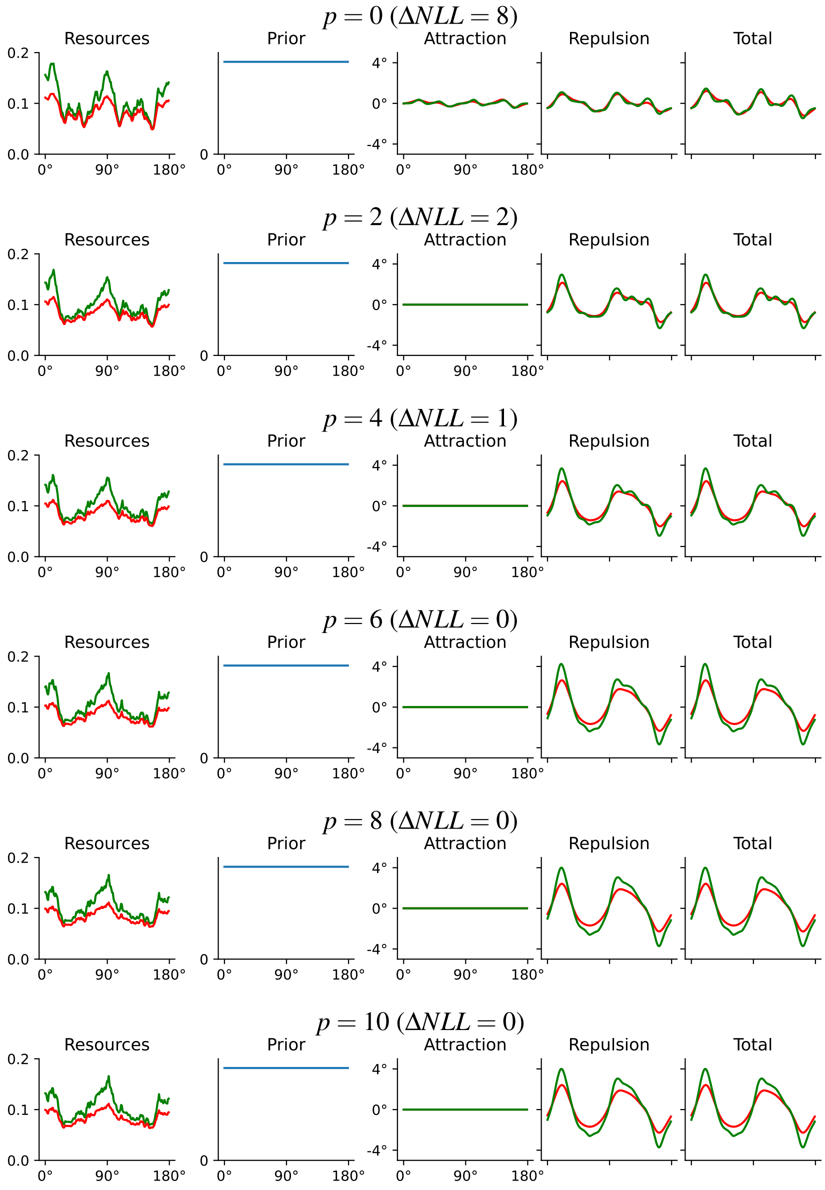

Figure S21: Orientation perception [data collected by 29]: uniform prior

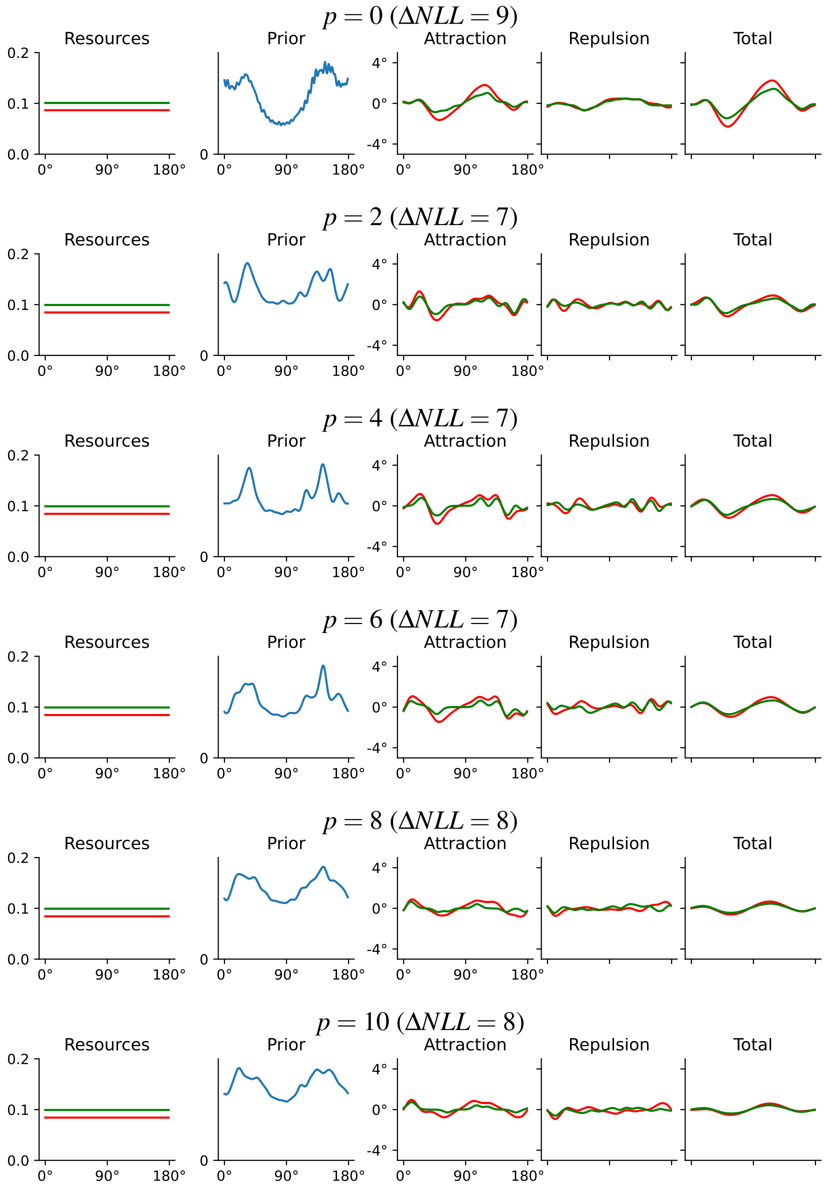

Figure S22: Orientation perception [data collected by 29]: uniform encoding

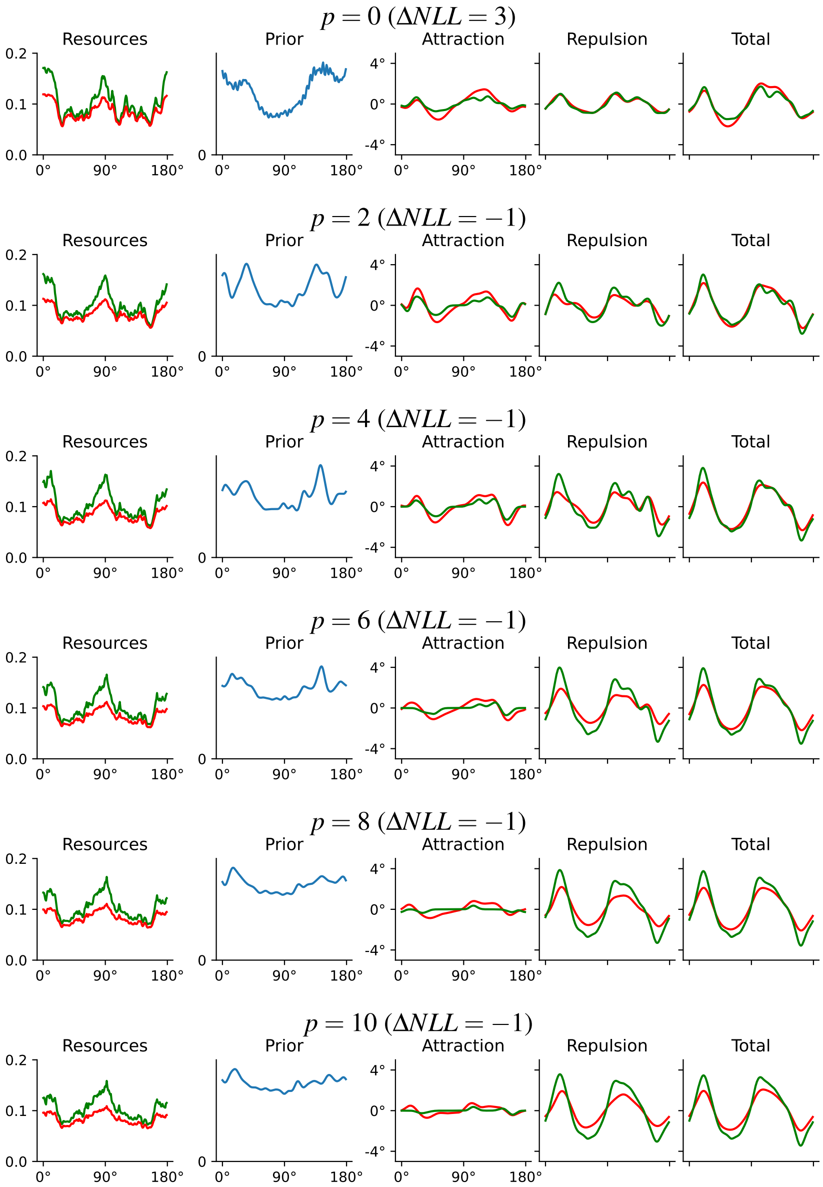

Figure S23: Orientation perception [data collected by 29]: Freely fitted prior and encoding. Role of stimulus noise in orientation perception. In agreement with the results we obtained one the data collected by de Gardelle et al. [8] (shown in Figure S12), exponents $> 2$ are needed to account for the pattern in biases, and the identified prior is approximately uniform. In agreement with Theorem 2, but unlike prior theories [12, 29], stimulus noise decreases biases towards obliques not due to increased prior attraction, but due to decreased likelihood repulsion. Accordingly, low exponents assumed in prior Bayesian modeling ($p = 0$ [12], $p = 2$ [31]) do not reproduce the observed reduction of biases due to stimulus noise well.

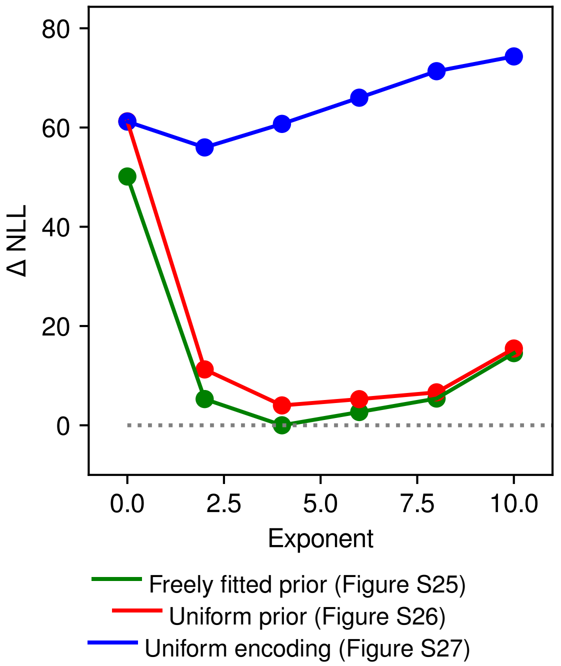

Figure S24: Perception of motion direction [data collected by 11]: Negative Log-Likelihood compared to the model reported in Main Text, Figure 4, as a function of the loss function exponent (lower is better). Due to computational cost of fitting per-subject parameters, the model was only run on one fold. While the MAP estimator cannot fit the data, larger exponents all provide similar fit. Assuming a uniform prior only slightly decreases the quality of model fit, whereas assuming a uniform encoding decreases it strongly.

###### S2.3.2 Motion Direction (Gekas et al. [11])

**Details** Within the relevant trials, the dataset has two contrast levels, for which we fitted different sensory noise parameters. In visualizations, we followed the original paper in pooling the two contrast levels. The resulting fit assigns higher sensory variance to the lower contrast level.

**Results** Table S24 shows model performance as a function of loss function.

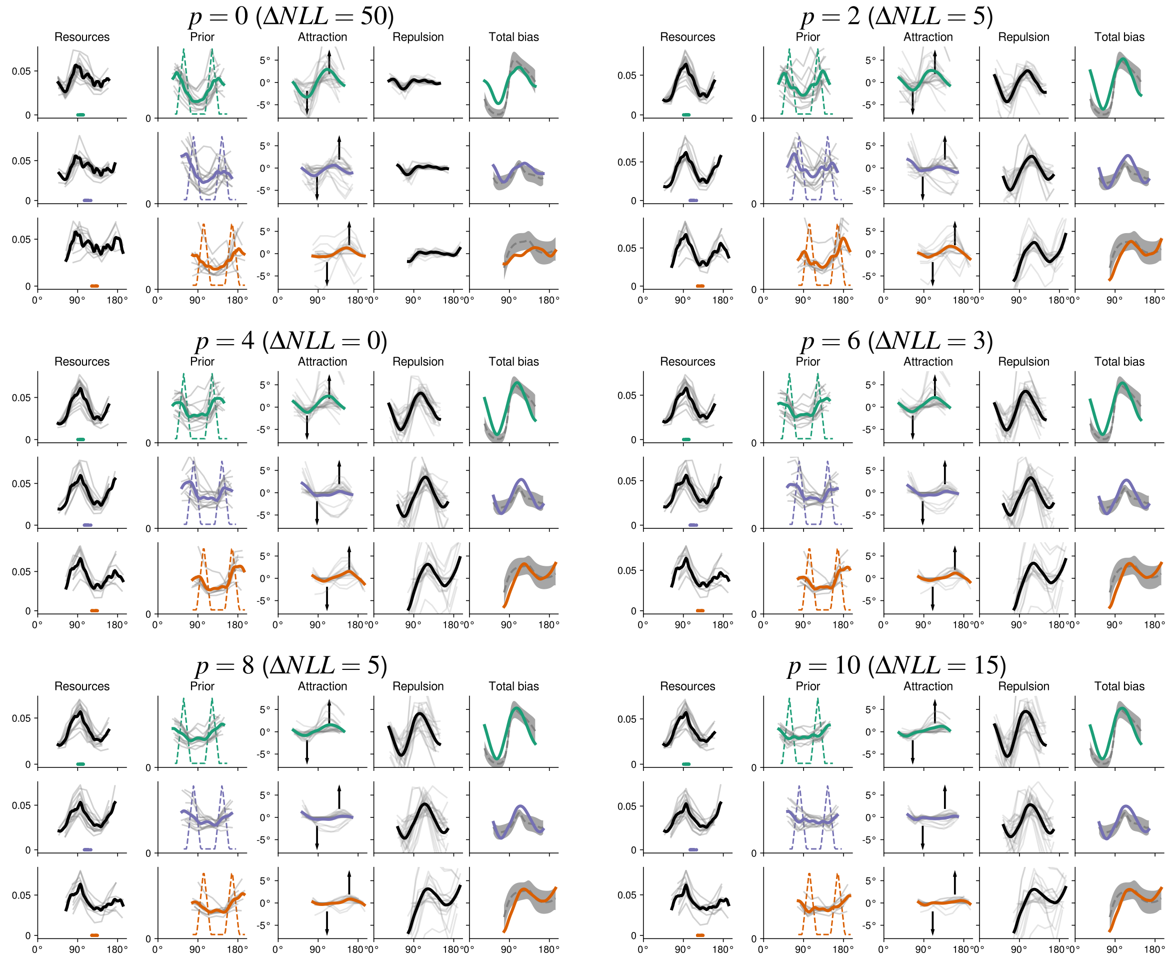

Figure S25: Perception of motion direction [data collected by 11]: Results corresponding to Main Text, Figure 4 A–C, across loss function exponents. A very similar qualitative pattern is predicted for all nonzero exponents.

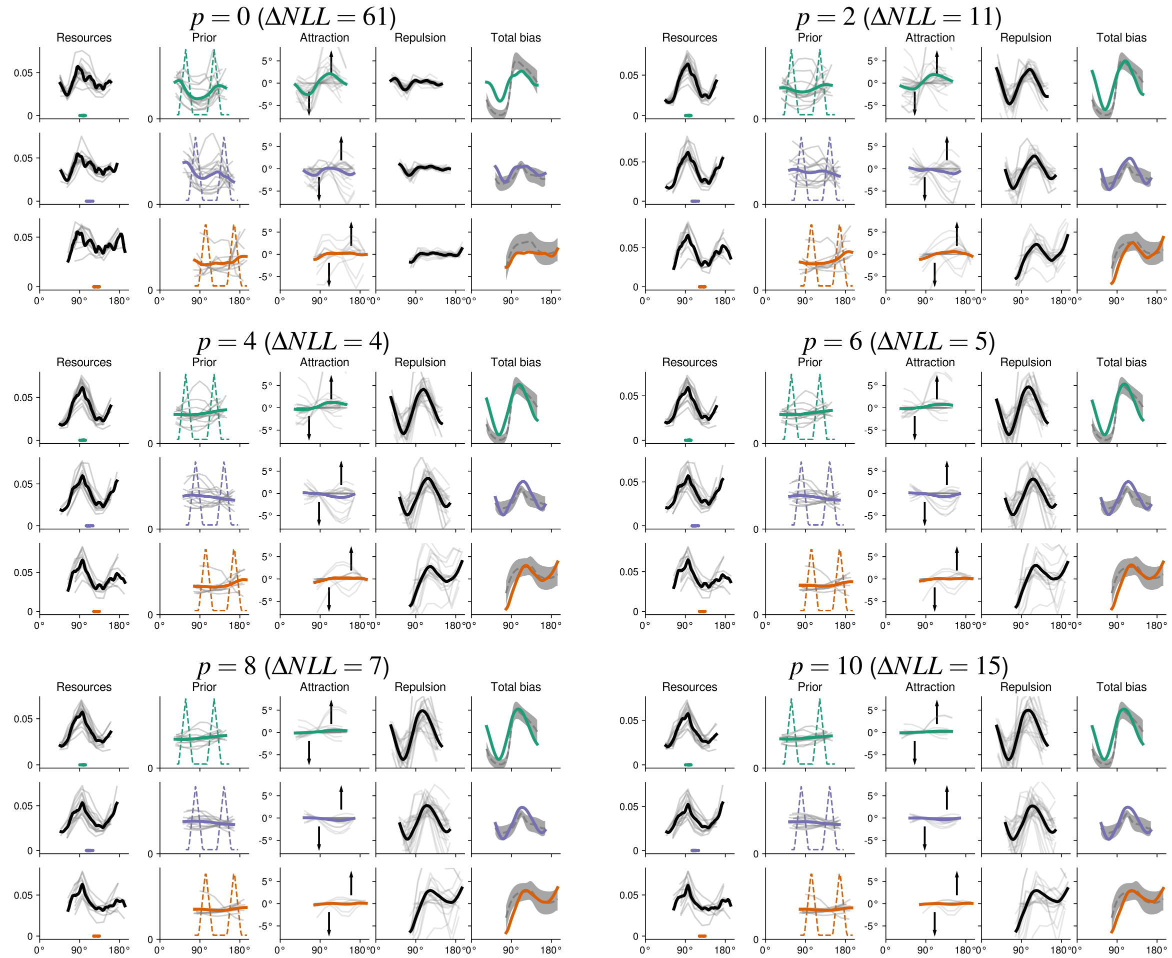

Figure S26: Perception of motion direction [data collected by 11]: Results corresponding to Figure S25, with the fixed-effects component of the prior ($\beta(\theta)$ in Equation 18) constrained to be zero. That is, nonuniformity in the prior is entirely relegated to per-participant adjustments. Results are similar to Figure S25. Forcing the fixed-effects component of the prior to be zero forces the model to assume a somwehat flatter resource allocation in the third row, in order to account for the difference in bias magnitude between the first and third rows.

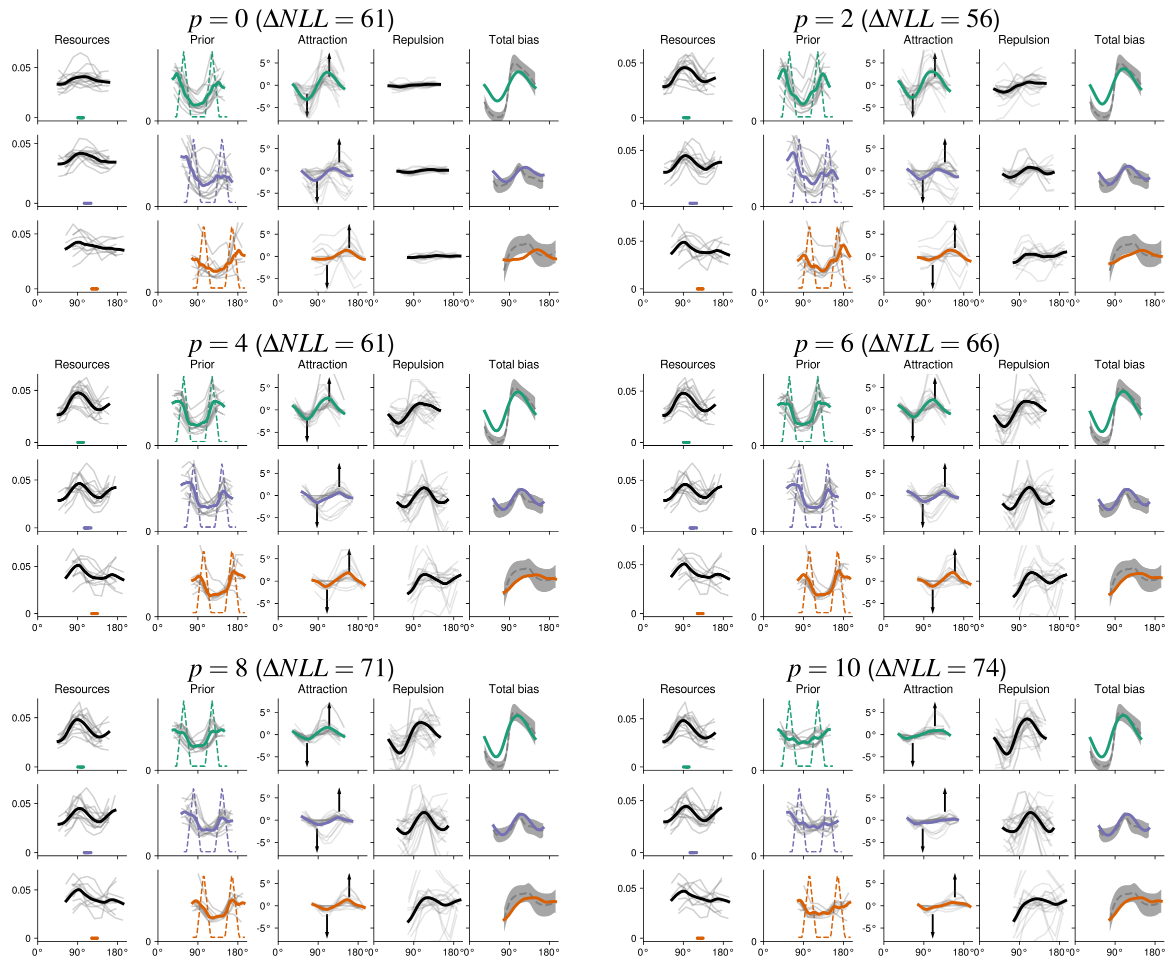

Figure S27: Perception of motion direction [data collected by 11]: Results corresponding to Figure S25, with the fixed-effects component of the prior ($\alpha(\theta)$ in Equation 17) constrained to be zero. That is, nonuniformity in the resource allocation is entirely relegated to per-participant adjustments. Per-participants adjustments broadly recover the oblique effect; the prior is also fitted consistently. This model can be viewed as an instantiation of the modeling considered in Gekas et al. [11] but with participant-specific adjustments. Those adjustments highlight the importance of the nonuniform resource allocation, and explain why the modeling of Gekas et al. [11] did not capture the qualitative pattern in the data.

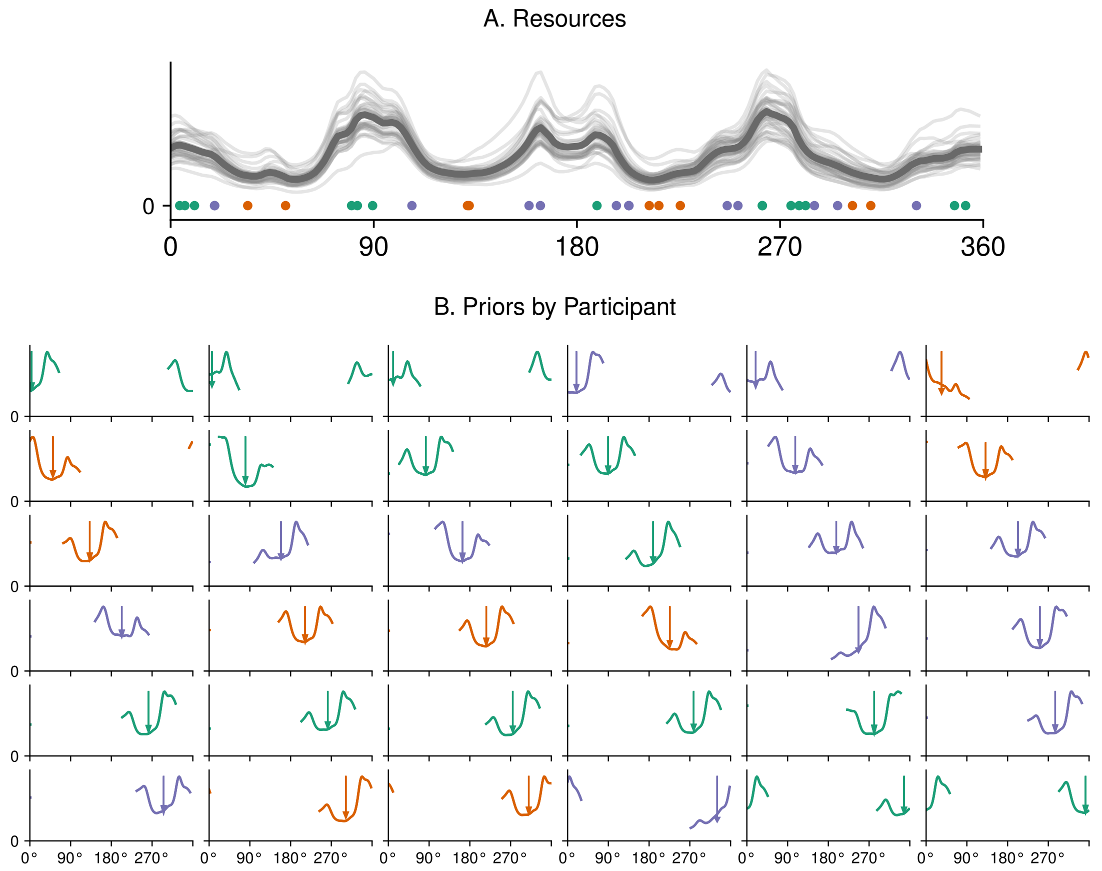

Figure S28: Perception of motion direction [data collected by 11]: Fitted model for perception of motion direction, but without transforming central directions into $[90°, 135°]$. Participants are grouped based on distance to cardinal and oblique directions as in Main text, Figure 4. Top: By pooling across participants, the model identifies peaks in encoding resource allocation at the cardinal directions (same as Main text, Figure 4a). Bottom: Fitted prior for each of the 36 subjects. Black arrows indicate the central direction for each participant. A bimodal prior is identified for most participants.

##### S2.3.3 Magnitude Estimation

###### S2.3.3.1 General Notes

**Priors** In addition to nonparametrically fitting priors to response data, we further compared the following priors based on prior work. A uniform prior on an interval is assumed in the Bayesian models of Jazayeri and Shadlen [15], Remington et al. [25]. The model of Petzschner and Glasauer [21] represents a Kalman filter in the sensory space; the history from previous trials is effectively summarized into a normal prior in sensory space. Similarly, the model of Cicchini et al. [7] resembles a Kalman filter in stimulus space; it effectively summarizes the history from previous trials into a normal prior in stimulus space, whose mean is a weighted average of the preceding stimuli.

**Role of Repulsion** It is perhaps surprising that our modeling indicates a repulsive bias component towards higher magnitudes, even though magnitude perception often shows underestimation biases, and overestimation biases are rarely directly observed [e.g. perception of temporal frequency in Figure 2B in 30]. Indeed, such a repulsive bias is a necessary consequence of Bayesian estimation models when Weber's law holds and $p > 0$, and is correspondingly implicit in prior Bayesian models, though it may be hard to directly detect due to the presence of a large central tendency effect and/or a prior favoring smaller magnitudes, which can cancel or outweigh the repulsive component. For instance, Petzschner and Glasauer [21] fitted a "shift term" in the estimate in their Bayesian model; this shift term plays a role corresponding to the loss-dependent likelihood repulsion. Fitted values of the shift term indicate a nonzero (positive) repulsive bias (see below). Relatedly, the posterior mean estimator used, for instance, by Jazayeri and Shadlen [15] necessarily involves some nonzero amount of likelihood repulsion.

We note that there are models that do not involve likelihood repulsion because they are not strictly Bayesian models. The model of Cicchini et al. [7], which is motivated as a Bayesian integration model akin to the Kalman filter, does not involve a repulsive bias component; it only shows attraction to a weighted average of previous stimuli. Indeed, the assumptions of the standard Kalman filter are violated by the stimulus-dependence of noise in magnitude estimation (their Equation 5). Exact Bayesian inference for such a model may perhaps be carried out with a suitable nonlinear Kalman filter [27] and would give rise to a repulsive bias component in addition to the central tendency effect, similar to Equation 36. Investigating such a fully Bayesian model and comparing it to the model described by Cicchini et al. [7] is an interesting problem for further research on numerosity perception. Another model, involving only an underestimation bias, is presented by Cheyette and Piantadosi [5]; this model differs from other proposals in that it optimizes the psychophysical mapping from $\theta$ to $\widehat{\theta}$ "end-to-end" to minimize the average $L^2$ loss subject to constraints on KL divergence from the prior, and does not involve Bayesian inference over uncertain sensory input [6]. Bayesian variants of this model would need to include likelihood repulsion; comparing such variants with the model of [5] is likewise an interesting problem for future research on numerosity perception.

**Role of Repulsion in the model of Petzschner and Glasauer [21]** Here, we elaborate the role of likelihood repulsion implicit in the Bayesian model of magnitude perception by Petzschner and Glasauer [21]. In particular, we show that the "shift term" they fitted corresponds to loss-dependent likelihood repulsion. The model of Petzschner and Glasauer [21] implements Weber's law by assuming $F(\theta) = \log(\theta)$. The likelihood and the posterior are both lognormal. Given parameters $\mu$, $\sigma_{post}$ of the posterior, [21] assume the following functional form for the estimate $\widehat{\theta}$:

$$\widehat{\theta} = \exp(\mu + \Delta x) \cdot d_0 \tag{35}$$

($d_0$ is a proportionality coefficient with units of distance or angles), where $\Delta x = 0$ at $p = 1$, $\Delta x = \sigma^2_{post}/2$ at $p = 2$, and $\Delta x = -\sigma^2_{post}$ at $p = 0$, by standard facts about lognormal distributions. Further, by Gaussianity of prior and likelihood in sensory space, $\mu = w \cdot (F(\theta) + \delta) + (1 - w) \cdot \mu_{prior}$, where the weight $w$ only depends on the relative widths of prior and likelihood. With this in mind, we can derive the bias of the estimate in the model of [21] as follows (omitting the proportionality unit $d_0$)

$$\begin{aligned}\mathbb{E}_{\delta\sim\mathcal{N}(0,\sigma^2_{Lik})}\left[\exp(w(F(\theta)+\delta)+(1-w)\mu_{prior}+\Delta x)\right]-\theta =& \mathbb{E}_{\delta\sim\mathcal{N}(0,w^2\sigma^2_{Lik})}[\exp(\delta)]\cdot\exp(w\log\theta+(1-w)\mu_{prior}+\Delta x)-\theta \\ =& \exp\left(w\log\theta+\frac{w^2\sigma^2_{Lik}}{2}+(1-w)\mu_{prior}+\Delta x\right)-\theta\end{aligned}$$

To lowest order in $t := \sigma_{Lik}^2$, this is ($w = \frac{1/\sigma^2}{1/\sigma^2 + 1/\sigma_{prior}^2} = 1 - \frac{t}{\sigma_{prior}^2} + O(t^2)$; $\sigma_{post} = \frac{t\sigma_{prior}^2}{t+\sigma_{prior}^2} = t + O(t^2)$) by differentiating w.r.t. $t$ at $t = 0$:

$$Bias = \underbrace{\sigma_{Lik}^2 \cdot \theta \cdot \left[\frac{(\mu_{prior} - \log\theta)}{\sigma_{prior}^2} - 1\right]}_{\text{Prior Attraction}} + \underbrace{\sigma_{Lik}^2 \cdot \theta \cdot \left(\frac{3}{2} + \frac{\Delta x}{\sigma_{post}^2}\right)}_{\text{Likelihood Repulsion}} + O(\sigma_{Lik}^4) \tag{36}$$

This result agrees with the result obtained by directly applying our Theorem 1 to the model of [21], plugging in the lognormal prior and the encoding assumed by [21] ($\mathcal{J} = \frac{1}{\theta^2\sigma_{lik}^2}$), here $p > 0$ (similarly for $p = 0$):

$$\frac{1}{\mathcal{J}}(\log p_{prior})' + \frac{p+2}{4}\left(\frac{1}{\mathcal{J}}\right)' = \frac{1}{\mathcal{J}}\frac{\mathrm{d}}{\mathrm{d}\theta}\left[\log\frac{1}{\theta}\exp\left(-\frac{(\log\theta - \mu_{prior})^2}{2\sigma_{lik}^2}\right)\right] + \frac{p+2}{4}\left(\frac{1}{\mathcal{J}}\right)' \tag{37}$$

$$= \sigma_{Lik}^2 \cdot \theta \cdot \left[\frac{(\mu_{prior} - \log\theta)}{\sigma_{prior}^2} - 1\right] + \sigma_{lik}^2 \cdot \theta \cdot \frac{p+2}{2} \tag{38}$$

which matches (36) under the identification $\Delta x = \sigma_{post}^2 \cdot \left(\frac{p-1}{2}\right)$ (for $p > 0$, similarly for $p = 0$). The likelihood repulsion term in (36) is always positive, even when $p = 0$ ($\Delta x = -\sigma_{post}^2$). Petzschner and Glasauer [21] made $\Delta x$ a free parameter fitted together with the rest of the model; the fitted value was positive on average across subjects, indicating a loss function exponent $> 1$, and a corresponding repulsive bias. Note, however, that just as in our model fits (Figure 5 in the main text), the overall bias may still be predominantly underestimating because of the shape of the prior; in (36), prior attraction results in underestimation even when $\log\theta = \mu_{prior}$.

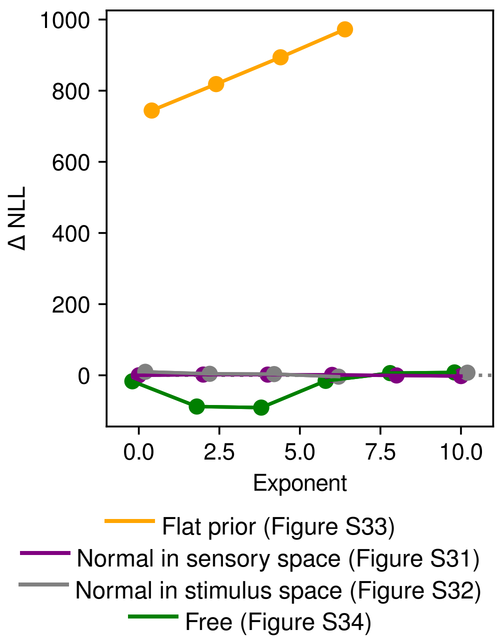

Figure S29: Magnitude estimation (analyzing data collected by Xiang et al. [33]). Negative Log-Likelihood as a function of the loss function exponent (lower is better), compared to the model reported in Main Text, Figure 4a. The model with freely fitted or normal (in stimulus space or sensory space) prior consistently improves over models with flat prior. *Note:* Numerical problems impacted freely fitting the model at $p \geq 6$. Hence, we initialized the model at the normal model and continued optimization; the green curve can only represent an upper bound in that range.

###### S2.3.3.2 Numerosity Estimation (Xiang et al. [33])

**Details** In order to avoid boundary effects, we considered the stimulus space $[0, 150]$. We used one grid point per integer in this range. We removed responses outside of the stimulus soace, removing 24 out of 71,408 observations (0.03% of data).

As the observations are discrete, we had to constrain the motor variability to exceed a minimum value chosen so that that per-trial likelihood was not numerically equal to one, preventing a situation where the gradients of the likelihood w.r.t. $\widehat{\theta}$ become numerically zero.

The normal-in-sensory-space priors varied only in their mean but not their variance, following Petzschner and Glasauer [21]. On the other hand, the normal-in-stimulus-space priors were allowed to vary in both parameters, following Cicchini et al. [7].

**Results** Model fit statistics are shown in Figure S29. Fitted priors for the 25 intervals are shown in Figure S30 (at $p = 2$). Parametric and nonparametric fits are qualitatively quite similar, in particular when visualized in sensory space (Figure S30). Unlike suggested by Cicchini et al. [7], the normal-in-stimulus-space prior has its mean parameter at small numbers, below the relevant range.

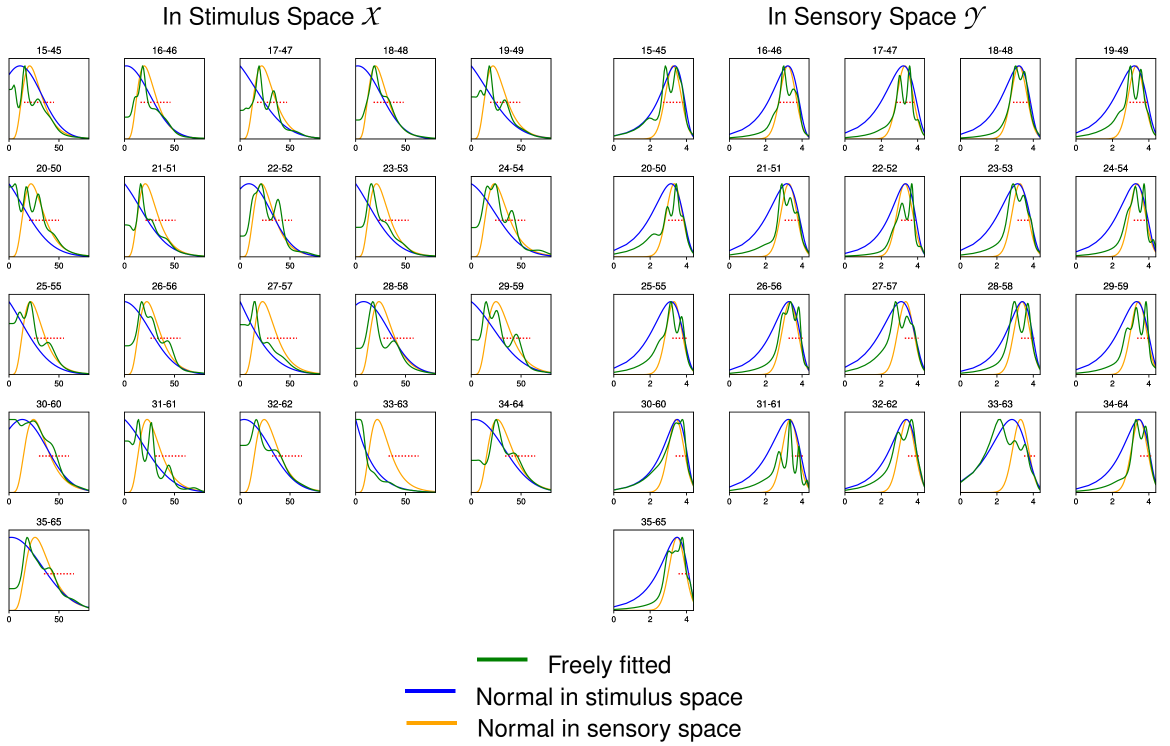

Figure S30: Fitted priors in numerosity estimation dataset of Xiang et al. [33], at $p = 2$, for each of the 21 intervals used in the experiment. All three types of prior achieve similar model fit. In sensory space, priors peak within or slightly below each interval. This is compatible with combination of an overall prior favoring lower numbers being combined with interval-specific priors.

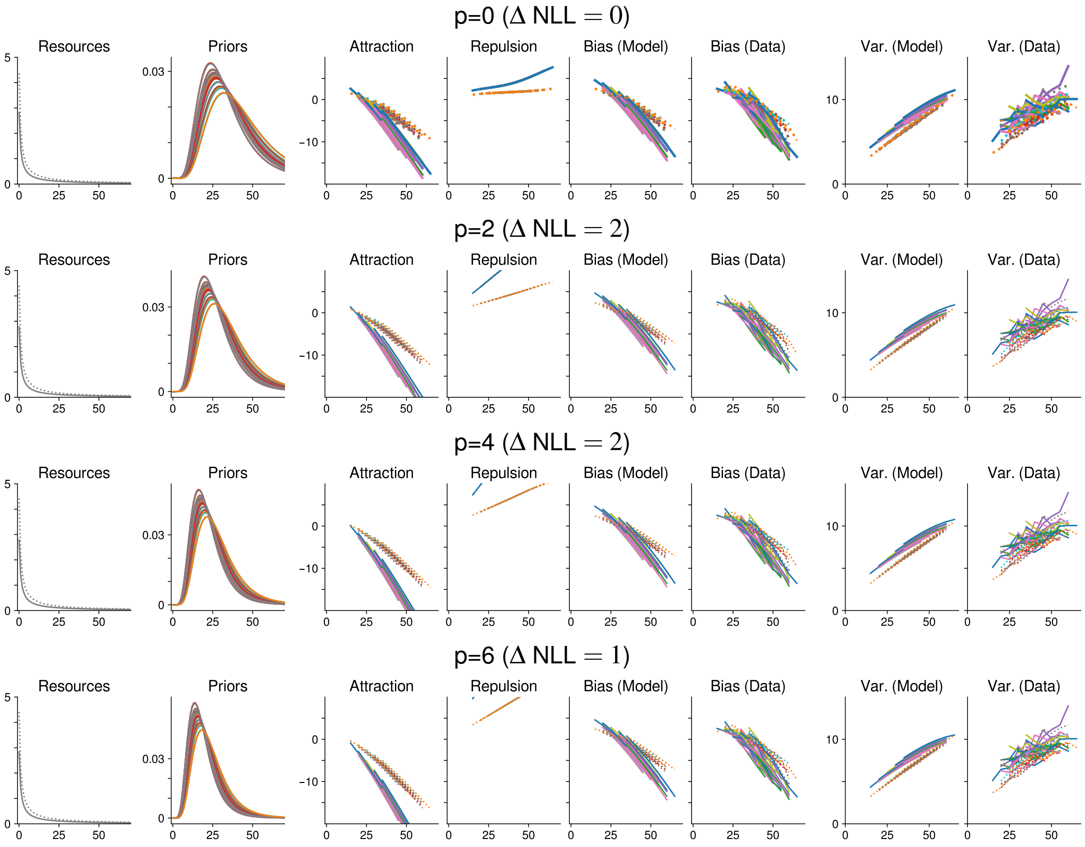

Figure S31: Numerosity estimation [data collected by 33]: normal (in sensory space) prior.

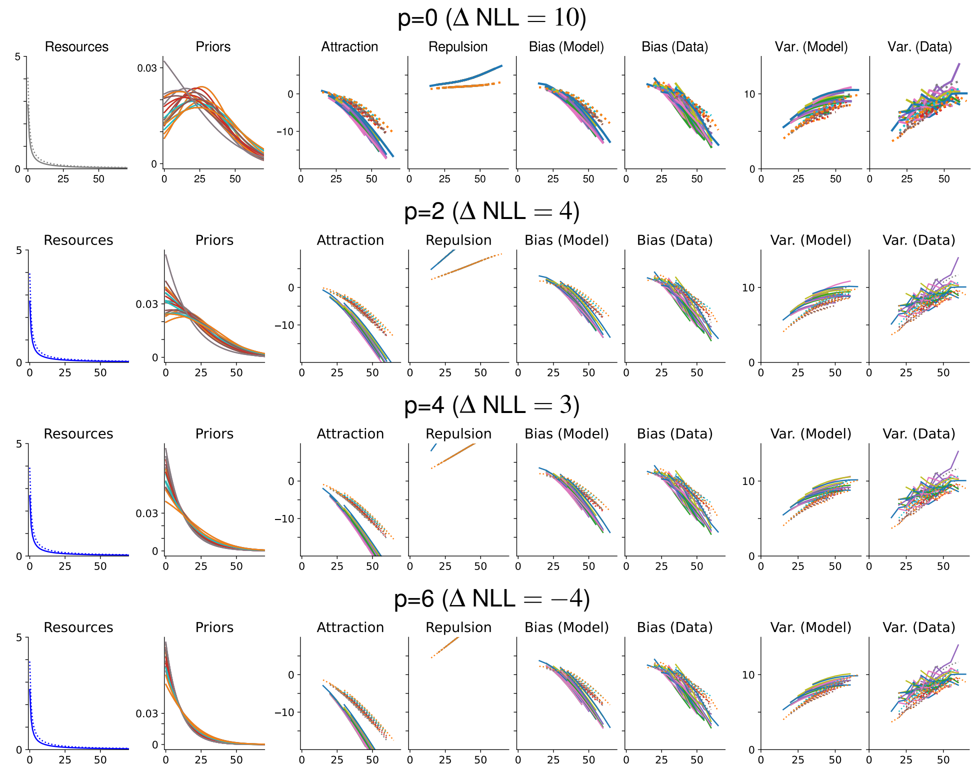

Figure S32: Numerosity estimation [data collected by 33]: normal (in stimulus space) prior.

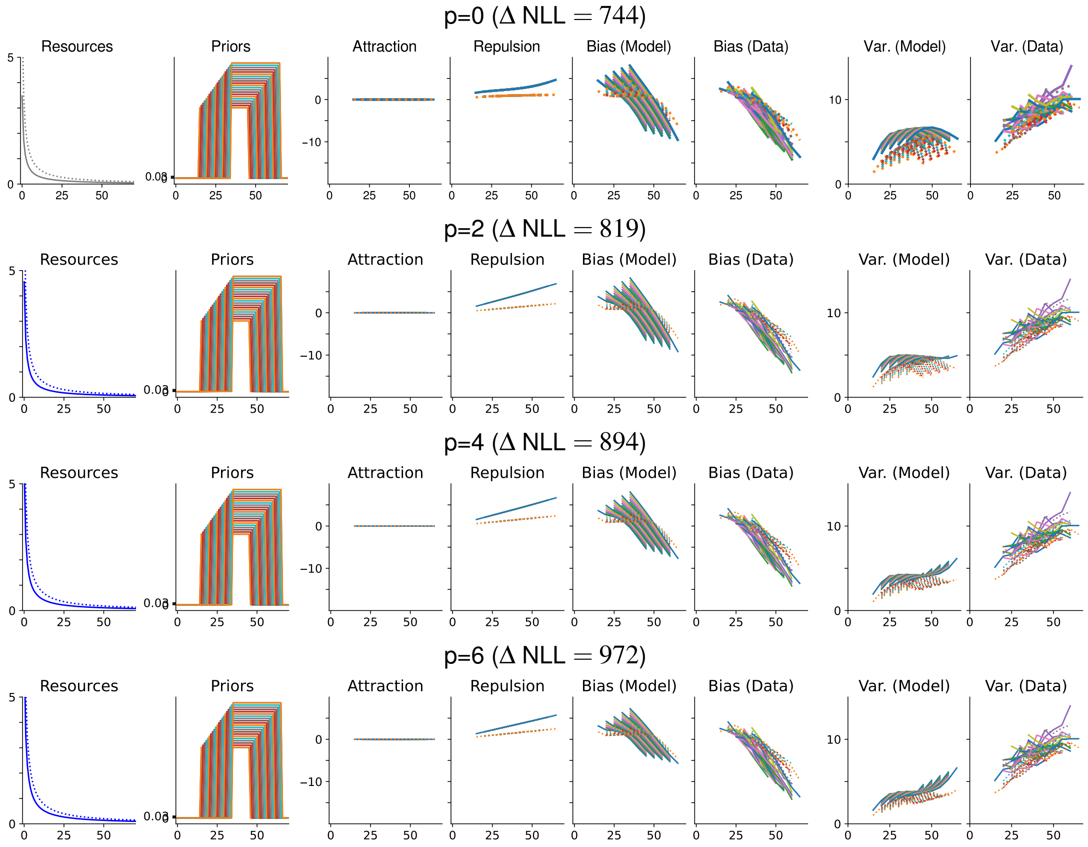

Figure S33: Numerosity estimation [data collected by 33]: flat prior

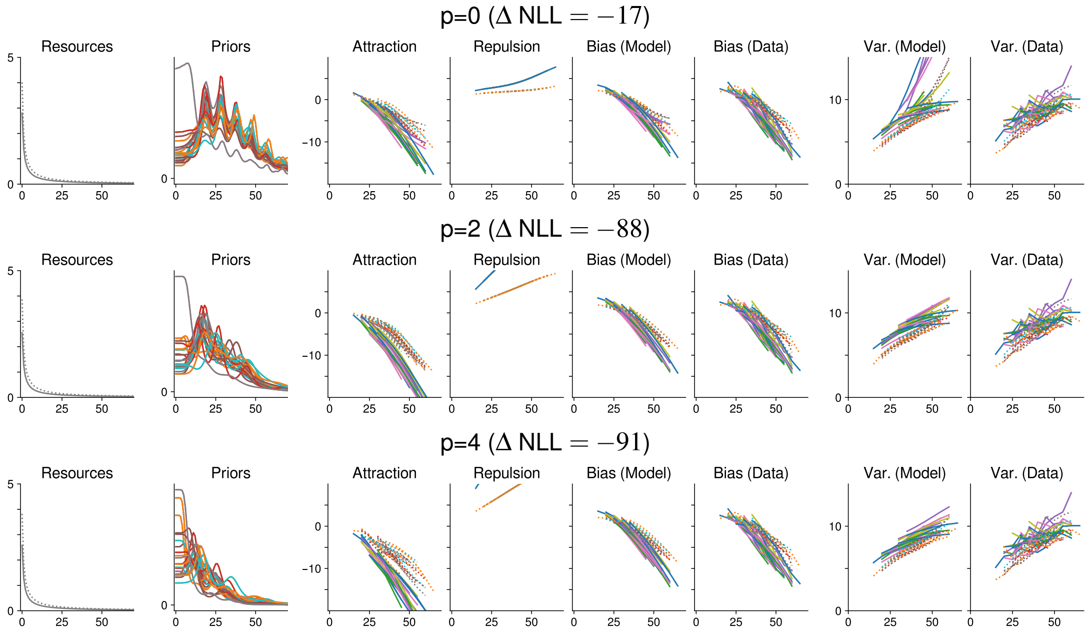

Figure S34: Numerosity estimation [data collected by 33]: freely fitted prior.

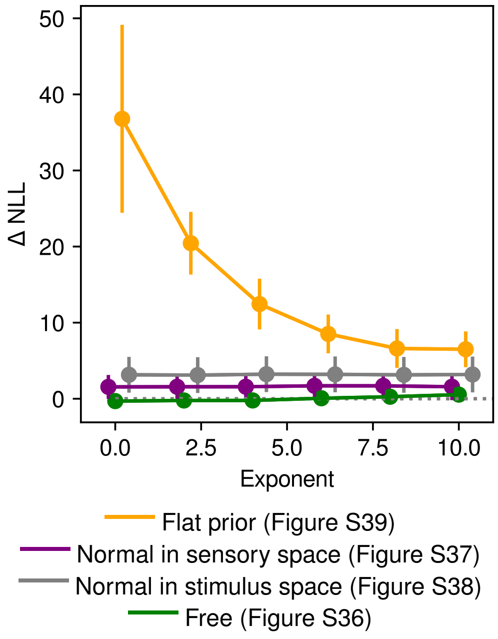

Figure S35: Time interval estimation (data from Remington et al. [25]). Difference in negative Log-Likelihood compared to the model reported in the main paper, as a function of the loss function exponent, averaged across the 10 folds (lower is better). Error bars indicate standard errors of the NLL (across the 10 folds). As predicted by our theory, the data is fitted well at a range of loss function exponents, as the bias is dominated by prior attraction. A flat prior as assumed by Jazayeri and Shadlen [15], Remington et al. [25] can only fit the data with a higher exponent.

###### S2.3.3.3 Time Interval Estimation (Remington et al. [25])

**Details** For implementation, we chose $\mathcal{X} = [0, 3]$, and used 200 grid points. The upper end of the interval was chosen in order to avoid boundary effects, as the "true" stimulus space is unbounded.

Model fit statistics are shown in Figure S35. Unimodal priors, normal in stimulus or sensory space, achieve near-optimal fit.

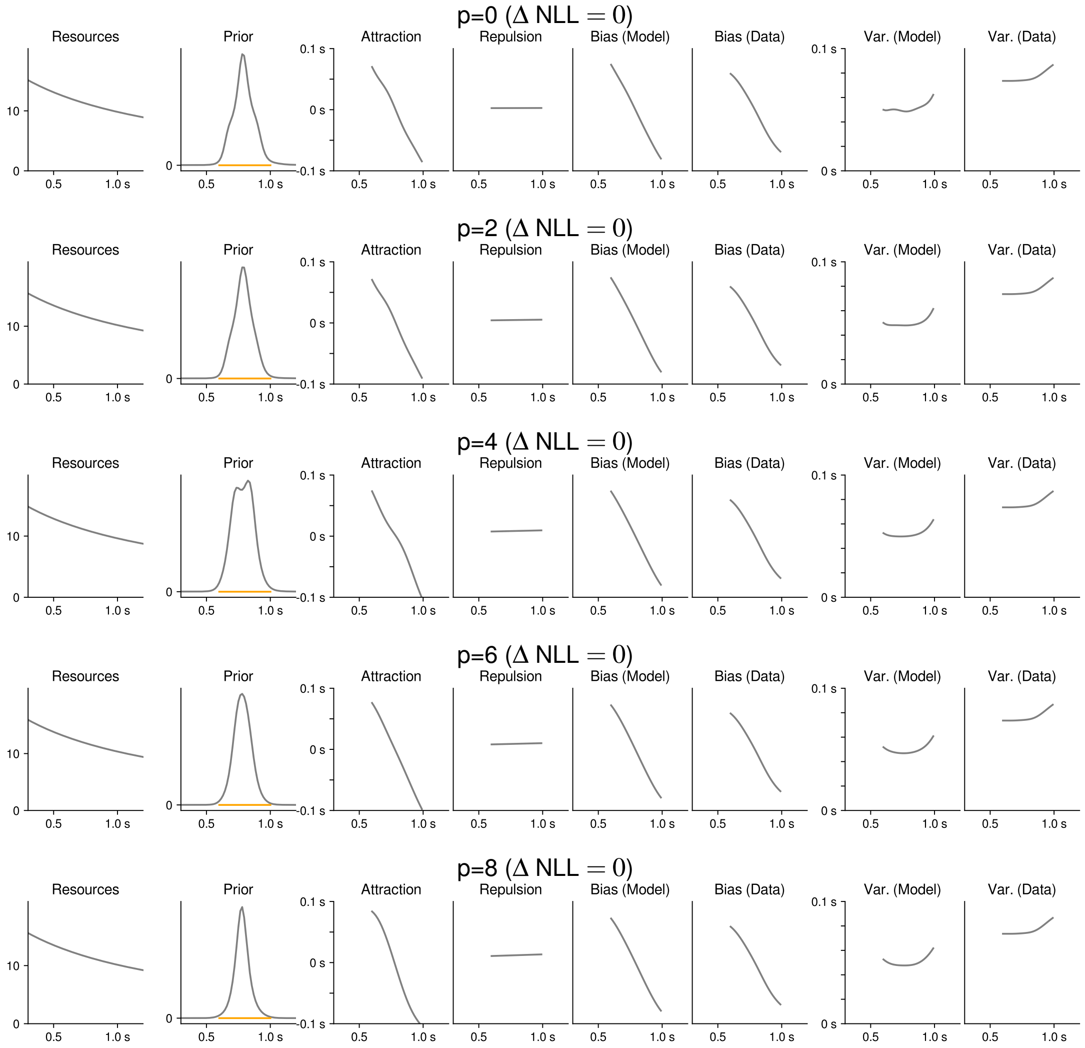

Figure S36: Time interval estimation [data collected by 25]: Freely fitted prior for time interval estimation, as a function of the loss function exponent. The orange bar indicates the range of stimuli in the experiment. All exponents achieve similar quantitative model fit (see Figure S35).

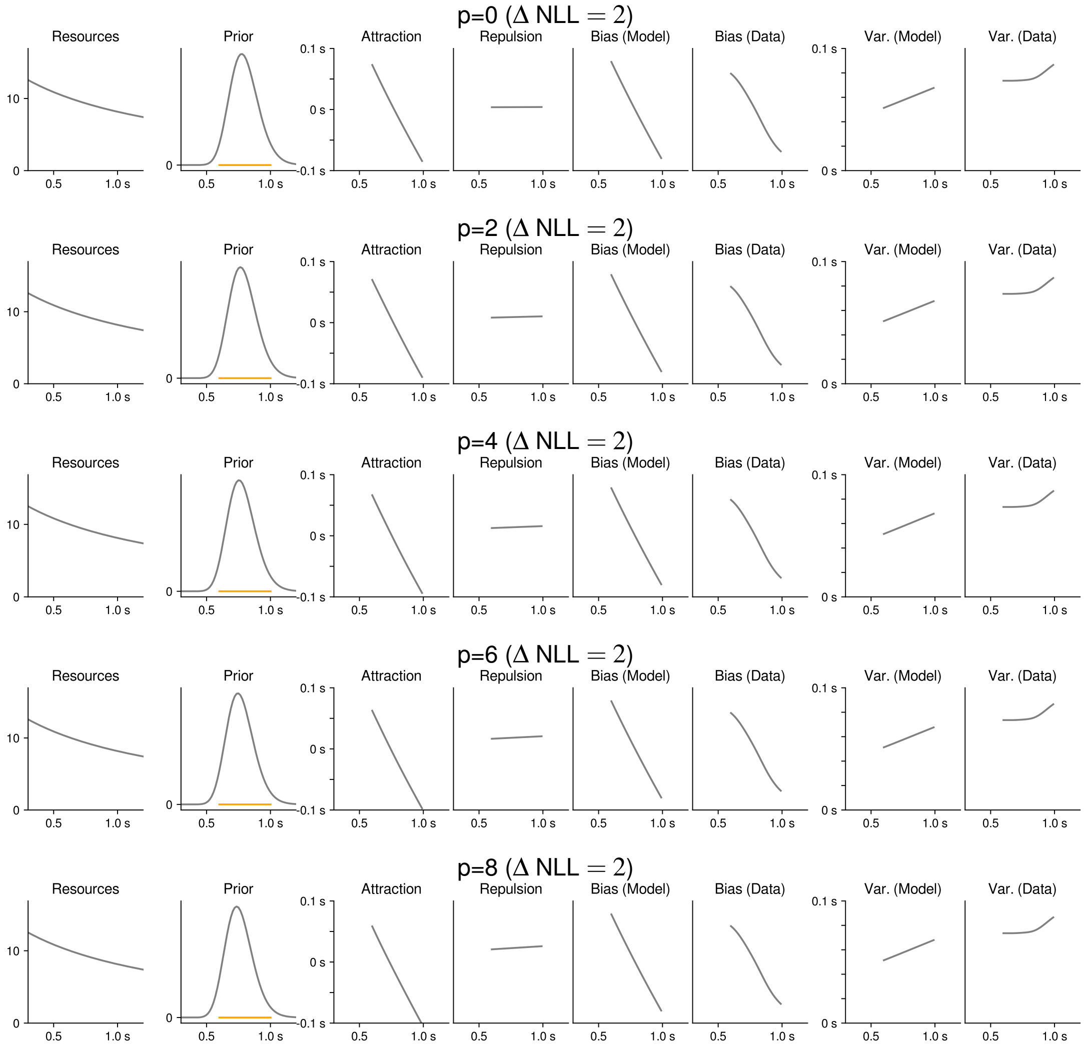

Figure S37: Time interval estimation [data collected by 25]: Parametric unimodal prior (normal in sensory space).

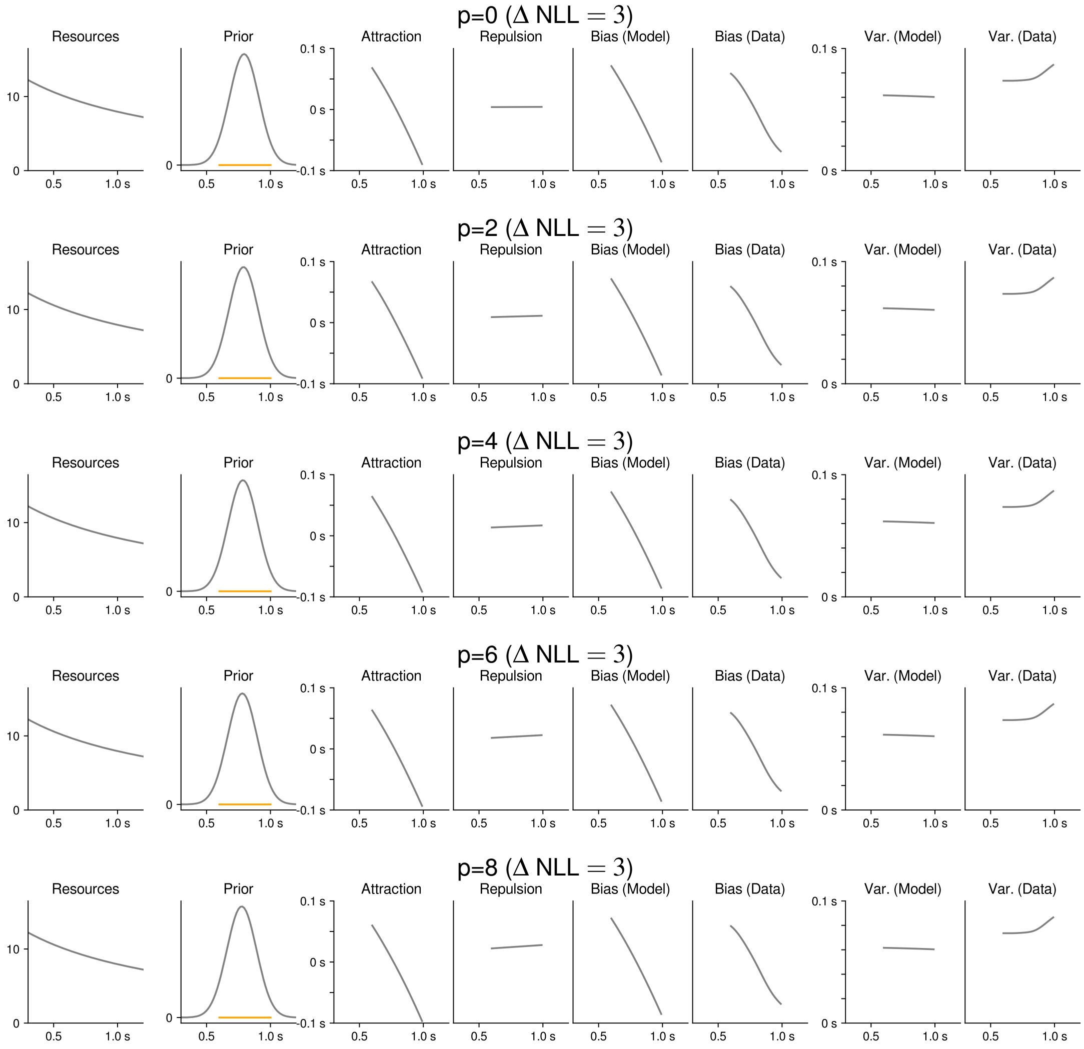

Figure S38: Time interval estimation [data collected by 25]: Parametric unimodal prior (normal in stimulus space).

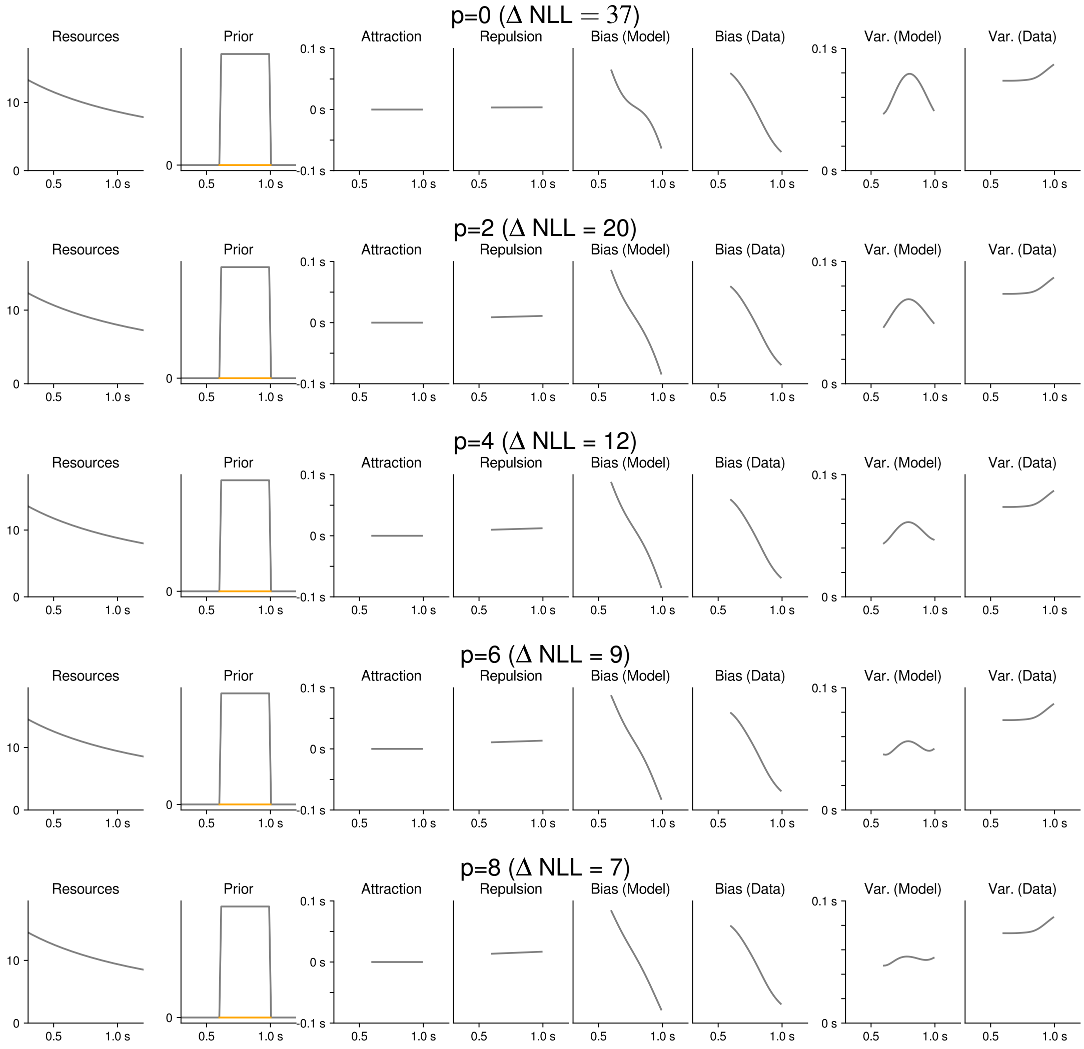

Figure S39: Time interval estimation [data collected by 25]: Flat prior

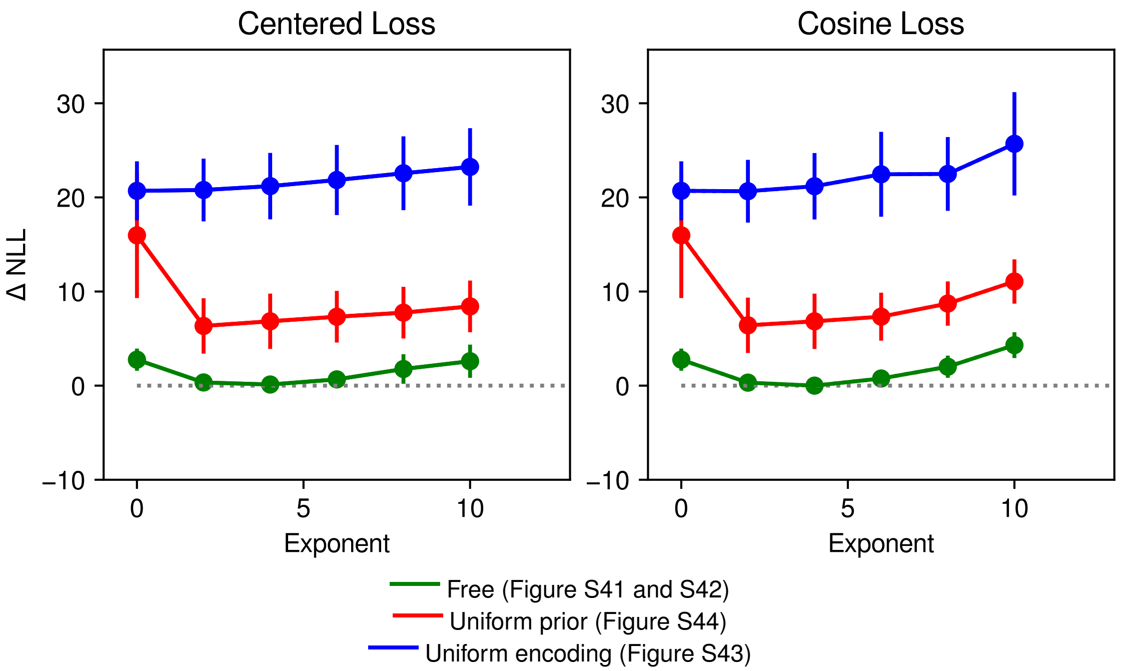

Figure S40: Color perception [data collected by 1]: Negative Log-Likelihood (y-axis) relative to the model plotted in the main paper as a function of the loss function exponent, averaged across the 10 folds, for two implementations of $L^p$ loss: Left: using the ordinary $L^p$ distance after centering the space at $F^{-1}(m)$. Right: using cosine-based loss. The model with freely fitted prior and encoding consistently improves over models where one is constrained to be uniform, particularly a model with uniform encoding. Error bars denote standard errors after regressing out variability between the 10 folds.

###### S2.3.4 Color Perception (Bae et al. [1])

**Results** Model fit statistics are shown in Figure S40, with references to further results. In addition to the data from 7 subjects analyzed in the main paper, Bae et al. [1] also collected data from 3 subjects where there was a delay after the stimulus. In Figure S45, we pooled the data from these different experiments and we fitted the version of the model including per-subject adjustments to the pooled data. The qualitative pattern of encoding, prior, bias, and variability are fitted across subjects; biases and variability are increased in the delayed condition, as memory load increases sensory noise.

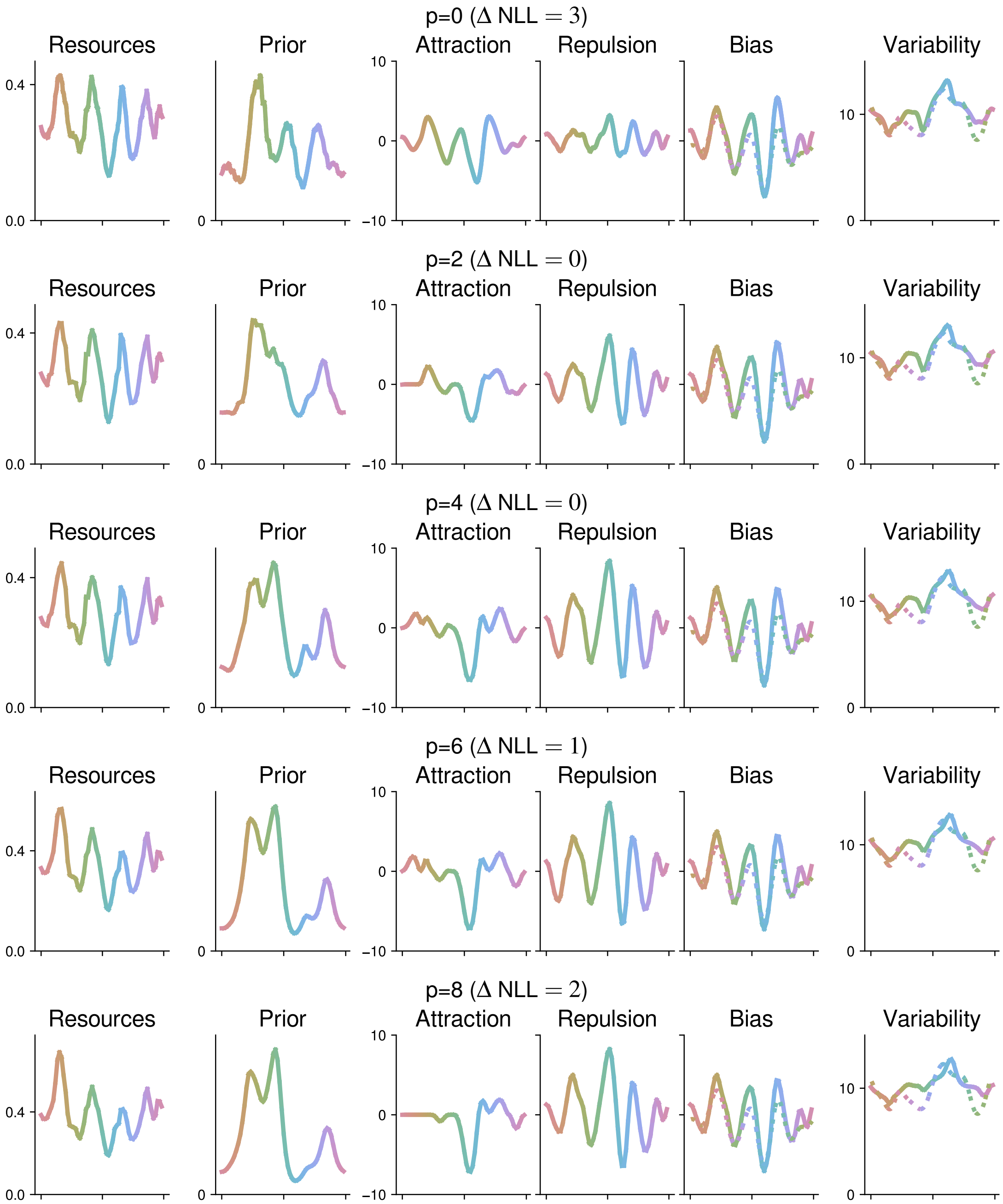

Figure S41: Perception of color hue [data collected by 1]: Fit as a function of the loss function exponent (using cosine loss implementation). Across exponents $p > 0$, the biases are explained in terms of a periodic resource allocation and an attraction effect accounting for unusally negative biases in the blue range. At p=0, prior and encoding both exhibit periodic patterns; however, unlike in categories-as-priors accounts, the prior peaks at category boundaries: We show distributions for color categories as measured by Bae et al. [1] in an independent experiment.

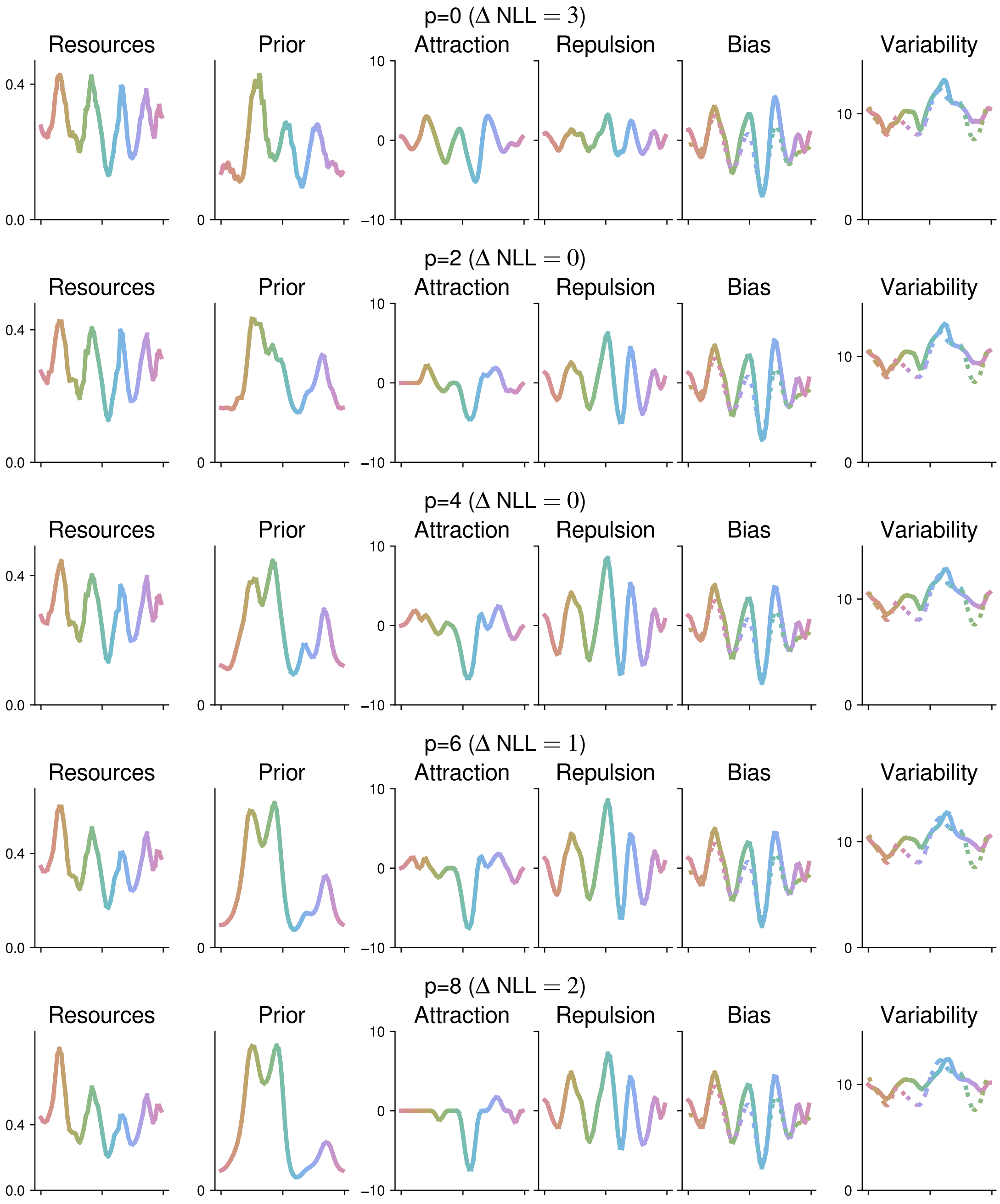

Figure S42: Perception of color hue [data collected by 1]: Perception of color hue: Fit as a function of the loss function exponent (using centered implementation of loss function, rather than cosine-based implementation).

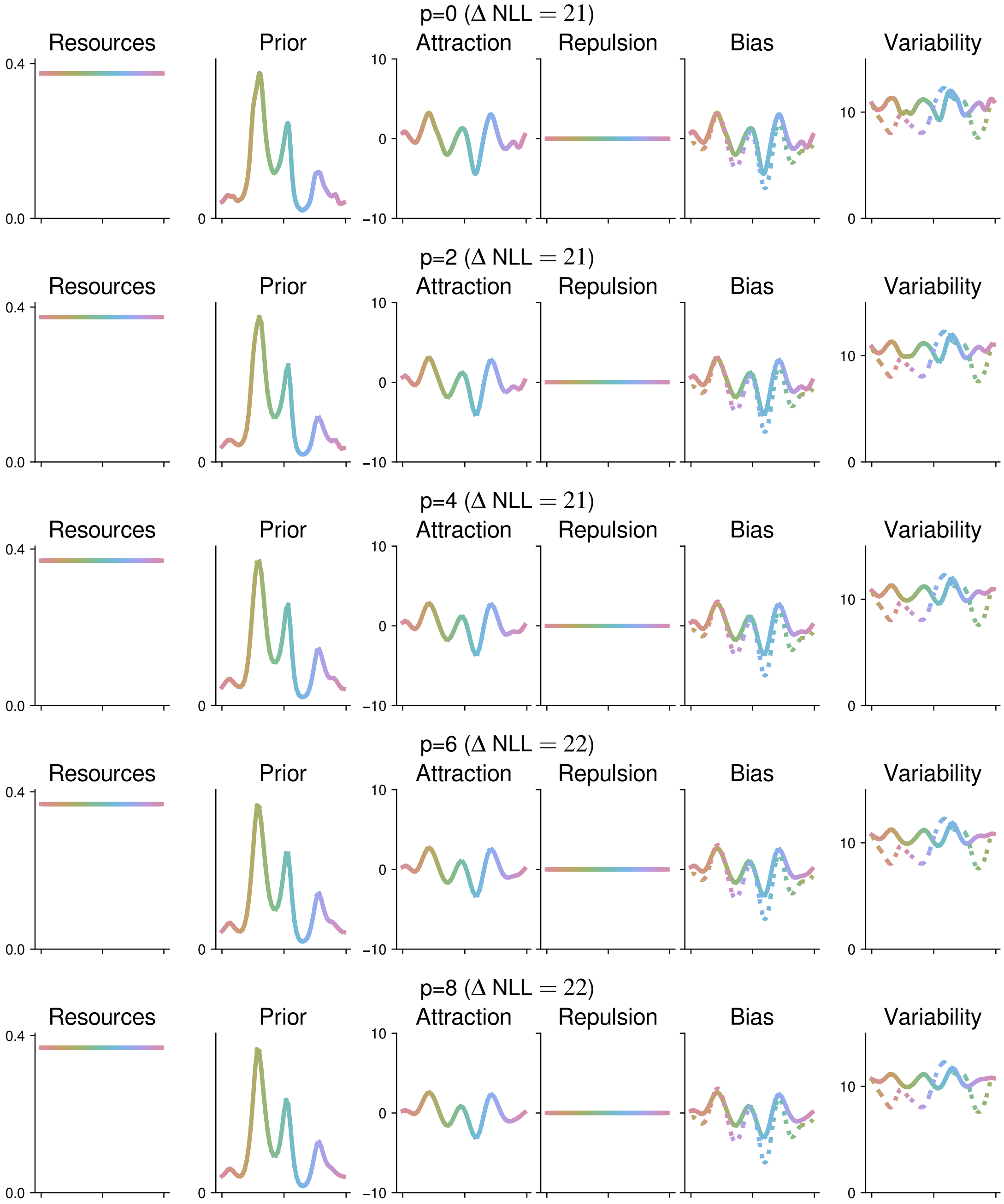

Figure S43: Perception of color hue [data collected by 1]: Uniform encoding, cosine-based implementation of loss function.

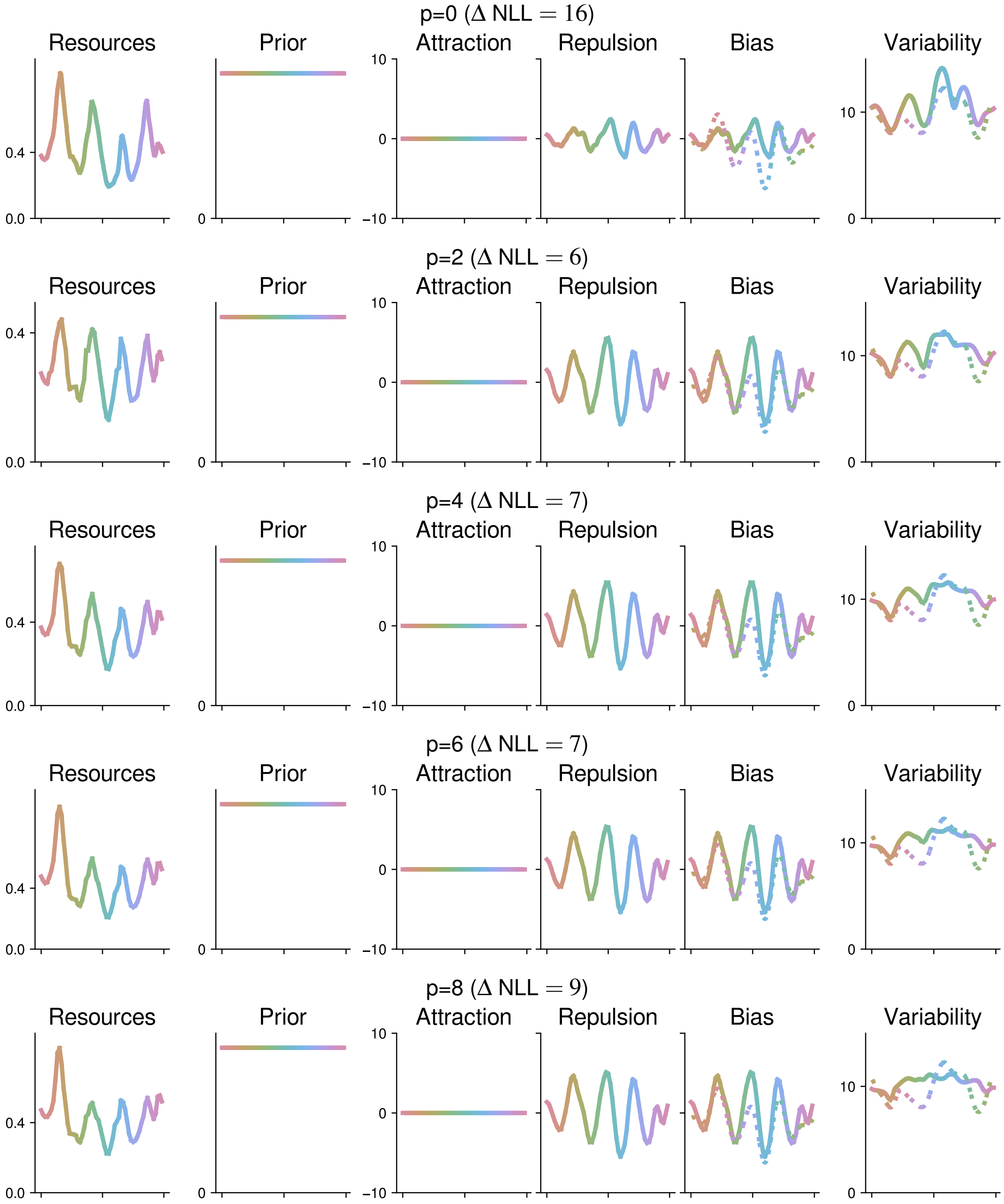

Figure S44: Perception of color hue [data collected by 1]: Uniform prior, cosine-based implementation of loss function.

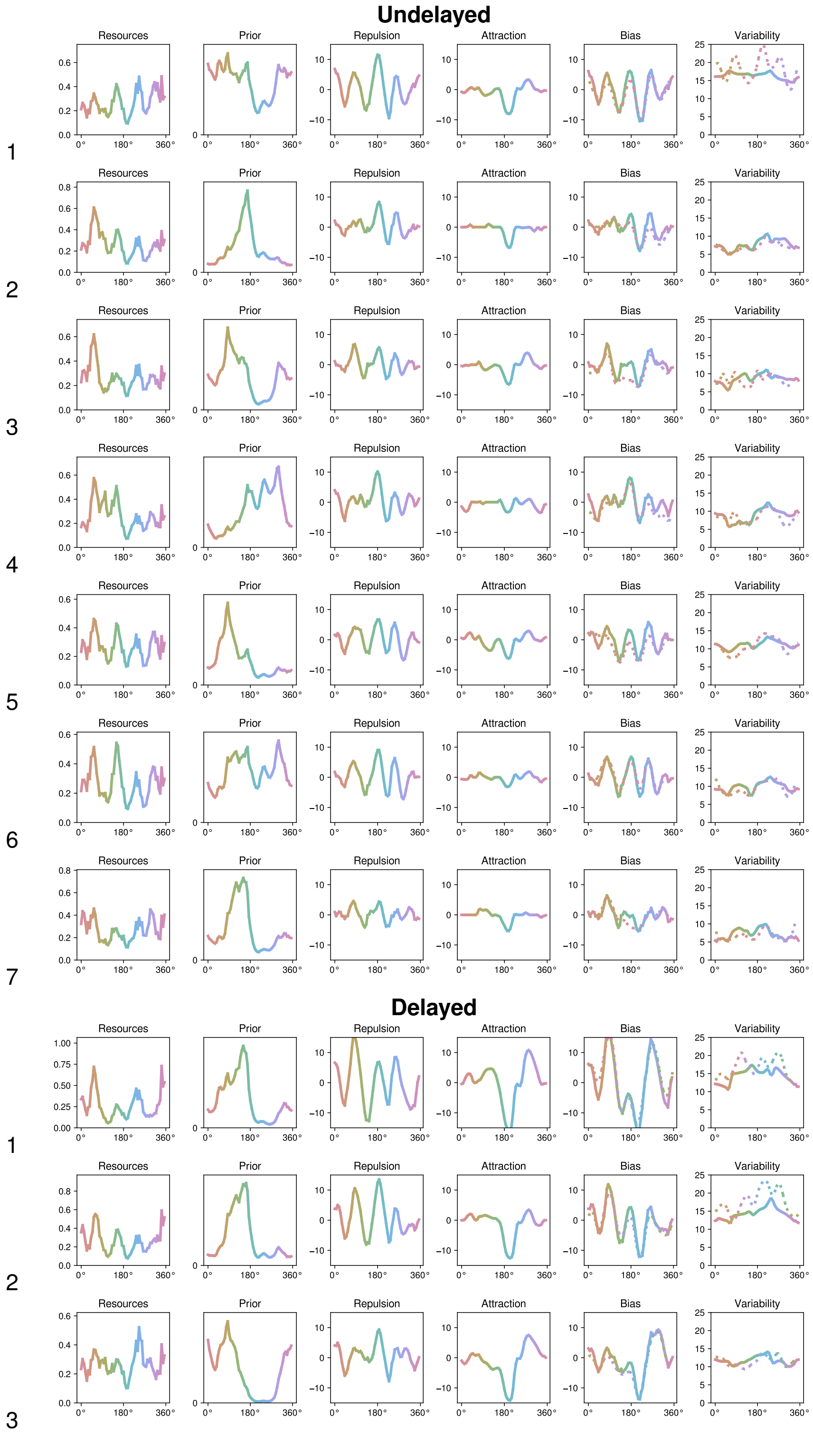

Figure S45: Perception of color hue [data collected by 1]: Per-subject fit of hierarchical model with subject-specific adjustments to FI, prior, and noise parameters; using loss function exponent p=4. In addition to the 7 subjects from the experiment fitted in the Main Text, we additionally included three further subjects where the stimulus was followed by a delay (N=3, bottom). Across subjects, the model infers the same periodic structure in resource allocation, and a prior consistently producing negative prior attraction at ≈200°.

### S3 Formal Proofs

#### S3.1 Proof of Theorem 1

##### S3.1.1 Preliminaries

**Formal Assumptions** We first recapitulate the formal assumptions stated in the main paper. We assume that the stimulus space $\mathcal{X}$ is a contiguous one-dimensional manifold (the real line, an interval, or a circle). The sensory space $\mathcal{Y}$ has the same topology as the stimulus space; it conceptually represents the one-dimensional submanifold of the space of neural activations spanned by the mean encodings of the stimuli in $\mathcal{X}$. We assume that $F : \mathcal{X} \to \mathcal{Y}$ is bijective and differentiable, such that its slope (and thus the Fisher information of the encoding) is nowhere zero. Thus, any two stimuli receive distinct encodings when noise is small enough (no two stimuli are mapped to the same output). We assume that the prior is nowhere zero. Further, we make basic regularity assumptions: $F'(\theta)$ and $\log p_{prior}(\theta)$ are twice continuously differentiable, and both they and their first and second derivatives cannot grow super-polynomially in $\theta$. We relax these assumptions for Theorem 3 (Boundary Effects) by allowing the prior to discontinuously become zero at a boundary $\theta_{Max}$.

These assumptions are very broadly satisfied. Bijectivity of $F$ is a prerequisite for any two stimuli being distinguishable when noise is sufficiently small. Differentiability of encoding and prior is also broadly satisfied.[16] Polynomially bounded growth is satisfied by power-law functions (such as encoding following Weber's law) and the log-density of common probability distributions such as Gaussian or lognormal distributions. It is automatically satisfied when $\mathcal{X}$ is a circle or a closed finite interval.

**Encoding Model** Recall that the encoding model is given as

$$m = F(\theta) + \delta \tag{39}$$

where $F$ is a transformation from stimulus space to sensory space, $m$ is a point in the sensory space, and $\delta$ is mean-zero noise, Gaussian with variance $\sigma^2$ or von Mises with $\kappa = \frac{1}{\sigma^2}$. Throughout, we will write $t$ for the sensory variance $\sigma^2$ to make notation simpler.

**Reduction to Unbounded Space** Throughout all proofs in this SI Appendix, we will reduce computation of biases to the special case where $\mathcal{X}$ is the full real line, by noting that, when noise is small, only exponentially small probability mass falls into regions outside a local environment of $\theta_0$. Hence, when encoding and prior obey basic regularity conditions, all relevant integrals can be computed equivalently on the real line or $\mathcal{X}$, up to higher-order error in $\sigma$. We describe this here at the example of computing an expectation involving the prior. Let $k \geq 0$ be any integer, and let $\theta_0$ be any point in the interior of an interval $\mathcal{X}$, then there is some $C_{\theta_0} > 0$ such that $B_{C_{\theta_0}}(\theta_0) \subseteq \mathcal{X}$, and

$$p_{prior}(\theta) = p_{prior}(\theta_0) + (\theta - \theta_0)\frac{\mathrm{d}}{\mathrm{d}\theta}p_{prior}(\theta_0) + O((\theta-\theta_0)^2) \tag{40}$$

for all $\theta$ such that $|\theta - \theta_0| < C_{\theta_0}$. Then, if $P$ is the likelihood,

$$\begin{aligned}
\int_{\mathcal{X}} \theta^k p_{prior}(\theta) dP(\theta) =& p_{prior}(\theta_0)\int_{\mathcal{X}} 1_{|\theta-\theta_0|<C_{\theta_0}} \theta^k dP(\theta) \\
&+ \frac{\mathrm{d}}{\mathrm{d}\theta}p_{prior}(\theta_0)\int_{\mathcal{X}} 1_{|\theta-\theta_0|<C_{\theta_0}} \theta^k (\theta-\theta_0) dP(\theta) \\
&+ \int_{\mathcal{X}} 1_{|\theta-\theta_0|<C_{\theta_0}} \theta^k O((\theta-\theta_0)^2) dP(\theta) \\
&+ \int_{\mathcal{X}} 1_{|\theta-\theta_0|>C_{\theta_0}} \theta^k [p_{prior}(\theta_0) + (\theta-\theta_0)\frac{\mathrm{d}}{\mathrm{d}\theta}p_{prior}(\theta_0) + O((\theta-\theta_0)^2)] dP(\theta) \\
&+ \int_{\mathcal{X}} 1_{|\theta-\theta_0|>C_{\theta_0}} \theta^k [p_{prior}(\theta) - [p_{prior}(\theta_0) + (\theta-\theta_0)\frac{\mathrm{d}}{\mathrm{d}\theta}p_{prior}(\theta_0) + O((\theta-\theta_0)^2)]] dP(\theta)
\end{aligned}$$

[16] An exception is the prior $2 - |\sin(\alpha)|$ assumed for orientation perception, but this can be approximated very closely with smooth functions.

Because the prior is assumed to have polynomially bounded growth, the last two integrals are bounded as $\int_{\mathbb{R}} 1_{|\theta-\theta_0|>C_{\theta_0}} |\theta|^q dP(\theta)$ for some nnonegative integer $q$, which is $o(\sigma^l)$ for any positive integer $l$ as $\sigma \to 0$; this remains true when the integration domains $\mathcal{X}$ are changed to $\mathbb{R}$. Thus, the integral can be evaluated up to higher-order error

$$\int_{\mathcal{X}} \theta^k p_{prior}(\theta) dP(\theta) = \int_{\mathbb{R}} \theta^k [p_{prior}(\theta_0) + (\theta-\theta_0)\frac{\mathrm{d}}{\mathrm{d}\theta} p_{prior}(\theta_0) + O((\theta-\theta_0)^2)] dP(\theta) + o(\sigma^l) \tag{41}$$

for any positive integer $l$. When $\mathcal{X}$ is not an interval but instead circular, behavior can be reduced to considering an interval around $\theta_0$ by the same arguments, because the von Mises distribution behaves like a Gaussian with variance $\frac{1}{\kappa}$ in the limit $\kappa \to \infty$.

**Likelihood and Posterior** The likelihood has the form

$$p(m|\theta) = \frac{1}{\sqrt{2\pi t}} \exp\left(-\frac{(m-F(\theta))^2}{2t}\right) \tag{42}$$

While the likelihood is Gaussian, mapping it to the stimulus space via $F^{-1}$ in general results in a non-Gaussian distribution, unless $F$ is linear. The posterior has the density

$$p(\theta|m) = \frac{p_{lik}(m|\theta) p_{prior}(\theta)}{\int p_{lik}(m|\theta) p_{prior}(\theta) d\theta} \propto \exp\left(-\frac{(m-F(\theta))^2}{2t}\right) p_{prior}(\theta) \tag{43}$$

Recall that the Fisher information of the model (Eq. 39) is given by the slope of the encoding and the sensory variance as follows:

$$\mathcal{J} = \frac{\mathcal{S}}{\sigma^2} = \frac{\mathcal{S}}{t} := \frac{(\frac{\mathrm{d}}{\mathrm{d}\theta} F)^2}{t} \tag{44}$$

**Statement of Theorem** The bias of the estimator $\widehat{\theta}$ is given as

$$\mathbb{E}[\widehat{\theta}] - \theta \tag{45}$$

where $\theta$ is fixed, and $\widehat{\theta}$ is a random variable that depends (deterministically) on the encoding $m$. Theorem 1 states:

**Theorem S1.** *Let $\theta$ be a point in the interior of $\mathcal{X}$. For the model defined in Eq. (1), the bias for the estimator $\widehat{\theta}$ for $L^p$ loss ($p \geq 1$ an integer) with arbitrary prior can be written as*

$$\mathbb{E}[\widehat{\theta}] - \theta = \underbrace{\frac{1}{\mathcal{J}} (\log p_{prior})'}_{\text{Prior Attraction}} + \underbrace{\frac{p+2}{4} \left(\frac{1}{\mathcal{J}}\right)'}_{\text{Likelihood Repulsion}} ; \tag{46}$$

*and, for the MAP estimator ($p \to 0$),*

$$\mathbb{E}[\widehat{\theta}] - \theta = \frac{1}{\mathcal{J}} (\log p_{prior})' + \frac{1}{4} \left(\frac{1}{\mathcal{J}}\right)', \tag{47}$$

*both up to approximation error $O(\sigma^4 \cdot C_{F,p_{prior},p,\theta})$, when $\sigma^2$ is sufficiently close to $0$, where the error contains constants depending on $F, p_{prior}, p, \theta$, but not $\sigma$. Here $\mathcal{J} = \frac{(F'(\theta))^2}{\sigma^2}$ is the Fisher information; both $\mathcal{J}$ and $p_{prior}$ are functions of $\theta$. All quantities are evaluated at $\theta$.*

**Encoding Bias and Decoding Bias** Given a stimulus $\theta$, the bias of an estimator $\widehat{\theta}$ can be decomposed as

$$\mathbb{E}[\widehat{\theta}] - \theta = \underbrace{\mathbb{E}[F^{-1}(m)] - \theta}_{\text{Encoding Bias}} + \underbrace{\mathbb{E}[\widehat{\theta} - F^{-1}(m)]}_{\text{Decoding Bias}} \tag{48}$$

where the expectation is over the encoding $m$; $\theta$ is taken to be constant. We will separately treat the two components. Throughout, we will compute $\mathbb{E}[\widehat{\theta} - F^{-1}(m)]$ in terms of $\mathcal{J}$ and $p_{prior}$ and their derivatives all evaluated at $F^{-1}(m)$, and then transfer this to an expression in terms of these quantities evaluated at $\theta$ using Eq. (71).

###### S3.1.2 Special Cases

Table S1 lists and discusses analytical results from prior work that fall out of Theorem 1 as special cases. Most of these are immediate consequences of Theorem 1. Here, we discuss the derivation of (d). Morais and Pillow [19] proposed a model family with power-law efficient coding ((d) in Table S1), i.e. $\mathcal{J} \propto p_{prior}^q$ or $p_{prior} \propto \mathcal{J}^{1/q}$. However, they did not derive a general expression for the bias beyond $p = 0$; the special case $p = 2$ was later found by Prat-Carrabin and Woodford [24]. Using Theorem 1, we can derive the general bias across loss function exponents $p$ and powers $q$ (here $p > 0$, analogously at $p = 0$):

$$\begin{aligned} &\frac{1}{\mathcal{J}}(\log p_{prior})' + \frac{p+2}{4}\left(\frac{1}{\mathcal{J}}\right)' \\ =&\frac{1}{q}\frac{\mathcal{J}'}{\mathcal{J}^2} + \frac{p+2}{4}\left(\frac{1}{\mathcal{J}}\right)' \\ =&\left(\frac{p+2}{4} - \frac{1}{q}\right)\left(\frac{1}{\mathcal{J}}\right)' \end{aligned}$$

The flip between attraction and repulsion occurs at

$$q = \frac{4}{p+2} \tag{49}$$

if $p > 0$.[17] These results recover special cases for $q = 2$ [32] ((c) in Table S1) and $p = 2$ [24].

In the Gaussian model originally studied by Morais and Pillow [19], our results in Section S3.5 analogously yield a general expression for the bias:

$$\frac{1}{\mathcal{J}}(\log p_{prior})' + \frac{p}{2}\left(\frac{1}{\mathcal{J}}\right)' = \left(\frac{p}{2} - \frac{1}{q}\right)\left(\frac{1}{\mathcal{J}}\right)' \tag{50}$$

The flip between attraction and repulsion occurs at

$$q = \frac{2}{p} \tag{51}$$

This correctly accounts for the location of the zero-crossing that they observed in numerical simulations [19, Figure 5] at $p = 0$ ($q = \infty$), $p = 1$ ($q = 2$), $p = 2$ ($q = 1$).

###### S3.1.3 Encoding Bias

Encoding bias is defined as the first term on the right-hand side of Eq. (48):

$$\mathbb{E}[F^{-1}(m)] - \theta \tag{52}$$

This was already solved by [31]; we recapitulate this argument here for completeness. As we aim to compute biases rather than absolute stimulus positions, only the relative positions of stimuli matter; we can therefore assume without loss of generality $\theta = 0$ and $F(\theta) = 0$. Then, the encoding bias is $\mathbb{E}[F^{-1}(m)|\theta]$. Now noting that the likelihood has the form

$$p(m|\theta) = \frac{1}{\sqrt{2\pi\sigma^2}}\exp\left(-\frac{(m - F(\theta))^2}{2\sigma^2}\right) = \frac{1}{\sqrt{2\pi\sigma^2}}\exp\left(-\frac{m^2}{2\sigma^2}\right) \tag{53}$$

and applying Taylor's theorem to $F^{-1}$, the encoding bias $\mathbb{E}[F^{-1}(m)|\theta]$ equals:

$$\int \frac{1}{\sqrt{2\pi t}}\left(\frac{1}{2}x^2\frac{\mathrm{d}^2}{\mathrm{d}x^2}F^{-1} + O(x^4)\right)\exp\left(-\frac{x^2}{2\sigma^2}\right)dt = \frac{\sigma^2}{2}\frac{\mathrm{d}^2}{\mathrm{d}^2}F^{-1} + O(\sigma^4) = \sigma^2\frac{1}{4}\frac{\mathrm{d}}{\mathrm{d}\theta}\frac{1}{\mathcal{S}} + O(\sigma^4) \tag{54}$$

Here, we have suppressed constants depending on $F, p_{prior}, \theta$ in the $O(\cdot)$ terms; we will do this throughout all derivations here in order to simplify notation.

[17] If $p = 0$, analogous derivation using Theorem 1 leads to $q = 4$.

| | Encoding | Loss | Bias | References |
|---|---|---|---|---|
| (a) | Arbitrary | $p=0$ | $\frac{1}{\mathcal{J}}(\log p_{prior})' + \frac{1}{4}\left(\frac{1}{\mathcal{J}}\right)'$ | [28] |
| (b) | Arbitrary | $p=2$ | $\frac{1}{\mathcal{J}}(\log p_{prior})' + \left(\frac{1}{\mathcal{J}}\right)'$ | [24] |
| (c) | $\mathcal{J} \propto p_{prior}^2$ | $p>0$ | $\frac{p}{4}\left(\frac{1}{\mathcal{J}}\right)'$ | [32] |
| (d) | $\mathcal{J} \propto p_{prior}^q$ | $p>0$ | $\left[\frac{p+2}{4} - \frac{1}{q}\right]\left(\frac{1}{\mathcal{J}}\right)'$ | $p=2$: [24] |
| (e) | Uniform | any $p$ | $\sigma^2 (\log p_{prior})'$ | e.g., [14] |

Table S1: Special cases of Theorem 1 recover analytical results from prior work. We note that the precise factors associated with likelihood repulsion depend systematically on the encoding model and are accordingly different in some previous studies; see Section S3.5 for more on this. (a-b) The exponents $p=0,2$ correspond to the posterior mode and mean, respectively; prior work found expressions for the bias at $p=0$ [28] and $p=2$ [24]. (c) Maximizing the mutual information between the stimulus and the neural response requires the square root of the FI to be proportional to the prior distribution. This setting was analyzed by Wei and Stocker [32] (here $p>0$). (d) Morais and Pillow [19] proposed a broader family of encoding models where the Fisher information is proportional to powers of the prior. Bias (here $p>0$) flips from repulsive to attractive when the coefficient crosses zero; the bias was found in the special case $p=2$ by Prat-Carrabin and Woodford [24]. See Section S3.1.2 for derivation of the general expression. (e) In the limit $q \to 0$, the encoding becomes uniform; this setting is often assumed in traditional Bayesian models of perceptual biases. Here, only prior attraction remains, independent of the loss function.

##### S3.1.4 Decoding Bias

We compute the decoding bias separately for even exponents $L^{2p}$ ($p>0$), for odd exponents $L^{2p+1}$, and for $p=1,0$.

**S3.1.4.1 Positive even exponent: $L^{2q}$ loss, $q>0$** Previous work computed this bias in the special case where $\mathcal{J} \propto p_{prior}$ [32] or where the exponent equals 2 [24]; we will compute it for arbitrary combinations of encocodings, priors, and loss function exponents. We begin by computing the Bayes estimator $\widehat{\theta}$ given a fixed encoding $m$. Throughout, we will write the exponent as $p=2q$. The Bayes estimator is defined as

$$\widehat{\theta} := \arg_{\widehat{\theta}} \min \int |\widehat{\theta} - \theta|^{2q} P(\theta|m) d\theta \tag{55}$$

Hence, we aim to solve the following equation for the estimator $\widehat{\theta}$:

$$0 = \frac{d}{d\widehat{\theta}} \int (\widehat{\theta} - \theta)^{2q} P(\theta|m) d\theta \tag{56}$$

$$= \frac{d}{d\widehat{\theta}} \int (\widehat{\theta} - \theta)^{2q} p_{prior}(\theta) P(m|\theta) d\theta \tag{57}$$

$$= \frac{d}{d\widehat{\theta}} \int (\widehat{\theta} - \theta)^{2q} p_{prior}(\theta) \frac{1}{\sqrt{2\pi\sigma^2}} \exp\left(-\frac{|F(\theta)-m|^2}{2\sigma^2}\right) d\theta \tag{58}$$

$$= 2q \int (\widehat{\theta} - \theta)^{2q-1} p_{prior}(\theta) \frac{1}{\sqrt{2\pi\sigma^2}} \exp\left(-\frac{|F(\theta)-m|^2}{2\sigma^2}\right) d\theta \tag{59}$$

Defining

$$q_{prior}(m) := \frac{p_{prior}(m)}{\sqrt{\mathcal{S}(F^{-1}(m))}}$$

as a function in sensory space, we write[18]

$$q_{prior}(x) = q_{prior}(m) + x \frac{d}{dm} q_{prior}(m) + O(x^2)$$

[18] It can be verified that the suppressed higher-order terms play no role in the lowest-order terms identified below.

Note that

$$\frac{\mathrm{d}}{\mathrm{d}m} = \frac{1}{\sqrt{\mathcal{S}}}\frac{\mathrm{d}}{\mathrm{d}\theta} \tag{60}$$

By shifting stimulus space and sensory space, we can assume without loss of generality that $m = 0$ and that $F(0) = 0$, without affecting the solution of Eq. (56). We then transform Eq. (56) into an integral in sensory space, reparameterizing $r := \widehat{\theta}$, $x := F(\theta)$:

$$0 = \text{Eq. (59)} \propto \int (r - F^{-1}(x))^{2q-1} q_{prior}(x) \frac{1}{\sqrt{2\pi\sigma^2}} \exp\left(-\frac{x^2}{2\sigma^2}\right) dx \tag{61}$$

Then we express

$$\theta = F^{-1}(m) = m + x \cdot \left(\frac{\mathrm{d}}{\mathrm{d}m} F^{-1}(m)\right) + \frac{1}{2} x^2 \cdot \left(\frac{\mathrm{d}^2}{\mathrm{d}m^2} F^{-1}(m)\right) + O(x^3)$$

Writing

$$(-A) := \frac{\mathrm{d}}{\mathrm{d}x} F^{-1} = \frac{1}{\sqrt{\mathcal{S}}} \tag{62}$$

$$(-B) := \frac{1}{2}\frac{\mathrm{d}^2}{\mathrm{d}x^2} F^{-1} = \frac{1}{2}\frac{\mathrm{d}}{\mathrm{d}x}\left(\frac{1}{\sqrt{\mathcal{S}}}\right) = \frac{1}{2}\frac{1}{\sqrt{\mathcal{S}}}\frac{\mathrm{d}}{\mathrm{d}\theta}\left(\frac{1}{\sqrt{\mathcal{S}}}\right) = \frac{1}{4}\frac{\mathrm{d}}{\mathrm{d}\theta}\frac{1}{\mathcal{S}} \tag{63}$$

Substituting these terms and suppressing the higher-order terms for conciseness, and writing $t := \sigma^2$ to make later formulas easier to read, we obtain from Eq. (61):

$$0 = \int \left(r + Ax + Bx^2\right)^{2q-1} \frac{1}{\sqrt{t}} \left(q_{prior}(m) + x \frac{\mathrm{d}}{\mathrm{d}m} q_{prior}(m)\right) \exp\left(-\frac{x^2}{2t}\right) \tag{64}$$

and hence due to Eq. 60:

$$\frac{\mathrm{d}}{\mathrm{d}m} \log q_{prior} = \frac{\mathrm{d}}{\mathrm{d}m} \log \frac{q_{prior}}{\sqrt{\mathcal{S}}} = \frac{1}{\sqrt{\mathcal{S}}}\left(\frac{\mathrm{d}}{\mathrm{d}\theta} \log \frac{q_{prior}}{\sqrt{\mathcal{S}}}\right) \tag{65}$$

What is the scaling of the solution $r$ as $t \to 0$? Presume $r = \omega(t)$ (e.g. $r \sim \sqrt{t}$) as $t \to 0$. Expanding Eq. (64), we get

$$\begin{aligned}
0 &= \int (r + Ax + Bx^2)^{2q-1} \frac{1}{\sqrt{t}} \left(q_{prior}(m) + x \frac{\mathrm{d}}{\mathrm{d}m} q_{prior}(m)\right) \exp\left(-\frac{x^2}{2t}\right) dx \\
&= \int \left(\sum_{n_1+n_2+n_3=2q-1} \binom{2q-1}{n_1 n_2 n_3} r^{n_1} A^{n_2} x^{n_2+2n_3} B^{n_3}\right) \frac{1}{\sqrt{t}} \left(q_{prior}(m) + x \frac{\mathrm{d}}{\mathrm{d}m} q_{prior}(m)\right) \exp\left(-\frac{x^2}{2t}\right) dx \\
&= \sum_{n_1+n_2+n_3=2q-1} \binom{2q-1}{n_1 n_2 n_3} r^{n_1} A^{n_2} B^{n_3} q_{prior}(m) \int x^{n_2+2n_3} \frac{1}{\sqrt{t}} \exp\left(-\frac{x^2}{2t}\right) dx \\
&\quad + \sum_{n_1+n_2+n_3=2q-1} \binom{2q-1}{n_1 n_2 n_3} r^{n_1} A^{n_2} B^{n_3} \frac{\mathrm{d}}{\mathrm{d}m} q_{prior}(m) \int x^{n_2+2n_3+1} \frac{1}{\sqrt{t}} \exp\left(-\frac{x^2}{2t}\right) dx
\end{aligned}$$

If $r = \omega(\sqrt{t})$, the lowest-order terms are those where $n_1 = 2q-1$ (for the $q_{prior}(m)$ part) or $2q-2$ (for the $\frac{\mathrm{d}}{\mathrm{d}m} q_{prior}(m)$ part) (note that a single power of $x$ already contributes a half power of $t$ under the Gaussian integral, and thus more than $r$):

$$r^{2q-1} q_{prior}(m) + (2q-1) r^{2q-2} A \left(\frac{\mathrm{d}}{\mathrm{d}m} q_{prior}(m)\right) t$$

In particular, the lowest order term is $r^{2q-1}q_{prior}(m)$ which is not zero if $r=\omega(t)$. This is a contradiction to the claim that the whole expression equals zero. If on the other hand the scaling of $r$ is between $\sqrt{t}$ and $t$, the lowest-order terms are those where $n_1=1$ for the $q_{prior}(m)$ part and $n_1=0$ for the $\frac{\mathrm{d}}{\mathrm{d}m}q_{prior}(m)$ part, and $n_2=2q-1-n_1$:

$$(2q-1)rA^{2q-2}q_{prior}(m)t^{q-1}+A^{2q-1}\left(\frac{\mathrm{d}}{\mathrm{d}m}q_{prior}(m)\right)t^q$$

Here, again the left term has lower order, and is thus the lowest-order term of the whole expression. Again, it is manifestly not zero, giving a contradiction to the claim that the whole expression can equal zero. Hence $r=O(t)$. Having shown this, we can write $r=tR+O(t^2)$, and obtain – suppressing higher-order terms for notational simplicity:

$$\begin{aligned}
0&=\int(tR+Ax+Bx^2)^{2q-1}\frac{1}{\sqrt{t}}(q_{prior}(m)+x\frac{\mathrm{d}}{\mathrm{d}m}q_{prior}(m))\exp\left(-\frac{x^2}{2t}\right)dx\\
&=\int\left(\sum_{n_1+n_2+n_3=2q-1}\binom{2q-1}{n_1n_2n_3}(tR)^{n_1}A^{n_2}x^{n_2+2n_3}B^{n_3}\right)\frac{1}{\sqrt{t}}(q_{prior}(m)+x\frac{\mathrm{d}}{\mathrm{d}m}q_{prior}(m))\exp\left(-\frac{x^2}{2t}\right)dx\\
&=\sum_{n_1+n_2+n_3=2q-1}\binom{2q-1}{n_1n_2n_3}(tR)^{n_1}A^{n_2}B^{n_3}q_{prior}(m)\int x^{n_2+2n_3}\frac{1}{\sqrt{t}}\exp\left(-\frac{x^2}{2t}\right)dx\\
&\quad+\sum_{n_1+n_2+n_3=2q-1}\binom{2q-1}{n_1n_2n_3}(tR)^{n_1}A^{n_2}B^{n_3}\frac{\mathrm{d}}{\mathrm{d}m}q_{prior}(m)\int x^{n_2+2n_3+1}\frac{1}{\sqrt{t}}\exp\left(-\frac{x^2}{2t}\right)dx
\end{aligned}$$

where we have applied the Binomial theorem. Next, we use the Gaussian integral

$$\int\frac{1}{\sqrt{2\pi t}}x^k\exp\left(-\frac{x^2}{2t}\right)dx=t^{\frac{k}{2}}(k-1)!!\,\mathrm{Even}(k) \tag{66}$$

where $\mathrm{Even}(x)$ is 1 if $x$ is even and 0 else; similarly for $\mathrm{Odd}(x)$, to get

$$\begin{aligned}
0=&\sum_{n_1+n_2+n_3=2q-1}\binom{2q-1}{n_1n_2n_3}(tR)^{n_1}A^{n_2}B^{n_3}q_{prior}(m)t^{(n_2+2n_3)/2}\,\mathrm{Even}(n_2)(n_2+2n_3-1)!!\\
&+\sum_{n_1+n_2+n_3=2q-1}\binom{2q-1}{n_1n_2n_3}(tR)^{n_1}A^{n_2}B^{n_3}\frac{\mathrm{d}}{\mathrm{d}m}q_{prior}(m)t^{(n_2+2n_3+1)/2}\,\mathrm{Odd}(n_2)(n_2+2n_3)!!
\end{aligned}$$

The lowest order (in $t$) terms must be those where $n_1+n_3\leq 1$.[19] Computing all combinations and keeping only those where the parity matches leads to

$$\begin{aligned}
0=&A^{2q-1}\left(\frac{\mathrm{d}}{\mathrm{d}m}q_{prior}(m)\right)t^q(2q-2)!! && n_1=0,n_3=0,n_2=2q-1 && O(t^q)\\
&+(2q-1)A^{2q-2}Bq_{prior}(m)t^q(2q)!! && n_1=0,n_3=1,n_2=2q-2 && O(t^q)\\
&+(2q-1)RA^{2q-2}q_{prior}(m)t^q(2q-2)!! && n_1=1,n_3=0,n_2=2q-2 && O(t^q)\\
&+O(t^{q+1})
\end{aligned}$$

Now we divide[20] by $(2q-1)A^{2q-2}q_{prior}(m)t^q(2q-2)!!$ to get

$$0=A\cdot\left(\frac{\mathrm{d}}{\mathrm{d}m}\log q_{prior}(m)\right)\frac{1}{2q-1}+2qB+R+O(t^{q+1}) \tag{67}$$

[19]It can be verified that the higher-order terms suppressed above do not contribute to terms of that order.

[20]This is nonzero unless the prior or the Fisher information vanish at $m$, both of which we exclude.

Plugging in the definitions of $A$, $B$, and $q_{prior}$, and Eq. 60, leads to

$$\frac{1}{\mathcal{S}(\theta)}\frac{\mathrm{d}}{\mathrm{d}\theta}\log\frac{p_{prior}(\theta)}{\sqrt{\mathcal{S}(\theta)}}+(2q-1)\frac{1}{4}\frac{\mathrm{d}}{\mathrm{d}\theta}\frac{1}{\mathcal{S}(\theta)}+O(t)=R$$

Recalling $r=Rt+O(t^2)$ and substituting $\sigma^2=t$, we obtain

$$r=\sigma^2\frac{1}{\mathcal{S}}\frac{\mathrm{d}}{\mathrm{d}\theta}\log\frac{p_{prior}}{\sqrt{\mathcal{S}(\theta)}}+(2q-1)\sigma^2\frac{1}{4}\frac{\mathrm{d}}{\mathrm{d}\theta}\frac{1}{\mathcal{S}}+O(\sigma^4) \tag{68}$$

Note

$$\sigma^2\frac{1}{\mathcal{S}}\frac{\mathrm{d}}{\mathrm{d}\theta}\log\frac{1}{\sqrt{\mathcal{S}(\theta)}}=\sigma^2\frac{1}{\mathcal{S}(\theta)^2}\frac{-1}{2}\frac{\mathrm{d}}{\mathrm{d}\theta}\mathcal{S}(\theta)=\frac{\sigma^2}{2}\frac{\mathrm{d}}{\mathrm{d}\theta}\frac{1}{\mathcal{S}(\theta)} \tag{69}$$

and hence

$$r=\sigma^2\frac{1}{\mathcal{S}}\frac{\mathrm{d}}{\mathrm{d}\theta}\log p_{prior}(\theta)+\frac{2q+1}{4}\sigma^2\frac{\mathrm{d}}{\mathrm{d}\theta}\frac{1}{\mathcal{S}}+O(\sigma^4) \tag{70}$$

where all functions are evaluated at $\theta=F^{-1}(m)$. Recall $r=\widehat{\theta}-F^{-1}(m)$; so far, we have evaluated $\widehat{\theta}$ given a fixed encoding $m$.

In order to evaluate the decoding bias, we need to evaluate the expectation of this quantity over encodings $m$ for the stimulus $\theta$. In fact, we can show that Eq. (70) remains valid in expectation when the right-hand side of Eq. (70) is evaluated at $\theta$ instead of $F^{-1}(m)$; intuitively, this is because $|\theta-F^{-1}(m)|$ is $O(t)$ with high probability, so that the right-hand side does not differ much when evaluated at the two points for likely encodings $m$. More formally, if $\psi(\theta)$ is the function on the right-hand side of the equation evaluated at any $\theta$, then the decoding bias equals (using the Lagrange form of the remainder in Taylor's theorem):

$$\mathbb{E}\left[\psi(F^{-1}(m))|\theta\right]=\psi(\theta)+\underbrace{\mathbb{E}\left[(F^{-1}(m)-\theta)|\theta\right]}_{O(t)}\underbrace{\frac{\mathrm{d}}{\mathrm{d}\theta}\psi(\theta)}_{O(t)}+\frac{1}{2}\underbrace{\mathbb{E}\left[(F^{-1}(m)-\theta)^2\frac{\mathrm{d}^2}{\mathrm{d}\theta^2}\psi(\theta_m)|\theta\right]}_{O(t^2)} \tag{71}$$

where the last term is $O(t^2)$ because $|\frac{\mathrm{d}^2}{\mathrm{d}\theta^2}\psi(\theta_m)|=O(t)$ across all $m$, and $\mathbb{E}\left[(F^{-1}(m)-\theta)^2\right]=O(t)$. Hence, recalling that the exponent is $p=2q$ and substituting $\mathcal{J}=\frac{\mathcal{S}}{\sigma^2}$, we get to the following equation for the decoding bias:

$$\boxed{\mathbb{E}\left[\widehat{\theta}-F^{-1}(m)|\theta\right]=\frac{1}{\mathcal{J}}\frac{\mathrm{d}}{\mathrm{d}\theta}\left[\log p_{prior}(\theta)\right]+\frac{p+1}{4}\frac{\mathrm{d}}{\mathrm{d}\theta}\left[\frac{1}{\mathcal{J}}\right]+O(\sigma^4)}$$

where all functions are evaluated at the stimulus $\theta$. The argument in Eq. (71) is highly general and will be used in all derivations of decoding biases in this paper without repeating the details.

**S3.1.4.2 Odd exponent $>1$: $L^{2q+1}$ loss** We write the exponent as $p=2q+1$, where $q\geq 1$ is an integer. We first note that

$$\frac{d}{dx}x^{2q+1}=(2q+1)\cdot\mathrm{sign}(x)\cdot x^{2q} \tag{72}$$

Hence, analogously to Eq. (64), we aim to solve the following equation for $r$:

$$0=\int sign(r+Ax+Bx^2)(r+Ax+Bx^2)^{2q}\frac{1}{\sqrt{t}}\left(q_{prior}(m)+x\frac{\mathrm{d}}{\mathrm{d}m}q_{prior}(m)\right)\exp\left(-\frac{x^2}{2t}\right)dx \tag{73}$$

This argument is an extension of that given for even exponents, requiring additional care to account for the sign operator. The *sign* operator flips at (here, without loss of generality, we assume that $A,B>0$ – otherwise, the argument proceeds analogously):

$$\frac{-A\pm\sqrt{A^2-4Br}}{2B}=\frac{-A}{2B}\pm\left(\frac{A}{2B}-\frac{r}{A}\right)+O(r^2) \tag{74}$$

leading to the bounds

$$-\frac{A}{B}+\frac{r}{A}+O(r^2) \tag{75}$$

$$-\frac{r}{A}+O(r^2) \tag{76}$$

such that the sign is negative between these bounds, and positive elsewhere (because we assumed $B>0$ – else, the signs would flip). The first section, ending at $\approx -A/B$ carries exponentially little posterior mass if $t$ is small, and can be neglected (as $t\to 0$, the contribution of this interval is $\sim \exp\left(-\frac{A}{tB}\right)=o(t^k)$ for any $k>0$). Hence, up to higher-order terms, the integral can be written as

$$\begin{aligned}0=&\int_{-\infty}^{-r/A}(-1)(r+Ax+Bx^2)^{2q}\frac{1}{\sqrt{t}}\left(q_{prior}(m)+x\frac{\mathrm{d}}{\mathrm{d}m}q_{prior}(m)\right)\exp\left(-\frac{x^2}{2t}\right)dx\\&+\int_{-r/A}^{\infty}(r+Ax+Bx^2)^{2q}\frac{1}{\sqrt{t}}\left(q_{prior}(m)+x\frac{\mathrm{d}}{\mathrm{d}m}q_{prior}(m)\right)\exp\left(-\frac{x^2}{2t}\right)dx\end{aligned}$$

If – again without loss of generality – $r>0$, we can write this as

$$\begin{aligned}0=&\int_{-\infty}^{0}(-1)(r+Ax+Bx^2)^{2q}\frac{1}{\sqrt{t}}\left(q_{prior}(m)+x\frac{\mathrm{d}}{\mathrm{d}m}q_{prior}(m)\right)\exp\left(-\frac{x^2}{2t}\right)dx\\&+\int_{0}^{\infty}(r+Ax+Bx^2)^{2q}\frac{1}{\sqrt{t}}\left(q_{prior}(m)+x\frac{\mathrm{d}}{\mathrm{d}m}q_{prior}(m)\right)\exp\left(-\frac{x^2}{2t}\right)dx\\&+2\int_{-r/A}^{0}(r+Ax+Bx^2)^{2q}\frac{1}{\sqrt{t}}\left(q_{prior}(m)+x\frac{\mathrm{d}}{\mathrm{d}m}q_{prior}(m)\right)\exp\left(-\frac{x^2}{2t}\right)dx\end{aligned}$$

Because we already know that $r=O(t)$, the last term is bounded as $O(t^{2q})$, and is not of lowest order, so we can neglect it. Then we can write

$$\begin{aligned}0=&\int_{-\infty}^{\infty}\mathrm{sign}(x)(r+Ax+Bx^2)^{2q}\frac{1}{\sqrt{t}}q_{prior}(m)\exp\left(-\frac{x^2}{2t}\right)dx\\&+\int_{-\infty}^{\infty}\mathrm{sign}(x)(r+Ax+Bx^2)^{2q}\frac{1}{\sqrt{t}}x\frac{\mathrm{d}}{\mathrm{d}m}q_{prior}(m)\exp\left(-\frac{x^2}{2t}\right)dx\end{aligned}$$

Now expanding the powers using the Binomial theorem leads to

$$\begin{aligned}0=&q_{prior}(m)\sum_{n_1+n_2+n_3=2q}\binom{2q}{n_1n_2n_3}r^{n_1}A^{n_2}B^{n_3}\int_{-\infty}^{\infty}\mathrm{sign}(x)x^{n_2+2n_3}\frac{1}{\sqrt{t}}\exp\left(-\frac{x^2}{2t}\right)dx\\&+\frac{\mathrm{d}}{\mathrm{d}m}q_{prior}(m)\sum_{n_1+n_2+n_3=2q}\binom{2q}{n_1n_2n_3}r^{n_1}A^{n_2}B^{n_3}\int_{-\infty}^{\infty}\mathrm{sign}(x)x^{n_2+2n_3+1}\frac{1}{\sqrt{t}}\exp\left(-\frac{x^2}{2t}\right)dx\end{aligned}$$

The lowest-order terms are those where $n_1, n_3 = 0, 1$; only those integrals with an odd power of $x$ play a role:

$$\begin{aligned}
0 = &\left(\frac{\mathrm{d}}{\mathrm{d}m} q_{prior}(m)\right) \sum_{n_2=2q} A^{n_2} \int_{-\infty}^{\infty} \mathrm{sign}(x) x^{2q+1} \frac{1}{\sqrt{t}} \exp\left(-\frac{x^2}{2t}\right) dx && O(t^{q+0.5}) \\
&+ q_{prior}(m) \sum_{0+n_2+1=2q} 2qA^{n_2} B \int_{-\infty}^{\infty} \mathrm{sign}(x) x^{2q+1} \frac{1}{\sqrt{t}} \exp\left(-\frac{x^2}{2t}\right) dx && O(t^{q+0.5}) \\
&+ q_{prior}(m) \sum_{n_2=2q-1} 2qrA^{n_2} \int_{-\infty}^{\infty} \mathrm{sign}(x) x^{2q-1} \frac{1}{\sqrt{t}} \exp\left(-\frac{x^2}{2t}\right) dx && O(t^{q+0.5}) \\
&+ 2\left(\frac{\mathrm{d}}{\mathrm{d}m} q_{prior}(m)\right) \sum_{n_2=2q-2} 2q(2q-1)rA^{n_2} B \int_{-\infty}^{\infty} \mathrm{sign}(x) x^{2q+1} \frac{1}{\sqrt{t}} \exp\left(-\frac{x^2}{2t}\right) dx && O(t^{q+1.5}) \\
&+ h.o.t.
\end{aligned}$$

Only considering the first three terms, dividing by $q_{prior}(m)$ and by $A^{2q-1}$ and rearranging, we get

$$\begin{aligned}
0 = &A\left(\frac{\mathrm{d}}{\mathrm{d}m} \log q_{prior}(m)\right) \int_{-\infty}^{\infty} \mathrm{sign}(x) x^{2q+1} \frac{1}{\sqrt{t}} \exp\left(-\frac{x^2}{2t}\right) dx && O(t^{q+0.5}) \\
&+ 2qB \int_{-\infty}^{\infty} \mathrm{sign}(x) x^{2q+1} \frac{1}{\sqrt{t}} \exp\left(-\frac{x^2}{2t}\right) dx && O(t^{q+0.5}) \\
&+ 2qr \int_{-\infty}^{\infty} \mathrm{sign}(x) x^{2q-1} \frac{1}{\sqrt{t}} \exp\left(-\frac{x^2}{2t}\right) dx && O(t^{q+0.5}) \\
&+ O(t^{q+1.5})
\end{aligned}$$

Evaluating the three integrals using the fact

$$\int_{-\infty}^{\infty} \mathrm{sign}(x) x^{2k+1} e^{-ax^2} dx = 2\int_{0}^{\infty} x^{2k+1} e^{-ax^2} dx = \frac{k!}{a^{k+1}} \tag{77}$$

and simplifying leads to

$$0 = A\left(\frac{\mathrm{d}}{\mathrm{d}m} \log p_{prior}(m)\right) \frac{1}{\sqrt{t}} \frac{q!}{\frac{1}{(2t)^{q+1}}} + 2qB \frac{1}{\sqrt{t}} \frac{q!}{\frac{1}{(2t)^{q+1}}} + 2qr \frac{1}{\sqrt{t}} \frac{(q-1)!}{\frac{1}{(2t)^{q}}} + O(t^{q+1.5})$$

Rearranging and further simplifying leads to

$$r = -tA \frac{\mathrm{d}}{\mathrm{d}m} \log q_{prior}(m) - 2tqB + O(t^2)$$

Substituting the definitions of $A$ and $B$, and steps analogous to Eq. (69), lead to ($p := 2q+1$):

$$\boxed{r = \sigma^2 \frac{1}{\mathcal{S}} \frac{\mathrm{d}}{\mathrm{d}\theta} \log p_{prior} + \frac{p+1}{4} \sigma^2 \frac{\mathrm{d}}{\mathrm{d}\theta} \frac{1}{\mathcal{S}} + O(\sigma^4)}$$

evaluated at $F^{-1}(m)$. The argument in Eq. (71) then leads to the formula for the decoding bias, such that all quantities are instead evaluated at $\theta$.

**S3.1.4.3 Posterior Median: $L^1$ loss** Noting

$$\frac{\mathrm{d}}{\mathrm{d}x}|x| = \mathrm{sign}(x) \tag{78}$$

we want to solve

$$0 = \int sign(r + Ax + Bx^2)\frac{1}{\sqrt{t}}\left(q_{prior}(m) + x\frac{\mathrm{d}}{\mathrm{d}m}q_{prior}(m)\right)\exp\left(-\frac{x^2}{2t}\right)dx$$

As in Section S3.1.4.2, the sign changes at:

$$\frac{-A \pm \sqrt{A^2 - 4Br}}{2B} = \frac{-A}{2B} \pm \left(\frac{A}{2B} - \frac{r}{A}\right) + O(r^2) \tag{79}$$

leading to the bounds

$$-\frac{A}{B} + \frac{r}{A} + O(r^2) \tag{80}$$
$$-\frac{r}{A} + O(r^2) \tag{81}$$

and the sign is negative between these bounds, and positive elsewhere (assuming $B > 0$ without loss of generality – else, the signs would reverse). For small $t$, the integral can be written approximately (the first section, ending at $\approx -A/B$ carries exponentially little posterior mass if $t$ is small, and can be neglected) as

$$\begin{aligned} 0 = &\int_{-\infty}^{-r/A}(-1)\frac{1}{\sqrt{t}}\left(q_{prior}(m) + x\frac{\mathrm{d}}{\mathrm{d}m}q_{prior}(m)\right)\exp\left(-\frac{x^2}{2t}\right)dx \\ &+ \int_{-r/A}^{\infty}\frac{1}{\sqrt{t}}\left(q_{prior}(m) + x\frac{\mathrm{d}}{\mathrm{d}m}q_{prior}(m)\right)\exp\left(-\frac{x^2}{2t}\right)dx \end{aligned}$$

If (again WLOG) $r > 0$, we can write this as

$$\begin{aligned} 0 = &\int_{-\infty}^{\infty}\mathrm{sign}(x)\frac{1}{\sqrt{t}}\left(q_{prior}(m) + x\frac{\mathrm{d}}{\mathrm{d}m}q_{prior}(m)\right)\exp\left(-\frac{x^2}{2t}\right)dx \\ &+ 2\int_{-r/A}^{0}\frac{1}{\sqrt{t}}\left(q_{prior}(m) + x\frac{\mathrm{d}}{\mathrm{d}m}q_{prior}(m)\right)\exp\left(-\frac{x^2}{2t}\right)dx \end{aligned}$$

which we can expand as

$$\begin{aligned} 0 = &\int_{-\infty}^{\infty}\mathrm{sign}(x)\frac{1}{\sqrt{t}}q_{prior}(m)\exp\left(-\frac{x^2}{2t}\right)dx \\ &+ 2\int_{-r/A}^{0}\frac{1}{\sqrt{t}}q_{prior}(m)\exp\left(-\frac{x^2}{2t}\right)dx \\ &+ \int_{-\infty}^{\infty}\mathrm{sign}(x)\frac{1}{\sqrt{t}}x\left(\frac{\mathrm{d}}{\mathrm{d}x}q_{prior}(m)\right)\exp\left(-\frac{x^2}{2t}\right)dx \\ &+ 2\int_{-r/A}^{0}\frac{1}{\sqrt{t}}x\left(\frac{\mathrm{d}}{\mathrm{d}x}q_{prior}(m)\right)\exp\left(-\frac{x^2}{2t}\right)dx \end{aligned}$$

The first term evaluates to zero. Using the fact

$$\int_0^{\infty} xe^{-ax^2} = \frac{1}{2a} \tag{82}$$

we obtain

$$
\begin{aligned}
0 =& 2q_{prior}(m)\int_{-r/A}^{0}\frac{1}{\sqrt{t}}\exp\left(-\frac{x^2}{2t}\right)dx \\
&+2\frac{1}{\sqrt{t}}\left(\frac{\mathrm{d}}{\mathrm{d}m}q_{prior}(m)\right)t \\
&+2\frac{\mathrm{d}}{\mathrm{d}m}q_{prior}(m)\int_{-r/A}^{0}\frac{1}{\sqrt{t}}x\exp\left(-\frac{x^2}{2t}\right)dx
\end{aligned}
$$

As we want to solve for $r$, we need to understand the two integrals. We rescale the integrands by a factor of $\sqrt{t}$:

$$
\begin{aligned}
0 =& 2q_{prior}(m)\int_{0}^{\frac{r}{A\sqrt{t}}}\exp\left(-\frac{x^2}{2}\right)dx \\
&+2\frac{1}{\sqrt{t}}\left(\frac{\mathrm{d}}{\mathrm{d}m}q_{prior}(m)\right)t \\
&-2\frac{\mathrm{d}}{\mathrm{d}m}q_{prior}(m)\sqrt{t}\int_{0}^{\frac{r}{A\sqrt{t}}}x\exp\left(-\frac{x^2}{2}\right)dx
\end{aligned}
$$

and expand the exponentials under the integrals into power series that converge uniformly on the integration interval:

$$
\begin{aligned}
0 =& 2q_{prior}(m)\int_{0}^{\frac{r}{A\sqrt{t}}}\left(1-\frac{x^2}{4}+\frac{x^4}{16\cdot 2!}-\frac{x^6}{64\cdot 3!}+\ldots\right)dx \\
&+2\frac{1}{\sqrt{t}}\left(\frac{\mathrm{d}}{\mathrm{d}m}q_{prior}(m)\right)t \\
&-2\frac{\mathrm{d}}{\mathrm{d}m}q_{prior}(m)\sqrt{t}\int_{0}^{\frac{r}{A\sqrt{t}}}\left(x-\frac{x^3}{4}+\frac{x^5}{16\cdot 2!}-\frac{x^7}{64\cdot 3!}+\ldots\right)dx
\end{aligned}
$$

and obtain

$$
\begin{aligned}
0 =& 2q_{prior}(m)\left(\left(\frac{r}{A\sqrt{t}}\right)-\left(\frac{r}{A\sqrt{t}}\right)^3\cdot\frac{1}{12}+\left(\frac{r}{A\sqrt{t}}\right)^5\cdot\frac{1}{5\cdot 16\cdot 2!}+\ldots\right) \\
&+2\frac{\mathrm{d}}{\mathrm{d}m}q_{prior}(m)\sqrt{t} \\
&-2\frac{\mathrm{d}}{\mathrm{d}m}q_{prior}(m)\sqrt{t}\left(((\frac{r}{A\sqrt{t}})^2/2-(\frac{r}{A\sqrt{t}})^4/16+(\frac{r}{A\sqrt{t}})^6/(6\cdot 16\cdot 2!)+\ldots\right)
\end{aligned}
$$

Now first suppose that $r=\omega(t)$, then the lowest-order term is $2q_{prior}(m)\left(\frac{r}{A\sqrt{t}}\right)=\omega(\sqrt{t})$, but this is not zero – contradiction. So $r=O(t)$. After multiplying with $\sqrt{t}$, we obtain

$$
0=\frac{2\cdot r\cdot q_{prior}(m)}{A}+2\frac{\mathrm{d}}{\mathrm{d}m}q_{prior}(m)+O(t^2)
$$

which can be rewritten as

$$
0=-2\frac{q_{prior}}{\sqrt{S}}(r\sqrt{S})+2t\frac{q_{prior}}{\sqrt{S}}\frac{\mathrm{d}}{\mathrm{d}m}\log\frac{q_{prior}}{\sqrt{S(m)}}+O(t^2)
$$

Now recalling the definition of $q_{prior}$ leads to

$$0 = -2p_{prior}r + 2t\frac{p_{prior}}{\sqrt{\mathcal{S}}}\frac{\mathrm{d}}{\mathrm{d}m}\log\frac{p_{prior}}{\sqrt{\mathcal{S}(m)}} + O(t^2)$$

and finally

$$r = t\frac{1}{\sqrt{\mathcal{S}}}\frac{\mathrm{d}}{\mathrm{d}m}\log\frac{p_{prior}}{\sqrt{\mathcal{S}(m)}} + O(t^2)$$

Overall, we arrive at the result:

$$r = t\frac{1}{\mathcal{S}}\frac{\mathrm{d}}{\mathrm{d}\theta}\log\frac{p_{prior}}{\sqrt{\mathcal{S}(m)}} + O(t^2) \tag{83}$$

Rewriting using Eq. (69) leads to ($q := 1$):

$$\boxed{r = \sigma^2\frac{1}{\mathcal{S}}\frac{\mathrm{d}}{\mathrm{d}\theta}\log p_{prior} + \frac{q+1}{4}\sigma^2\frac{\mathrm{d}}{\mathrm{d}\theta}\frac{1}{\mathcal{S}} + O(\sigma^4)}$$

The argument in Eq. (71) then leads to the formula for the decoding bias.

**S3.1.4.4 MAP Estimator** The bias of the MAP estimator was already found for a model with Gaussian encoding by Stocker and Simoncelli [28]; we recapitulate an analogous argument for our observer model and with careful tracking of the order of the remainder. The posterior has the density given in Eq. 43. As before, we may assume without loss of generality $F(0) = 0 = m$. At the posterior mode, $\frac{\mathrm{d}}{\mathrm{d}\theta}\log p(\widehat{\theta}|m) = 0$ and thus:

$$\begin{aligned}
0 =& \sigma^2\frac{\mathrm{d}}{\mathrm{d}\theta}\log p_{Posterior}(\widehat{\theta}|m)\\
=& \sigma^2\frac{\mathrm{d}}{\mathrm{d}\theta}\log p_{lik}(m|\widehat{\theta}) + \sigma^2\frac{\mathrm{d}}{\mathrm{d}\theta}\log p_{prior}(\widehat{\theta})\\
=& -\frac{1}{2}\frac{\mathrm{d}}{\mathrm{d}\theta}|F(\widehat{\theta}) - m|^2 + \sigma^2\frac{\mathrm{d}}{\mathrm{d}\theta}\log p_{prior}(\widehat{\theta})\\
=& -\left(\frac{\mathrm{d}}{\mathrm{d}\theta}F(\widehat{\theta})\right)\cdot(F(\widehat{\theta}) - m) + \sigma^2\frac{\mathrm{d}}{\mathrm{d}\theta}\log p_{prior}(\widehat{\theta})\\
=& -\mathcal{S}\cdot\widehat{\theta} + \sigma^2\frac{\mathrm{d}}{\mathrm{d}\theta}\log p_{prior}(\widehat{\theta}) + O((\widehat{\theta} - m)^2)\\
=& -\mathcal{S}\cdot\widehat{\theta} + \sigma^2\frac{\mathrm{d}}{\mathrm{d}\theta}\log p_{prior}(0) + \sigma^2\widehat{\theta}\frac{\mathrm{d}^2}{\mathrm{d}\theta^2}\log p_{prior}(0) + O(\widehat{\theta}^2)
\end{aligned}$$

The bias $(\widehat{\theta} - F^{-1}(m)) = \widehat{\theta} - 0 = \widehat{\theta}$ must be of order $O(\sigma^2)$ because, otherwise, the lowest-order term would be $-\mathcal{S}\cdot\widehat{\theta} = \omega(\sigma^2)$, which is not zero, and the overall expression could not be zero. Hence

$$\sigma^2\theta\frac{\mathrm{d}^2}{\mathrm{d}\theta^2}\log p_{prior}(0) = O(\theta^2) = O(\sigma^4) \tag{84}$$

Then, with $F^{-1}(m) = 0 = m$, rearranging leads to

$$\widehat{\theta} - F^{-1}(m) = \sigma^2\frac{1}{\mathcal{S}}\frac{\mathrm{d}}{\mathrm{d}\theta}\log p_{prior}(F^{-1}(m)) + O(\sigma^4) \tag{85}$$

By the same argument as in Eq. (71), the expectation of Eq. (85) is:

$$\boxed{\mathbb{E}\left[\widehat{\theta} - F^{-1}(m)\right] = \sigma^2\frac{1}{\mathcal{S}}\frac{\mathrm{d}}{\mathrm{d}\theta}\log p_{prior}(\theta) + O(\sigma^4)}$$

###### S3.1.4.5 More General Loss Functions

We now consider a general symmetric loss function that is smooth at $x = 0$; such a function must have an even Taylor expansion:

$$\ell(x) = ax^p + bx^{p+2} + O(x^{p+4}) \tag{86}$$

where $p \in \{2, 4, \dots\}$. In this case, we want to solve

$$0 = \int \ell'(r + Ax + Bx^2) \frac{1}{\sqrt{t}} (p_{prior}(m) + x \frac{\mathrm{d}}{\mathrm{d}m} p_{prior}(m)) \exp\left(-\frac{x^2}{2t}\right) dx$$

Writing $p = 2q$, we get

$$\begin{aligned} 0 =& 2aq \int (r + Ax + Bx^2)^{2q-1} \frac{1}{\sqrt{t}} (p_{prior}(m) + x \frac{\mathrm{d}}{\mathrm{d}m} p_{prior}(m)) \exp\left(-\frac{x^2}{2t}\right) dx \\ &+ \int O(|r + Ax + Bx^2|^{2q+1}) \frac{1}{\sqrt{t}} (p_{prior}(m) + x \frac{\mathrm{d}}{\mathrm{d}m} p_{prior}(m)) \exp\left(-\frac{x^2}{2t}\right) dx \end{aligned}$$

Plugging in the expressions evaluating the integrals found individually for each power, we find that only the first term contributes terms of the lowest order. Hence, up to error $O(t^2) = O(\sigma^4)$, the bias is determined by the lowest-order term of $\ell$. Our results in Theorem 1 regarding $L^p$ losses in the small-noise regime thus extend to all symmetric loss functions that are smooth at 0.

#### S3.2 Proof of Theorem 2: Interaction of External and Internal Noise

As before, we use $t$ for the sensory noise variance $\sigma^2$, and additionally use $s$ for the stimulus noise variance $\tau^2$. The encoding is given as

$$m = F(\theta+\varepsilon)+\delta \tag{87}$$

where $\varepsilon, \delta$ are isotropic, $\varepsilon \sim N(0,s)$, $\delta \sim N(0,t)$. Here, we prove the following theorem:

**Theorem S2.** *Let $p > 0$ be even. Assume $\theta$ is in the interior of $\mathcal{X}$. The bias is*

$$\left(\frac{\sigma^2}{\mathcal{S}}+\tau^2\right)(\log p_{prior}(\theta))' + \left[1+\frac{p-2}{4}\frac{1}{1+\tau^2\mathcal{J}}\right]\left(\frac{\sigma^2}{\mathcal{S}}\right)' \tag{88}$$

*up to a remainder of order $O\left((\sigma^4+\tau^4+\sigma^2\tau^2)\cdot C_{F,p_{prior},p,\theta}\right)$, for $\sigma,\tau > 0$ sufficiently small.*

This result is illustrated by simulaions in Figure S5: First, stimulus noise increases prior attraction; Eq. (88) shows that prior attraction effects responding to sensory noise and to stimulus noise combine additively. Second, when $p=2$, likelihood repulsion is not influenced by stimulus noise. However, when $p > 2$, increasing stimulus noise decreases likelihood repulsion. This equally applies to even or odd exponents in simulation, showing that the conclusions of the theorem generalize to odd exponents.

We prove the theorem in two subsections below; as before, we separately treat the encoding and decoding components of the bias.

##### S3.2.1 Encoding Bias

We can write the encoding bias depending on sensory variance $t=\sigma^2$ and stimulus variance $s=\tau^2$ as follows:

$$\frac{\sigma^2}{4}\frac{\mathrm{d}}{\mathrm{d}\theta}\frac{1}{\mathcal{S}(\theta)} + O(\sigma^2\tau^2)+O(\sigma^4)+O(\tau^4) \tag{89}$$

where we've suppressed constants depending on $F, p_{prior}, \theta$ as elsewhere.

*Proof.* Taking into account both types of noise, the likelihood has the density

$$p(m|\theta) = \int_{-\infty}^{\infty} \exp\left(-\frac{(m-F(\theta+\varepsilon))^2}{2t}-\frac{\varepsilon^2}{2s}\right)\frac{1}{2\pi\sqrt{ts}}d\varepsilon \tag{90}$$

Using Eq (90):

$$\begin{aligned}
&\int_{-\infty}^{\infty} F^{-1}(m)\int_{-\infty}^{\infty}\exp\left(-\frac{(m-F(\theta+\varepsilon))^2}{2t}-\frac{\varepsilon^2}{2s}\right)\frac{1}{2\pi\sqrt{ts}}d\varepsilon dm\\
=&\int_{-\infty}^{\infty}\frac{1}{\sqrt{2\pi s}}\exp\left(-\frac{\varepsilon^2}{2s}\right)\int_{-\infty}^{\infty}F^{-1}(m)\exp\left(-\frac{(m-F(\theta+\varepsilon))^2}{2t}\right)\frac{1}{\sqrt{2\pi t}}dmd\varepsilon\\
=&\int_{-\infty}^{\infty}\frac{1}{\sqrt{2\pi s}}\exp\left(-\frac{\varepsilon^2}{2s}\right)\left[t\frac{1}{4}\frac{\mathrm{d}}{\mathrm{d}\theta}\frac{1}{\mathcal{S}(\theta+\varepsilon)}+O(t^2)\right]d\varepsilon\\
=&\int_{-\infty}^{\infty}\frac{1}{\sqrt{2\pi s}}\exp\left(-\frac{\varepsilon^2}{2s}\right)\left[t\frac{1}{4}\frac{\mathrm{d}}{\mathrm{d}\theta}\frac{1}{\mathcal{S}(\theta)}+t\frac{\varepsilon^2}{2}\frac{1}{4}\frac{\mathrm{d}^3}{\mathrm{d}\theta^3}\frac{1}{\mathcal{S}(\theta)}+O(t\varepsilon^4)\right]d\varepsilon+O(t^2)\\
=&t\frac{1}{4}\frac{\mathrm{d}}{\mathrm{d}\theta}\frac{1}{\mathcal{S}(\theta)}+\frac{ts}{2}\frac{1}{4}\frac{\mathrm{d}^3}{\mathrm{d}\theta^3}\frac{1}{\mathcal{S}(\theta)}+O(ts^2)+O(t^2)\\
=&t\frac{1}{4}\frac{\mathrm{d}}{\mathrm{d}\theta}\frac{1}{\mathcal{S}(\theta)}+O(ts)+O(t^2)
\end{aligned}$$

where we dropped odd orders of $\varepsilon$ in the Taylor approximation, as they vanish under the Gaussian integral. □

###### S3.2.2 Decoding Bias

**Theorem S3.** *Let $p > 0$ be even. The decoding bias is*

$$\left(\frac{1}{\mathcal{J}}+\tau^2\right)(\log p_{prior}(\theta))' + \left[\frac{9}{4}+\frac{p-1}{4}\frac{1}{1+\tau^2\mathcal{J}}\right]\left(\frac{1}{\mathcal{J}}\right)' \tag{91}$$

*up to a remainder of order $O\left((\sigma^4+\tau^4+\sigma^2\tau^2)\cdot C_{F,p_{prior},p,\theta}\right)$, for $\sigma,\tau > 0$ sufficiently small.*

We will write

$$\alpha = \frac{s}{t} = \frac{\tau^2}{\sigma^2}$$

We may assume without loss of generality that the observed encoding is $m = 0$, and $F(0) = 0$. We will write

$$q := p-1 \tag{92}$$

where $q$ is odd.

Then, as in Section S3.1.4.1, but now carrying out all integrals in stimulus space, we want to solve the following equation for $r$:

$$\begin{aligned}
0 =& \frac{\mathrm{d}}{\mathrm{d}r}\int_{-\infty}^{\infty}(r-\theta)^{q+1}p_{prior}(\theta)p(\theta|m)d\theta \\
\propto& \int_{-\infty}^{\infty}(r-\theta)^{q}p_{prior}(\theta)p(\theta|m)d\theta \\
\propto& \int_{-\infty}^{\infty}(r-\theta)^{q}p_{prior}(\theta)p(m|\theta)d\theta \\
=& \int_{-\infty}^{\infty}(r-\theta)^{q}p_{prior}(\theta)\int_{-\infty}^{\infty}p(0|\theta+\varepsilon)p(\varepsilon)d\varepsilon d\theta \\
=& \int_{-\infty}^{\infty}(r-\theta)^{q}p_{prior}(\theta)\int_{-\infty}^{\infty}\exp\left(-\frac{(F(\theta+\varepsilon))^2}{2t}\right)\exp\left(-\frac{\varepsilon^2}{2s}\right)\frac{1}{2\pi\sqrt{ts}}d\varepsilon d\theta \\
=& \int_{-\infty}^{\infty}\int_{-\infty}^{\infty}(r-\theta)^{q}p_{prior}(\theta)\exp\left(-\frac{(F(\theta+\varepsilon))^2}{2t}\right)\exp\left(-\frac{\varepsilon^2}{2s}\right)\frac{1}{2\pi\sqrt{ts}}d\varepsilon d\theta
\end{aligned}$$

where we have used Eq. (90). Rather than calculating this directly, we first compute this integral when the integrand is a monomial $\theta^k$, and then apply Taylor's theorem. Write

$$\begin{aligned}
\mathcal{A} :=& \alpha + \left[\frac{\mathrm{d}}{\mathrm{d}m}F^{-1}(0)\right]^2 \\
\mathcal{B} :=& \frac{\mathrm{d}}{\mathrm{d}m}F^{-1}(0) \\
\mathcal{C} :=& \frac{\mathrm{d}^2}{\mathrm{d}m^2}F^{-1}(0)
\end{aligned}$$

The key technical tool will be the following result:

**Proposition S4.** *Let $k > 0$ be an integer. The integral*

$$\mathcal{K}(k) := \int\int_{-\infty}^{\infty}\theta^k\exp\left(-\frac{(F(\theta+\varepsilon))^2}{2t}\right)\frac{1}{\sqrt{2\pi t}}d\theta\frac{1}{\sqrt{2\pi s}}\exp\left(-\varepsilon^2/(2s)\right)d\varepsilon \tag{93}$$

*equals, if $k$ is even:*

$$\mathcal{K}(k) = t^{\frac{k}{2}}(k-1)!!\mathcal{A}^{k/2}\mathcal{B} + O\left(t^{\frac{k+2}{2}}\right) \tag{94}$$

*and if k is odd:*

$$\mathcal{K}(k) = t^{\frac{k+1}{2}}\left[\frac{3}{2}k!!\mathcal{A}^{\frac{k-1}{2}}\mathcal{B}\mathcal{C} + \frac{1}{2}k!!(k-1)\mathcal{A}^{\frac{k-3}{2}}\mathcal{B}^3\mathcal{C}\right] + O\left(t^{\frac{k+3}{2}}\right) \tag{95}$$

As elsewhere, we suppress the dependency of the remainder on objects that do not depend on the noise parameters $t$ or $s = \alpha t$ (i.e., its dependency on $F$, $p_{prior}$, $p$, $\theta$). This proposition is shown in Section S3.2.2.1. We first show how the theorem follows:

*Proof of the Theorem.* As explained above, we want to solve the following equation for $r$:

$$\begin{aligned} 0 &= \int_{-\infty}^{\infty} (r-\theta)^q p_{prior}(\theta) p(\theta|m) d\theta \\ &\propto \int_{-\infty}^{\infty} (r-\theta)^q p_{prior}(\theta) p(m|\theta) d\theta \end{aligned}$$

Applying Taylor's theorem to the prior, and suppressing the remainder $O(\theta^3)$ for readability, we get

$$\begin{aligned} &= \sum_{i=0}^{q} \binom{q}{i} \int_{-\infty}^{\infty} r^i (-1)^{q-i} \theta^{q-i} (p_{prior}(\theta) + \theta \frac{\mathrm{d}}{\mathrm{d}\theta} p_{prior}(\theta) + \frac{\theta^2}{2} \frac{\mathrm{d}^2}{\mathrm{d}\theta^2} p_{prior}(\theta)) p(m|\theta) d\theta \\ &= \sum_{i=0}^{q} \binom{q}{i} r^i (-1)^{q-i} \int_{-\infty}^{\infty} \theta^{q-i} \cdot \left( p_{prior}(\theta) + \theta \frac{\mathrm{d}}{\mathrm{d}\theta} p_{prior}(\theta) + \frac{\theta^2}{2} \frac{\mathrm{d}^2}{\mathrm{d}\theta^2} p_{prior}(\theta) \right) p(m|\theta) d\theta \end{aligned}$$

Expanding, substituting the definition of $\mathcal{K}$, and indicating the order of each term (where $r = O(t)$, by reasoning as in Section S3.1.4.1):

$$\begin{aligned} 0 = &\sum_{i=0}^{q} \binom{q}{i} r^i (-1)^{q-i} p_{prior}(\theta) \mathcal{K}(q-i) && O(t^{i+\frac{(q-i)}{2}}) \\ &+ \sum_{i=0}^{q} \binom{q}{i} r^i (-1)^{q-i} \frac{\mathrm{d}}{\mathrm{d}\theta} p_{prior}(\theta) \mathcal{K}(q-i+1) && O(t^{i+\frac{(q-i+1)}{2}}) \\ &+ \frac{1}{2} \sum_{i=0}^{q} \binom{q}{i} r^i (-1)^{q-i} \frac{\mathrm{d}^2}{\mathrm{d}\theta^2} p_{prior}(\theta) \mathcal{K}(q-i+2) && O(t^{i+\frac{(q-i+2)}{2}}) \end{aligned}$$

We get the lowest orders at $i = 0, 1$. Considering that $q$ is odd, we obtain the following equation:

$$\begin{aligned} &p_{prior}(\theta) \mathcal{K}(q) && O(t^{\frac{(q+1)}{2}}) \\ &+ \frac{\mathrm{d}}{\mathrm{d}\theta} p_{prior}(\theta) \mathcal{K}(q+1) && O(t^{\frac{(q+1)}{2}}) \\ &+ 1/2 \frac{\mathrm{d}^2}{\mathrm{d}\theta^2} p_{prior}(\theta) \mathcal{K}(q+2) && O(t^{\frac{(q+3)}{2}}) \\ = &qr p_{prior}(\theta) \mathcal{K}(q-1) && O(t^{\frac{(q+1)}{2}}) \\ &+ qr \frac{\mathrm{d}}{\mathrm{d}\theta} p_{prior}(\theta) \mathcal{K}(q) && O(t^{\frac{(q+3)}{2}}) \\ &+ 1/2 qr \frac{\mathrm{d}^2}{\mathrm{d}\theta^2} p_{prior}(\theta) \mathcal{K}(q+1) && O(t^{\frac{(q+3)}{2}}) \\ &+ O(t^{\frac{q+3}{2}}) \end{aligned}$$

Hence, we need to solve

$$\mathcal{K}(q)+\frac{\mathrm{d}}{\mathrm{d}\theta}\log p_{prior}(\theta)\mathcal{K}(q+1)=qr\mathcal{K}(q-1)+O(t^{\frac{q+3}{2}}) \tag{96}$$

Inserting the expressions for $\mathcal{K}(\cdot)$ obtained in the proposition, considering that $q$ is odd:

$$t^{\frac{q+1}{2}}\left[\frac{3}{2}q!!\mathcal{A}^{\frac{q-1}{2}}\mathcal{B}\mathcal{C}+\frac{1}{2}q!!(q-1)\mathcal{A}^{\frac{q-3}{2}}\mathcal{B}^3\mathcal{C}\right]+\frac{\mathrm{d}}{\mathrm{d}\theta}\log p_{prior}(\theta)t^{\frac{q+1}{2}}(q+1-1)!!\mathcal{A}^{\frac{q+1}{2}}\mathcal{B}=qrt^{\frac{q-1}{2}}(q-1-1)!!\mathcal{A}^{\frac{q-1}{2}}\mathcal{B}+O(t^{\frac{q+3}{2}}) \tag{97}$$

and dividing both sides by $q!!t^{\frac{q-1}{2}}\mathcal{A}^{\frac{q-1}{2}}\mathcal{B}$, we get:

$$r=t\left[\frac{3}{2}\mathcal{C}+\frac{1}{2}(q-1)\mathcal{A}^{-1}\mathcal{B}^2\mathcal{C}\right]+t\mathcal{A}\frac{\mathrm{d}}{\mathrm{d}\theta}\log p_{prior}(\theta)+O(t^2) \tag{98}$$

and slight rewriting leads to the result:

$$r=t\mathcal{C}\left[\frac{3}{2}+\frac{q-1}{2}\frac{\mathcal{B}^2}{\alpha+\mathcal{B}^2}\right]+t(\alpha+\mathcal{B}^2)\frac{\mathrm{d}}{\mathrm{d}\theta}\log p_{prior}(\theta)+O(t^2)$$

Recall

$$\mathcal{A}:=\alpha+\left[\frac{\mathrm{d}}{\mathrm{d}m}F^{-1}(m)\right]^2$$
$$\mathcal{B}:=\frac{\mathrm{d}}{\mathrm{d}m}F^{-1}(0)$$
$$\mathcal{C}:=\frac{\mathrm{d}^2}{\mathrm{d}m^2}F^{-1}(0)$$

and thus

$$r=\sigma^2\left(\frac{\mathrm{d}^2}{\mathrm{d}m^2}F^{-1}(0)\right)\left[\frac{3}{2}+\frac{q-1}{2}\frac{(\frac{\mathrm{d}}{\mathrm{d}m}F^{-1}(0))^2}{\frac{\tau^2}{\sigma^2}+(\frac{\mathrm{d}}{\mathrm{d}m}F^{-1}(0))^2}\right]+\sigma^2\left(\frac{\tau^2}{\sigma^2}+\left(\frac{\mathrm{d}}{\mathrm{d}m}F^{-1}(0)\right)^2\right)\frac{\mathrm{d}}{\mathrm{d}\theta}\log p_{prior}(\theta)+O(\sigma^4)$$

Now

$$\frac{\mathrm{d}}{\mathrm{d}m}F^{-1}(0)=\frac{1}{\frac{\mathrm{d}}{\mathrm{d}\theta}F(0)}=\frac{1}{\sqrt{\mathcal{S}}}$$

and as in Eq. (54)

$$\frac{\mathrm{d}^2}{\mathrm{d}m^2}F^{-1}(0)=\frac{1}{2}\frac{\mathrm{d}}{\mathrm{d}\theta}\frac{1}{\mathcal{S}}$$

leading to the result:

$$r=\sigma^2\left[\frac{3}{4}+\frac{p-2}{4}\frac{1}{\frac{\mathcal{S}\tau^2}{\sigma^2}+1}\right]\left(\frac{\mathrm{d}}{\mathrm{d}\theta}\frac{1}{\mathcal{S}}\right)+\left(\tau^2+\frac{\sigma^2}{\mathcal{S}}\right)\frac{\mathrm{d}}{\mathrm{d}\theta}\log p_{prior}(\theta)+O(\sigma^4)+O(\tau^4)+O(\sigma^2\tau^2)$$

Combining this with the encoding bias (Eq. 89) yields the result. □

**S3.2.2.1 Proving Proposition S4** We aim to compute the lowest-order terms (in $t$) of:

$$\int\int_{-\infty}^{\infty}\theta^p\exp\left(-\frac{(F(\theta+\varepsilon))^2}{2t}\right)\frac{1}{\sqrt{2\pi t}}d\theta\frac{1}{\sqrt{2\pi s}}\exp\left(-\varepsilon^2/(2s)\right)d\varepsilon$$

First, we reparameterize $\theta \mapsto \theta - \varepsilon$:

$$= \int \int_{-\infty}^{\infty} (\theta - \varepsilon)^p \exp\left(-\frac{(F(\theta))^2}{2t}\right) \frac{1}{\sqrt{2\pi t}} d\theta \frac{1}{\sqrt{2\pi s}} \exp\left(-\varepsilon^2/(2s)\right) d\varepsilon$$

In order to separately treat $\theta$ and $\varepsilon$, we use the Binomial Theorem:

$$= \sum_{i=0}^{p} \binom{p}{i} \int (-\varepsilon)^{p-i} \frac{1}{\sqrt{2\pi s}} \exp\left(-\varepsilon^2/(2s)\right) d\varepsilon \int_{-\infty}^{\infty} \theta^i \exp\left(-\frac{(F(\theta))^2}{2t}\right) \frac{1}{\sqrt{2\pi t}} d\theta$$
$$= \sum_{i=0}^{p} \binom{p}{i} (-1)^{p-i}(p-i-1)!!s^{\frac{p-i}{2}} \int_{-\infty}^{\infty} \theta^i \exp\left(-\frac{(F(\theta))^2}{2t}\right) \frac{1}{\sqrt{2\pi t}} d\theta$$

Here, we have integrated out $\varepsilon$ using

$$\int_{-\infty}^{\infty} \frac{x^k}{2\pi t} \exp\left(-\frac{x^2}{2t}\right) dx = \mathrm{Even}(k)(k-1)!!t^{\frac{k}{2}} \tag{99}$$

where $x!! = x \cdot (x-2) \cdot (x-4) \cdot ... 3 \cdot 1$ (if $x$ is odd, ending in $4 \cdot 2$ if $x$ is even) is the double factorial. In order to integrate over $\theta$, we transform into the sensory space:

$$= \sum_{i=0}^{p} \binom{p}{i} (-1)^{p-i}(p-i-1)!!s^{\frac{p-i}{2}} \int_{-\infty}^{\infty} \left(F^{-1}(m)\right)^i \left(\frac{\mathrm{d}}{\mathrm{d}m} F^{-1}(m)\right) \exp\left(-\frac{m^2}{2t}\right) \frac{1}{\sqrt{2\pi t}} dm$$

Applying Taylor's theorem to both $F^{-1}$ and $\frac{\mathrm{d}}{\mathrm{d}m}F^{-1}$, and suppressing the remainders of order $O(m^3)$ for readability:

$$= \sum_{i=0}^{p} \binom{p}{i} (-1)^{p-i}(p-i-1)!!s^{\frac{p-i}{2}}$$
$$\int_{-\infty}^{\infty} \left(m \frac{\mathrm{d}}{\mathrm{d}m} F^{-1}(0) + \frac{m^2}{2} \frac{\mathrm{d}^2}{\mathrm{d}m^2} F^{-1}(0)\right)^i \left(\frac{\mathrm{d}}{\mathrm{d}m} F^{-1}(0) + m \frac{\mathrm{d}^2}{\mathrm{d}m^2} F^{-1}(0) + \frac{m^2}{2} \frac{\mathrm{d}}{\mathrm{d}m}^3 F^{-1}(0)\right) \exp\left(-\frac{m^2}{2t}\right) \frac{1}{\sqrt{2\pi t}} dm$$

Once again applying the Binomial theorem:

$$= \sum_{i=0}^{p} \binom{p}{i} (-1)^{p-i}(p-i-1)!!s^{\frac{p-i}{2}}$$
$$\int_{-\infty}^{\infty} \sum_{w=0}^{i} \binom{i}{w} m^w \left(\frac{\mathrm{d}}{\mathrm{d}m} F^{-1}(0)\right)^w \left(\frac{m^2}{2}\right)^{i-w} \left(\frac{\mathrm{d}^2}{\mathrm{d}m^2} F^{-1}(0)\right)^{i-w} \left(\frac{\mathrm{d}}{\mathrm{d}m} F^{-1}(0) + m \frac{\mathrm{d}^2}{\mathrm{d}m^2} F^{-1}(0) + \frac{m^2}{2} \frac{\mathrm{d}}{\mathrm{d}m}^3 F^{-1}(0)\right)$$
$$\exp\left(-\frac{m^2}{2t}\right) \frac{1}{\sqrt{2\pi t}} dm$$
$$= \sum_{i=0}^{p} \sum_{w=0}^{i} \binom{i}{w} \binom{p}{i} \frac{1}{2^{i-w}} (-1)^{p-i}(p-i-1)!!s^{\frac{p-i}{2}} \left(\frac{\mathrm{d}}{\mathrm{d}m} F^{-1}(0)\right)^w \left(\frac{\mathrm{d}^2}{\mathrm{d}m^2} F^{-1}(0)\right)^{i-w}$$
$$\int_{-\infty}^{\infty} m^{w+2\cdot(i-w)} \left(\frac{\mathrm{d}}{\mathrm{d}m} F^{-1}(0) + m \frac{\mathrm{d}^2}{\mathrm{d}m^2} F^{-1}(0) + \frac{m^2}{2} \frac{\mathrm{d}^3}{\mathrm{d}m^3} F^{-1}(0)\right) \exp\left(-\frac{m^2}{2t}\right) \frac{1}{\sqrt{2\pi t}} dm$$

Separating out the terms:

$$=\sum_{i=0}^{p}\sum_{w=0}^{i}\binom{i}{w}\binom{p}{i}(1/2)^{i-w}(-1)^{p-i}(p-i-1)!!s^{\frac{p-i}{2}}\left(\frac{\mathrm{d}}{\mathrm{d}m}F^{-1}(0)\right)^{w}\left(\frac{\mathrm{d}^2}{\mathrm{d}m^2}F^{-1}(0)\right)^{i-w}\left(\frac{\mathrm{d}}{\mathrm{d}m}F^{-1}(0)\right)$$
$$\int_{-\infty}^{\infty}m^{w+2\cdot(i-w)}\exp\left(-\frac{m^2}{2t}\right)\frac{1}{\sqrt{2\pi t}}dm$$
$$+\sum_{i=0}^{p}\sum_{w=0}^{i}\binom{i}{w}\binom{p}{i}\frac{1}{2^{i-w}}(-1)^{p-i}(p-i-1)!!s^{\frac{p-i}{2}}\left(\frac{\mathrm{d}}{\mathrm{d}m}F^{-1}(0)\right)^{w}\left(\frac{\mathrm{d}^2}{\mathrm{d}m^2}F^{-1}(0)\right)^{i-w}\left(\frac{\mathrm{d}^2}{\mathrm{d}m^2}F^{-1}(0)\right)$$
$$\int_{-\infty}^{\infty}m^{w+2\cdot(i-w)+1}\exp\left(-\frac{m^2}{2t}\right)\frac{1}{\sqrt{2\pi t}}dm$$
$$+\sum_{i=0}^{p}\sum_{w=0}^{i}\binom{i}{w}\binom{p}{i}\frac{1}{2^{i-w}}(-1)^{p-i}(p-i-1)!!s^{\frac{p-i}{2}}\left(\frac{\mathrm{d}}{\mathrm{d}m}F^{-1}(0)\right)^{w}\left(\frac{\mathrm{d}^2}{\mathrm{d}m^2}F^{-1}(0)\right)^{i-w}\left(\frac{\mathrm{d}^3}{\mathrm{d}m^3}F^{-1}(0)\right)$$
$$\int_{-\infty}^{\infty}m^{w+2\cdot(i-w)+2}\frac{1}{2}\exp\left(-\frac{m^2}{2t}\right)\frac{1}{\sqrt{2\pi t}}dm$$

and carrying out the Gaussian integrals, and using the definitions of $\mathcal{B},\mathcal{C}$, we arrive at:

$$=\sum_{i=0}^{p}\sum_{w=0}^{i}t^{\frac{p+i-w}{2}}\binom{i}{w}\binom{p}{i}\frac{(-1)^{p-i}}{2)^{i-w}}(p-i-1)!!\alpha^{\frac{p-i}{2}}\mathcal{B}^{w+1}\mathcal{C}^{i-w}(2i-w-1)!!\,\mathrm{Even}(2i-w)\,\mathrm{Even}(p-i)$$
$$+\sum_{i=0}^{p}\sum_{w=0}^{i}t^{\frac{p+i-w+1}{2}}\binom{i}{w}\binom{p}{i}\frac{(-1)^{p-i}}{2^{i-w}}(p-i-1)!!\alpha^{\frac{p-i}{2}}\mathcal{B}^{w}\mathcal{C}^{i-w+1}(2i-w)!!\,\mathrm{Even}(2i-w+1)\,\mathrm{Even}(p-i)$$
$$+\sum_{i=0}^{p}\sum_{w=0}^{i}t^{\frac{p+i-w+2}{2}}\binom{i}{w}\binom{p}{i}\frac{(-1)^{p-i}}{2^{i-w+1}}(p-i-1)!!\alpha^{\frac{p-i}{2}}\mathcal{B}^{w}\mathcal{C}^{i-w}\left(\frac{\mathrm{d}^3}{\mathrm{d}m^3}F^{-1}(0)\right)(2i-w+1)!!\,\mathrm{Even}(2i-w)\,\mathrm{Even}(p-i)$$

We only require the lowest order terms in $t$; these are attained when $w=i,i-1$. We will separately consider these cases, and for even and odd $p$. In all four cases, the reasoning will be analogous: We will obtain partial cancellation between double factorials and binomial coefficients, leaving each summand with a binomial coefficient as the only factor depending on $i$, and apply the binomial theorem to obtain a power of $\mathcal{A}$. Specifically, we will use the following fact to convert double factorials into standard factorials: For any positive integer $m$,

$$(2m)!!=2^m m! \tag{100}$$

In particular,

$$\binom{n}{k}=\frac{n!}{k!(n-k)!}=\frac{(2n)!!}{(2k)!!(2(n-k))!!}=:\left(\binom{2n}{2k}\right) \tag{101}$$

**Part A: $w=i$, and $p$ is even** Inserting $w=i$, we obtain

$$=t^{\frac{p}{2}}\sum_{i=0}^{p}\binom{p}{i}(-1)^{p-i}(p-i-1)!!\alpha^{\frac{p-i}{2}}\left(\frac{\mathrm{d}}{\mathrm{d}m}F^{-1}(m)\right)^{i+1}(i-1)!!$$
$$+t^{\frac{p+1}{2}}\sum_{i=0}^{p}\binom{p}{i}(-1)^{p-i}(p-i-1)!!\alpha^{\frac{p-i}{2}}\left(\frac{\mathrm{d}}{\mathrm{d}m}F^{-1}(m)\right)^{i}\left(\frac{\mathrm{d}^2}{\mathrm{d}m^2}F^{-1}(m)\right)(i)!!$$
$$+\frac{1}{2}t^{\frac{p+2}{2}}\sum_{i=0}^{p}\binom{p}{i}(-1)^{p-i}(p-i-1)!!\alpha^{\frac{p-i}{2}}\left(\frac{\mathrm{d}}{\mathrm{d}}F^{-1}(m)\right)^{i}\frac{\mathrm{d}^3}{\mathrm{d}m^3}F^{-1}(m)(i+1)!!$$

The third term has order $O(t^{\frac{p+2}{2}})$, and we focus on the first two terms. If $p$ is even, the second term is zero, and we have:

$$=t^{\frac{p}{2}} \sum_{i=0,even}^{p} \binom{p}{i} (p-i-1)!!\alpha^{\frac{p-i}{2}} \mathcal{B}^{i+1}(i-1)!!$$

where we have written $\sum_{i=0,even}^{p}$ for $\sum_{i=0,2,4,\ldots,p}$; we will continue using analogous notation below. Spelling out the binomial coefficients:

$$=t^{\frac{p}{2}} \sum_{i=0,even}^{p} \frac{p!(i-1)!!(p-i-1)!!}{i!(p-i)!} \alpha^{\frac{p-i}{2}} \mathcal{B}^{i+1}$$

Applying the identity $k! = k!!(k-1)!!$, we rewrite as:

$$=t^{\frac{p}{2}} \sum_{i=0,even}^{p} \frac{p!!(p-1)!!(i-1)!!(p-i-1)!!}{i!!(i-1)!!(p-i)!!(p-i-1)!!} \alpha^{\frac{p-i}{2}} \mathcal{B}^{i+1}$$

Now terms cancel, and we get

$$=t^{\frac{p}{2}} \sum_{i=0,even}^{p} (p-1)!! \frac{p!!}{i!!(p-i)!!} \alpha^{\frac{p-i}{2}} \mathcal{B}^{i+1}$$
$$=t^{\frac{p}{2}} (p-1)!! \sum_{i=0,even}^{p} \left(\binom{p}{i}\right) \alpha^{\frac{p-i}{2}} \mathcal{B}^{i+1}$$

Reparameterizing $j := i/2$ and applying Eq. (101):

$$=t^{\frac{p}{2}} (p-1)!! \sum_{j=0}^{p/2} \binom{p/2}{j} \alpha^{\frac{p}{2}-j} \mathcal{B}^{2j+1}$$

Once again applying the Binomial theorem, but in the other direction, we obtain the result:

$$\boxed{t^{\frac{p}{2}} (p-1)!! [\alpha + \mathcal{B}^2]^{\frac{p}{2}} \mathcal{B}}$$

**Part B: $w = i$, and $p$ is odd** The reasoning is analogous to the A part. If $p$ is odd, only the second term remains:

$$\left(\frac{\mathrm{d}^2}{\mathrm{d}^2} F^{-1}(m)\right) t^{\frac{p+1}{2}} \sum_{i=0,odd}^{p} \binom{p}{i} (p-i-1)!!\alpha^{\frac{p-i}{2}} \left(\frac{\mathrm{d}}{\mathrm{d}} F^{-1}(m)\right)^{i} i!!$$

We spell out the binomial coefficient:

$$=\left(\frac{\mathrm{d}^2}{\mathrm{d}^2} F^{-1}(m)\right) t^{\frac{p+1}{2}} \sum_{i=0,odd}^{p} \frac{p!(p-i-1)!!(i)!!}{i!(p-i)!} \alpha^{\frac{p-i}{2}} \left(\frac{\mathrm{d}}{\mathrm{d}} F^{-1}(m)\right)^{i}$$

and again use the identity $k! = k!!(k-1)!!$:

$$
\begin{aligned}
&= Ct^{\frac{p+1}{2}} \sum_{i=0,odd}^{p} \frac{p!!(p-1)!!(p-i-1)!!(i)!!}{i!!(i-1)!!(p-i)!!(p-i-1)!!} \alpha^{\frac{p-i}{2}} \mathcal{B}^{i} \\
&= Ct^{\frac{p+1}{2}} \sum_{i=0,odd}^{p} p!! \frac{(p-1)!!}{(i-1)!!((p-1)-(i-1))!!} \alpha^{\frac{p-i}{2}} \mathcal{B}^{i} \\
&= Ct^{\frac{p+1}{2}} \sum_{i=0,odd}^{p} p!! \left( \binom{p-1}{i-1} \right) \alpha^{\frac{p-i}{2}} \mathcal{B}^{i}
\end{aligned}
$$

Substituting $j := i-1$, so $i = j+1$, and then $k := j/2$:

$$
\begin{aligned}
&= Ct^{\frac{p+1}{2}} \sum_{j=0,even}^{p-1} p!! \left( \binom{p-1}{j} \right) \alpha^{\frac{p-j-1}{2}} \mathcal{B}^{j+1} \\
&= Ct^{\frac{p+1}{2}} \sum_{k=0}^{\frac{p-1}{2}} p!! \binom{\frac{p-1}{2}}{k} \alpha^{\frac{p-1}{2}-k} \mathcal{B}^{2k+1}
\end{aligned}
$$

With the Binomial theorem, we obtain the result:

$$
\boxed{Ct^{\frac{p+1}{2}} p!! \mathcal{B} \left[ \alpha + \mathcal{B}^2 \right]^{\frac{(p-1)}{2}}}
$$

**Part C:** $w = i-1$**, and** $p$ **is even** Substituting $w = i-1$:

$$
\begin{aligned}
&= (\frac{d^2}{d^2} F^{-1}(m)) \sum_{i=0,odd}^{p} i \binom{p}{i} (1/2)(-1)^{p-i}(p-i-1)!!\alpha^{\frac{p-i}{2}} t^{\frac{p+1}{2}} (\frac{d}{d} F^{-1}(m))^{i}(i)!! \\
&+ (\frac{d^2}{d^2} F^{-1}(m))^2 \sum_{i=0,even}^{p} i \binom{p}{i} (1/2)(-1)^{p-i}(p-i-1)!!\alpha^{\frac{p-i}{2}} t^{\frac{p+2}{2}} (\frac{d}{d} F^{-1}(m))^{i-1}(i+1)!! \\
&+ (\frac{d^2}{d^2} F^{-1}(m)) \frac{d^3}{dm^3} F^{-1}(m) \sum_{i=0,odd}^{p} i \binom{p}{i} (1/2)(-1)^{p-i}(p-i-1)!!\alpha^{\frac{p-i}{2}} t^{\frac{p+3}{2}} (\frac{d}{d} F^{-1}(m))^{i-1}(i+2)!!(1/2)
\end{aligned}
$$

The third term has order $t^{\frac{p+3}{2}}$ and need not concern us further. If $p$ is even, only the second term remains:

$$
= t^{\frac{p+2}{2}} \left( \frac{d^2}{dm^2} F^{-1}(m) \right)^2 \sum_{i=0,even}^{p} ip! \frac{(p-i-1)!!(i+1)!!}{i!(p-i)!} (1/2) \alpha^{\frac{p-i}{2}} \left( \frac{d}{d} F^{-1}(m) \right)^{i-1}
$$

As it has order $t^{\frac{p+2}{2}}$, it need not concern us further.

**Part D:** $w = i-1$**, and** $p$ **is odd** The first term remains:

$$
= \sum_{i=1,3,\dots,p} i \frac{1}{2} \frac{p!(p-i-1)!!(i)!!}{i!(p-i)!} \alpha^{\frac{p-i}{2}} t^{\frac{p+1}{2}} \left( \frac{d}{d} F^{-1}(m) \right)^{i} \left( \frac{d^2}{d^2} F^{-1}(m) \right)
$$

Rewriting factorials as double factorials, and canceling:

$$
\begin{aligned}
&= \sum_{i=1,3,\ldots,p} i\frac{1}{2}\frac{p!!(p-1)!!(p-i-1)!!(i)!!}{i!!(i-1)!!(p-i)!!(p-i-1)!!}\alpha^{\frac{p-i}{2}}t^{\frac{p+1}{2}}\left(\frac{\mathrm{d}}{\mathrm{d}}F^{-1}(m)\right)^{i}\left(\frac{\mathrm{d}^2}{\mathrm{d}^2}F^{-1}(m)\right)\\
&= \sum_{i=1,3,\ldots,p} i\frac{1}{2}\frac{p!!(p-1)!!}{(i-1)!!(p-i)!!}\alpha^{\frac{p-i}{2}}t^{\frac{p+1}{2}}\mathcal{B}^i\mathcal{C}\\
&= \sum_{i=1,3,\ldots,p} i\frac{1}{2}p!!\frac{(p-1)!!}{(i-1)!!((p-1)-(i-1))!!}\alpha^{\frac{p-i}{2}}t^{\frac{p+1}{2}}\mathcal{B}^i\mathcal{C}
\end{aligned}
$$

In order to absorb the factor $i$ in a binomial coefficient, we write it as $(i-1)+1$:

$$
=t^{\frac{p+1}{2}}\sum_{i=1,3,\ldots,p}\frac{1}{2}p!!\frac{(i-1+1)(p-1)!!}{(i-1)!!((p-1)-(i-1))!!}\alpha^{\frac{p-i}{2}}\mathcal{B}^i\mathcal{C}
$$

Separating terms:

$$
\begin{aligned}
=&t^{\frac{p+1}{2}}\sum_{i=1,3,\ldots,p}\frac{1}{2}p!!\frac{(i-1)(p-1)!!}{(i-1)!!((p-1)-(i-1))!!}\alpha^{\frac{p-i}{2}}\mathcal{B}^i\mathcal{C}\\
+&t^{\frac{p+1}{2}}\sum_{i=1,3,\ldots,p}\frac{1}{2}p!!\frac{(p-1)!!}{(i-1)!!((p-1)-(i-1))!!}\alpha^{\frac{p-i}{2}}\mathcal{B}^i\mathcal{C}
\end{aligned}
$$

The first term is zero when $i=1$; we can thus start summation at 3. Then, we can cancel $(i-1)$ in the first term, and both terms are ready for conversion into a binomial coefficient:

$$
\begin{aligned}
=&t^{\frac{p+1}{2}}\sum_{i=3,5,\ldots,p}\frac{1}{2}p!!(p-1)\frac{(p-3)!!}{(i-3)!!((p-3)-(i-3))!!}\alpha^{\frac{p-i}{2}}\mathcal{B}^i\mathcal{C}\\
+&t^{\frac{p+1}{2}}\sum_{i=1,3,\ldots,p}\frac{1}{2}p!!\frac{(p-1)!!}{(i-1)!!((p-1)-(i-1))!!}\alpha^{\frac{p-i}{2}}\mathcal{B}^i\mathcal{C}\\
=[A]&t^{\frac{p+1}{2}}\sum_{i=3,5,\ldots,p}\frac{1}{2}p!!(p-1)\left(\binom{p-3}{i-3}\right)\alpha^{\frac{p-i}{2}}\mathcal{B}^i\mathcal{C}\\
+[B]&t^{\frac{p+1}{2}}\sum_{i=1,3,\ldots,p}\frac{1}{2}p!!\left(\binom{p-1}{i-1}\right)\alpha^{\frac{p-i}{2}}\mathcal{B}^i\mathcal{C}
\end{aligned}
$$

where we have separated into terms [A], [B]. For the first term, we substitute $k := \frac{i-3}{2}$, so $i = 2k+3$:

$$
[A] = t^{\frac{p+1}{2}}\sum_{k=0}^{\frac{p-3}{2}}\frac{1}{2}p!!(p-1)\left(\binom{p-3}{2k}\right)\alpha^{\frac{p-3}{2}-k}\mathcal{B}^{2k+3}\mathcal{C}
$$

and obtain for [A]:

$$
\boxed{t^{\frac{p+1}{2}}\frac{1}{2}p!!(p-1)[\alpha-\mathcal{B}^2]^{\frac{p-3}{2}}\mathcal{B}^3\mathcal{C}}
$$

For the second term, we substitute $k := \frac{i-1}{2}$, so $i = 2k+1$:

$$[B] = t^{\frac{p+1}{2}} \sum_{k=0}^{\frac{p-1}{2}} \frac{1}{2} p!! \left( \binom{p-1}{2k} \right) \alpha^{\frac{p-1}{2}-k} \mathcal{B}^{2k+1} \mathcal{C}$$

and obtain for [B]:

$$\boxed{t^{\frac{p+1}{2}} \frac{1}{2} p!! (\alpha + \mathcal{B}^2)^{\frac{p-1}{2}} \mathcal{B}\mathcal{C}}$$

##### S3.3 Proof of Theorem 3: Boundary Effects

We assume that the support of the prior is $[-\infty, \theta_{Max}]$. Write

$$Q := F(\theta_{Max}) - F(\theta) \tag{102}$$

and

$$D := \frac{Q}{\sigma} \tag{103}$$

As in Section S3.2, we focus on proving the result for even positive exponents.

**Theorem S5** (Theorem 3 from main text)**.** *Assume $p_{prior}(x) \equiv 0$ when $x > \theta_{Max}$. Assume $\theta < \theta_{Max}$. For some quantity $C_{1,p,D,\theta,F,\sigma}$ and universal constants $C_{2,p,D}, C_{3,p,D}$, the bias (for even integers $p > 0$) is given as*

$$\underbrace{-\frac{C_{1,p,D,\theta,F,\sigma}}{\sqrt{\mathcal{J}}}}_{Regression} + C_{2,p,D} \underbrace{\frac{1}{\mathcal{J}} (\log p_{prior})'}_{Prior\ Attraction} + C_{3,p,D} \underbrace{\frac{p+2}{4} \left(\frac{1}{\mathcal{J}}\right)'}_{Likelihood\ Repulsion} \tag{104}$$

*up to approximation error of order $O(\sigma^3 \cdot C_{F,p_{prior},p,\theta,D})$ when $\sigma > 0$ is sufficiently small, where $C_{\dots} > 0$ and $C_{1,p,D,\theta,F,\sigma} = \Theta(1)$ as $\sigma \to 0$, $\lim_{D\to\infty} C_{1,p,D,\theta,F,\sigma} = 0$, $\lim_{D\to\infty} C_{2/3,p,D} = 1$.*

The encoding bias is unchanged, as the likelihood is unchanged.[21] However, the decoding bias now reflects the boundary effect; this is proven below, focusing on even exponents $p > 0$.

We note the following. For stimuli close to the boundary, such that $C_{1,p,D,\theta,F,\sigma}$ is on the order of 1, the regression effect scales with $\frac{1}{\sqrt{\mathcal{J}}} \propto \sigma$, whereas the other components scale with $\frac{1}{\mathcal{J}} \propto \sigma^2$. When noise is small, the first dominates the second ($\sigma^2 = o(\sigma)$). Hence, we expect that the boundary effect will overwhelm other biases in the vicinity of the boundary. However, when the effective distance $Q/\sigma$ increases, the boundary effect disappears because $C_{1,p,D,\theta,F,\sigma} \to 0$. We illustrate the interaction between the regression effect and the other components in Figure S6: Boundary effects indeed overwhelm the other components close to the boundary, but disappear in the interior. Second, we plot $C_{1,p,D,\theta,F,\sigma}$ for both even and odd exponents in Figure S7; the regression effect rapidly decays to zero with the effective distance, and is larger for higher exponents.

We now proceed to proving the theorem, calculating the decoding bias for positive even exponents. Throughout, we will write the exponent as $p = 2q$. As the prior is truncated at $Q$, the same will be true for the posterior. Hence, the Bayes estimator is defined as

$$\widehat{\theta} := \arg_{\widehat{\theta}} \min \int_{-\infty}^{Q} |\widehat{\theta} - \theta|^{2q} P(\theta|m) d\theta \tag{105}$$

Performing the same steps as in Section S3.1.4.1, assuming $m = F^{-1}(m) = 0$ for now and writing $Q = F(\theta_{Max}) - m$[22], we can write the problem – up to a higher-order remainder – as:

$$\begin{aligned}
0 &= \int_{-\infty}^{Q} (r + Ax + Bx^2)^{2q-1} \frac{1}{\sqrt{t}} (q_{prior}(m) + x \frac{\mathrm{d}}{\mathrm{d}x} q_{prior}(m)) \exp\left(-\frac{x^2}{2t}\right) dx \\
&= \int_{-\infty}^{Q} \left( \sum_{n_1+n_2+n_3=2q-1} \binom{2q-1}{n_1 n_2 n_3} r^{n_1} A^{n_2} x^{n_2+2n_3} B^{n_3} \right) \frac{1}{\sqrt{t}} (q_{prior}(m) + x \frac{\mathrm{d}}{\mathrm{d}m} q_{prior}(m)) \exp\left(-\frac{x^2}{2t}\right) dx \\
&= \sum_{n_1+n_2+n_3=2q-1} \binom{2q-1}{n_1 n_2 n_3} r^{n_1} A^{n_2} B^{n_3} q_{prior}(m) \int_{-\infty}^{Q} x^{n_2+2n_3} \frac{1}{\sqrt{t}} \exp\left(-\frac{x^2}{2t}\right) dx \\
&\quad + \sum_{n_1+n_2+n_3=2q-1} \binom{2q-1}{n_1 n_2 n_3} r^{n_1} A^{n_2} B^{n_3} \frac{\mathrm{d}}{\mathrm{d}m} q_{prior}(m) \int_{-\infty}^{Q} x^{n_2+2n_3+1} \frac{1}{\sqrt{t}} \exp\left(-\frac{x^2}{2t}\right) dx
\end{aligned}$$

[21] Under an alternative model where the sensory space itself has a boundary, so that the likelihood is also truncated, there is an additional regression effect from the encoding bias; the overall conclusions will be the same.

[22] We note that $Q$ can be negative for any individual $m$, even though the average, and thus the quantity referred to the in theorem, is always positive as $\theta < \theta_{Max}$.

where we have applied the Binomial theorem in the second step. We note that for any positive integer $k$, as long as $Q$ is finite,

$$\int_{-\infty}^{Q} x^k \frac{1}{\sqrt{t}} \exp\left(-\frac{x^2}{2t}\right) dx = t^{k/2} \int_{-\infty}^{Q/\sqrt{t}} x^k \exp\left(-\frac{x^2}{2}\right) dx = \Theta(t^{k/2}) \tag{106}$$

Hence, the lowest-order terms will be those where $n_3 = 0, 1$, so the following is true to lowest order:

$$\begin{aligned}
0 = &\sum_{n_1+n_2=2q-1} \binom{2q-1}{n_1 n_2} r^{n_1} A^{n_2} q_{prior}(m) \int_{-\infty}^{Q} x^{n_2} \frac{1}{\sqrt{t}} \exp\left(-\frac{x^2}{2t}\right) dx \\
&+ \sum_{n_1+n_2=2q-1} \binom{2q-1}{n_1 n_2} r^{n_1} A^{n_2} \frac{\mathrm{d}}{\mathrm{d}m} q_{prior}(m) \int_{-\infty}^{Q} x^{n_2+1} \frac{1}{\sqrt{t}} \exp\left(-\frac{x^2}{2t}\right) dx \\
&+ \sum_{n_1+n_2+1=2q-1} \binom{2q-1}{n_1 n_2 1} r^{n_1} A^{n_2} B q_{prior}(m) \int_{-\infty}^{Q} x^{n_2+2} \frac{1}{\sqrt{t}} \exp\left(-\frac{x^2}{2t}\right) dx \\
&+ \sum_{n_1+n_2+1=2q-1} \binom{2q-1}{n_1 n_2 1} r^{n_1} A^{n_2} B \frac{\mathrm{d}}{\mathrm{d}m} q_{prior}(m) \int_{-\infty}^{Q} x^{n_2+3} \frac{1}{\sqrt{t}} \exp\left(-\frac{x^2}{2t}\right) dx
\end{aligned}$$

We write

$$r = v\sqrt{t} + wt + O(t^2)$$

Multiplying out the powers, the lowest orders will be those where $wt$ receives power 0 or 1. First, consider those terms where $wt$ receives power 0:

$$\begin{aligned}
= &\sum_{n_1+n_2=2q-1} \binom{2q-1}{n_1 n_2} (v\sqrt{t})^{n_1} A^{n_2} q_{prior}(m) \int_{-\infty}^{Q} x^{n_2} \frac{1}{\sqrt{t}} \exp\left(-\frac{x^2}{2t}\right) dx \\
&+ \sum_{n_1+n_2=2q-1} \binom{2q-1}{n_1 n_2} (v\sqrt{t})^{n_1} A^{n_2} \frac{\mathrm{d}}{\mathrm{d}m} q_{prior}(m) \int_{-\infty}^{Q} x^{n_2+1} \frac{1}{\sqrt{t}} \exp\left(-\frac{x^2}{2t}\right) dx \\
&+ \sum_{n_1+n_2+1=2q-1} \binom{2q-1}{n_1 n_2 1} (v\sqrt{t})^{n_1} A^{n_2} B q_{prior}(m) \int_{-\infty}^{Q} x^{n_2+2} \frac{1}{\sqrt{t}} \exp\left(-\frac{x^2}{2t}\right) dx \\
&+ \sum_{n_1+n_2+1=2q-1} \binom{2q-1}{n_1 n_2 1} (v\sqrt{t})^{n_1} A^{n_2} B \frac{\mathrm{d}}{\mathrm{d}m} q_{prior}(m) \int_{-\infty}^{Q} x^{n_2+3} \frac{1}{\sqrt{t}} \exp\left(-\frac{x^2}{2t}\right) dx
\end{aligned}$$

Appealing to the Binomial theorem,

$$\begin{aligned}
= &q_{prior}(m) \int_{-\infty}^{Q} (v\sqrt{t} + Ax)^{2q-1} \frac{1}{\sqrt{t}} \exp\left(-\frac{x^2}{2t}\right) dx \\
&+ \frac{\mathrm{d}}{\mathrm{d}m} q_{prior}(m) \int_{-\infty}^{Q} (v\sqrt{t} + Ax)^{2q-1} x \frac{1}{\sqrt{t}} \exp\left(-\frac{x^2}{2t}\right) dx \\
&+ (2q-1) B q_{prior}(m) \int_{-\infty}^{Q} (v\sqrt{t} + A)^{2q-2} x^2 \frac{1}{\sqrt{t}} \exp\left(-\frac{x^2}{2t}\right) dx \\
&+ (2q-1) B \frac{\mathrm{d}}{\mathrm{d}m} q_{prior}(m) \int_{-\infty}^{Q} (v\sqrt{t} + A)^{2q-2} x^3 \frac{1}{\sqrt{t}} \exp\left(-\frac{x^2}{2t}\right) dx
\end{aligned}$$

Second, those terms where $wt$ receives power 1:

$$
\begin{aligned}
= &\sum_{n_1+n_2=2q-1} \binom{2q-1}{n_1 n_2} (v\sqrt{t})^{n_1-1}(wt)n_1 A^{n_2} q_{prior}(m) \int_{-\infty}^{Q} x^{n_2} \frac{1}{\sqrt{t}} \exp\left(-\frac{x^2}{2t}\right) dx \\
&+ \sum_{n_1+n_2=2q-1} \binom{2q-1}{n_1 n_2} (v\sqrt{t})^{n_1-1}(wt)n_1 A^{n_2} \frac{\mathrm{d}}{\mathrm{d}m} q_{prior}(m) \int_{-\infty}^{Q} x^{n_2+1} \frac{1}{\sqrt{t}} \exp\left(-\frac{x^2}{2t}\right) dx \\
&+ \sum_{n_1+n_2+1=2q-1} \binom{2q-1}{n_1 n_2 1} (v\sqrt{t})^{n_1-1}(wt)n_1 A^{n_2} B q_{prior}(m) \int_{-\infty}^{Q} x^{n_2+2} \frac{1}{\sqrt{t}} \exp\left(-\frac{x^2}{2t}\right) dx \\
&+ \sum_{n_1+n_2+1=2q-1} \binom{2q-1}{n_1 n_2 1} (v\sqrt{t})^{n_1-1}(wt)n_1 A^{n_2} B \frac{\mathrm{d}}{\mathrm{d}m} q_{prior}(m) \int_{-\infty}^{Q} x^{n_2+3} \frac{1}{\sqrt{t}} \exp\left(-\frac{x^2}{2t}\right) dx
\end{aligned}
$$

Rewriting the binomial coefficients, where the sums are restricted to $n_1 > 0$:

$$
\begin{aligned}
= &\sum_{n_1-1+n_2=2q-2} (2q-1) \frac{(2q-2)!}{(n_1-1)!((2q-2)-(n_1-1))!} (wt) q_{prior}(m) \int_{-\infty}^{Q} (v\sqrt{t})^{n_1-1} A^{n_2} x^{n_2} \frac{1}{\sqrt{t}} \exp\left(-\frac{x^2}{2t}\right) dx \\
&+ \sum_{n_1+n_2=2q-1} (2q-1) \frac{(2q-2)!}{(n_1-1)!((2q-2)-(n_1-1))!} (wt) \frac{\mathrm{d}}{\mathrm{d}m} q_{prior}(m) \int_{-\infty}^{Q} (v\sqrt{t})^{n_1-1} A^{n_2} x^{n_2+1} \frac{1}{\sqrt{t}} \exp\left(-\frac{x^2}{2t}\right) dx \\
&+ (2q-1)(2q-2) \sum_{(n_1-1)+n_2=2q-3} \frac{(2q-3)!}{(n_1-1)!((2q-3)-(n_1-1))!} (wt) B q_{prior}(m) \int_{-\infty}^{Q} (v\sqrt{t})^{n_1-1} A^{n_2} x^{n_2+2} \frac{1}{\sqrt{t}} \exp\left(-\frac{x^2}{2t}\right) dx \\
&+ (2q-1)(2q-2) \sum_{(n_1-1)+n_2=2q-3} \frac{(2q-3)!}{(n_1-1)!((2q-3)-(n_1-1))!} (wt) B \frac{\mathrm{d}}{\mathrm{d}m} q_{prior}(m) \int_{-\infty}^{Q} (v\sqrt{t})^{n_1-1} A^{n_2} x^{n_2+3} \frac{1}{\sqrt{t}} \exp\left(-\frac{x^2}{2t}\right) dx
\end{aligned}
$$

and applying the Binomial theorem,

$$
\begin{aligned}
= &(2q-1) wt q_{prior}(m) \int_{-\infty}^{Q} (v\sqrt{t}+Ax)^{2q-2} \frac{1}{\sqrt{t}} \exp\left(-\frac{x^2}{2t}\right) dx \\
&+ (2q-1) wt \frac{\mathrm{d}}{\mathrm{d}m} q_{prior}(m) \int_{-\infty}^{Q} (v\sqrt{t}+Ax)^{2q-2} x \frac{1}{\sqrt{t}} \exp\left(-\frac{x^2}{2t}\right) dx \\
&+ (2q-1)(2q-2) wt B q_{prior}(m) \int_{-\infty}^{Q} (v\sqrt{t}+Ax)^{2q-3} x^2 \frac{1}{\sqrt{t}} \exp\left(-\frac{x^2}{2t}\right) dx \\
&+ (2q-1)(2q-2) wt B \frac{\mathrm{d}}{\mathrm{d}m} q_{prior}(m) \int_{-\infty}^{Q} (v\sqrt{t}+Ax)^{2q-3} x^3 \frac{1}{\sqrt{t}} \exp\left(-\frac{x^2}{2t}\right) dx
\end{aligned}
$$

Putting both groups of terms together, and dividing by the prior, gives us:

$$
\begin{aligned}
0 = &\int_{-\infty}^{Q} (v\sqrt{t}+Ax)^{2q-1} \frac{1}{\sqrt{t}} \exp\left(-\frac{x^2}{2t}\right) dx \\
&+ \frac{\mathrm{d}}{\mathrm{d}m} \log q_{prior}(m) \int_{-\infty}^{Q} (v\sqrt{t}+Ax)^{2q-1} x \frac{1}{\sqrt{t}} \exp\left(-\frac{x^2}{2t}\right) dx \\
&+ (2q-1) B \int_{-\infty}^{Q} (v\sqrt{t}+A)^{2q-2} x^2 \frac{1}{\sqrt{t}} \exp\left(-\frac{x^2}{2t}\right) dx \\
&+ (2q-1) B \frac{\mathrm{d}}{\mathrm{d}m} \log q_{prior}(m) \int_{-\infty}^{Q} (v\sqrt{t}+A)^{2q-2} x^3 \frac{1}{\sqrt{t}} \exp\left(-\frac{x^2}{2t}\right) dx
\end{aligned}
$$

$$+(2q-1)wt\int_{-\infty}^{Q}(v\sqrt{t}+Ax)^{2q-2}\frac{1}{\sqrt{t}}\exp\left(-\frac{x^2}{2t}\right)dx$$
$$+(2q-1)wt\frac{\mathrm{d}}{\mathrm{d}m}\log q_{prior}(m)\int_{-\infty}^{Q}(v\sqrt{t}+Ax)^{2q-2}x\frac{1}{\sqrt{t}}\exp\left(-\frac{x^2}{2t}\right)dx$$
$$+(2q-1)(2q-2)wtB\int_{-\infty}^{Q}(v\sqrt{t}+Ax)^{2q-3}x^2\frac{1}{\sqrt{t}}\exp\left(-\frac{x^2}{2t}\right)dx$$
$$+(2q-1)(2q-2)wtB\frac{\mathrm{d}}{\mathrm{d}m}\log q_{prior}(m)\int_{-\infty}^{Q}(v\sqrt{t}+Ax)^{2q-3}x^3\frac{1}{\sqrt{t}}\exp\left(-\frac{x^2}{2t}\right)dx$$

Reparameterizing the integrals using $y=\frac{x}{\sqrt{t}}$:

$$\begin{aligned}
0 =&\sqrt{t}^{2q-1}\int_{-\infty}^{Q/\sqrt{t}}(v+Ay)^{2q-1}\exp\left(-\frac{y^2}{2}\right)dy\\
&+\sqrt{t}^{2q}\frac{\mathrm{d}}{\mathrm{d}m}\log q_{prior}(m)\int_{-\infty}^{Q/\sqrt{t}}(v+Ay)^{2q-1}y\exp\left(-\frac{y^2}{2}\right)dy\\
&+(2q-1)B\int_{-\infty}^{Q/\sqrt{t}}(v+Ay)^{2q-2}\sqrt{t}^{2q}(y)^2\exp\left(-\frac{y^2}{2}\right)dy\\
&+(2q-1)B\sqrt{t}^{2q+1}\frac{\mathrm{d}}{\mathrm{d}m}\log q_{prior}(m)\int_{-\infty}^{Q/\sqrt{t}}(v+Ay)^{2q-2}y\exp\left(-\frac{y^2}{2}\right)dy\\
&+(2q-1)wt\sqrt{t}^{2q-2}\int_{-\infty}^{Q/\sqrt{t}}(v+Ay)^{2q-2}\exp\left(-\frac{y^2}{2}\right)dy\\
&+(2q-1)wt\sqrt{t}^{2q-1}\frac{\mathrm{d}}{\mathrm{d}m}\log q_{prior}(m)\int_{-\infty}^{Q/\sqrt{t}}(v+Ay))^{2q-2}y\exp\left(-\frac{y^2}{2}\right)dy\\
&+(2q-1)(2q-2)wt\sqrt{t}^{2q-1}B\int_{-\infty}^{Q/\sqrt{t}}(v+Ay)^{2q-3}(y)^2\exp\left(-\frac{y^2}{2}\right)dy\\
&+(2q-1)(2q-2)wt\sqrt{t}^{2q}B\frac{\mathrm{d}}{\mathrm{d}m}\log q_{prior}(m)\int_{-\infty}^{Q/\sqrt{t}}(v+Ay)^{2q-3}(y)^3\exp\left(-\frac{y^2}{2}\right)dy
\end{aligned}$$

The first term has the lowest order, giving us the equation:

$$0=\int_{-\infty}^{Q/\sqrt{t}}(v_0+y)^{2q-1}\exp\left(-\frac{y^2}{2}\right)dy$$

implicitly defining $v_0=\frac{v}{A}$. While we are not aware of a closed-form expression for the solution $H_{1,p,Q/\sigma}$[23], it is monotonically increasing as a function of $Q/\sigma$, and $H_{1,p,Q/\sigma}\to 0$ as $Q/\sigma\to\infty$. Its numerical behavior can be read off Figure S46.

We now consider the remaining terms in order to obtain $w$; the following three terms have the second-lowest order $\sqrt{t}^{2q}$:

$$\begin{aligned}
0 =&\sqrt{t}^{2q}\frac{\mathrm{d}}{\mathrm{d}m}\log q_{prior}(m)\int_{-\infty}^{Q/\sqrt{t}}(v+Ay)^{2q-1}y\exp\left(-\frac{y^2}{2}\right)dy\\
&+(2q-1)B\sqrt{t}^{2q}\int_{-\infty}^{Q/\sqrt{t}}(v+Ay)^{2q-2}(y)^2\exp\left(-\frac{y^2}{2}\right)dy
\end{aligned}$$

[23] It can be written as the solution of an algebraic equation of degree $\leq 2q$ with coefficients involving $\mathrm{erf}(A)$, by applying integration by parts and known [13] recurrence equation for the iterated integral of the error function.

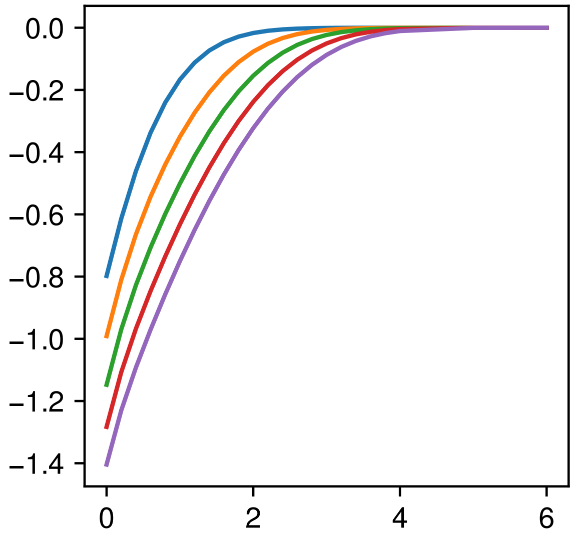

Figure S46: Behavior of $H_{1,p,D}$, the lowest-order magnitude of regression to the mean, as a function of $D := Q/\sigma$ (x-axis, positive values only) and the exponent $p$. To lowest order, the regression to the mean (simulated in Figure S7; that figure has the x-axis $\theta = \theta_{Max} - Q$) is given by $H_{1,p,D}/\sqrt{\mathcal{J}}$.

$$+(2q-1)w\sqrt{t}^{2q}\int_{-\infty}^{Q/\sqrt{t}}(v+Ay)^{2q-2}\exp\left(-\frac{y^2}{2}\right)dy$$

and thus

$$w = \frac{\mathrm{d}}{\mathrm{d}m}\log q_{prior}(m)\frac{\int_{-\infty}^{Q/\sqrt{t}}(v+Ay)^{2q-1}y\exp\left(-\frac{y^2}{2}\right)dy}{(2q-1)\int_{-\infty}^{Q/\sqrt{t}}(v+Ay)^{2q-2}\exp\left(-\frac{y^2}{2}\right)dy}+(2q-1)B\frac{\int_{-\infty}^{Q/\sqrt{t}}(v+Ay)^{2q-2}(y)^2\exp\left(-\frac{y^2}{2}\right)dy}{(2q-1)\int_{-\infty}^{Q/\sqrt{t}}(v+Ay)^{2q-2}\exp\left(-\frac{y^2}{2}\right)dy} \tag{107}$$

As before, it will be convenient to rewrite using $v_0 = H_{1,p}$ ($v = -Av_0$), and subsequently divide by $A^{2q-2}$. Then

$$w = A\left(\frac{\mathrm{d}}{\mathrm{d}m}\log q_{prior}(m)\right)\underbrace{\frac{\int_{-\infty}^{Q/\sigma}(x-v_0)^{2q-1}x\,dP(x)}{(2q-1)\int_{-\infty}^{Q/\sigma}(x-v_0)^{2q-2}dP(x)}}_{W_1}+(2q-1)B\underbrace{\frac{\int_{-\infty}^{Q/\sigma}(x-v_0)^{2q-2}x^2dP(x)}{(2q-1)\int_{-\infty}^{Q/\sigma}(x-v_0)^{2q-2}dP(x)}}_{W_2} \tag{108}$$

We now have a form analogous to that for the decoding bias in the absence of a boundary found in Eq. (67), but with the bias modified by factors $W_1$, $W_2$. These factors are determined by the effective distance $Q/\sigma$ to the boundary and the loss function exponent $q = 2p$. As $Q/\sigma \to \infty$, both factors converge to 1, recovering Eq. (67) (and thus Theorem 1) when the limit is far away. The numerical behavior for finite $Q/\sigma$ is shown in Figure S47: Both factors are in [0,1], and rapidly converge to 1 when the effective distance increases. Plugging in definitions of $A$, $B$, $q_{prior}$ as in Section S3.1.4.1, we have thus far obtained:

$$\widehat{\theta} - F^{-1}(m) = \underbrace{\frac{H_{1,p,D}}{\sqrt{\mathcal{J}}}}_{Regression} + C_{2,p,D}\underbrace{\frac{1}{\mathcal{J}}(\log p_{prior})'}_{Prior\ Attraction} + C_{3,p,D}\underbrace{\frac{q+2}{4}\left(\frac{1}{\mathcal{J}}\right)'}_{Likelihood\ Repulsion} + O(\sigma^3) \tag{109}$$

where $D$, $\mathcal{J}$, $p_{prior}$ are all evaluated at $F^{-1}(m)$. We now want to evaluate the decoding bias, i.e., the expectation over $m$. Rescaling as $D_0 = \frac{\theta}{\sigma}$, and using the Lagrange form of the remainder, we obtain for the lowest-order term in the decoding bias, the expectation of Eq. (109) (this is the quantity plotted in Figure S7) – here, $\tilde{D}$ is a function of $m$ and lies between $F^{-1}(m)/\sigma$ and $\theta/\sigma$, and we have $\mathbb{E}[\tilde{D} - D_0] = O(1)$ in $\sigma$:

$$\sigma\mathbb{E}\left[\frac{H_{1,p,D}}{\sqrt{\mathcal{S}}}\right] = \sigma\frac{H_{1,p,D_0}}{\sqrt{\mathcal{S}(D_0)}} + \sigma\mathbb{E}[\tilde{D} - D_0]\frac{\mathrm{d}}{\mathrm{d}D}\frac{H_{1,p,D_0}}{\sqrt{\mathcal{S}(D_0)}} \tag{110}$$

$$= \sigma\frac{H_{1,p,D_0}}{\sqrt{\mathcal{S}(D_0)}} + \mathbb{E}[\tilde{D} - D_0]\sigma\frac{\frac{\mathrm{d}}{\mathrm{d}D}H_{1,p,D_0}}{\sqrt{\mathcal{S}(D_0)}} + \mathbb{E}[\tilde{D} - D_0]H_{1,p,D_0}\sigma^2\frac{\mathrm{d}}{\mathrm{d}\theta}\frac{1}{\sqrt{\mathcal{S}(D_0)}} \tag{111}$$

$$= \sigma\frac{H_{1,p,D_0}}{\sqrt{\mathcal{S}(D_0)}} + \mathbb{E}[\tilde{D} - D_0]\sigma\frac{\frac{\mathrm{d}}{\mathrm{d}D}H_{1,p,D_0}}{\sqrt{\mathcal{S}(D_0)}} - \frac{1}{2}\mathbb{E}[\tilde{D} - D_0]H_{1,p,D_0}\sigma^2\frac{1}{\sqrt{\mathcal{S}(D_0)}}\frac{\mathrm{d}}{\mathrm{d}\theta}\log\mathcal{S} \tag{112}$$

$$= \frac{H_{1,p,D_0} + \mathbb{E}[\tilde{D} - D_0]\frac{\mathrm{d}}{\mathrm{d}D}H_{1,p,D_0} - \frac{1}{2}\mathbb{E}[\tilde{D} - D_0]H_{1,p,D_0}\sigma\frac{\mathrm{d}}{\mathrm{d}\theta}\log\mathcal{J}}{\sqrt{\mathcal{J}(\theta)}} \tag{113}$$

$$=: \frac{-C_{1,p,D,\theta,F,\sigma}}{\sqrt{\mathcal{J}(\theta)}} \tag{114}$$

where $C_{1,p,D,\theta,F,\sigma} > 0$, $C_{1,p,D,\theta,F,\sigma} = \Theta(1+\sigma)$ in $\sigma$, $\lim_{D\to\infty} C_{1,p,D,\theta,F,\sigma} = 0$. Next, we consider

$$\mathbb{E}[C_{2,p,D}\frac{1}{\mathcal{J}}(\log p_{prior})']$$
$$= C_{2,p,D}\frac{1}{\mathcal{J}}(\log p_{prior})' + \mathbb{E}[\tilde{D} - D_0]\sigma\left[\frac{\mathrm{d}}{\mathrm{d}\theta}C_{2,p,D}\right]\frac{1}{\mathcal{J}}\frac{\mathrm{d}}{\mathrm{d}\theta}(\log p_{prior}) + \mathbb{E}[\tilde{D} - D_0]C_{2,p,D}\sigma\frac{\mathrm{d}}{\mathrm{d}\theta}\left[\frac{1}{\mathcal{J}}\frac{\mathrm{d}}{\mathrm{d}\theta}(\log p_{prior})\right]$$

The second and third term have higher order. The same holds for the repulsion term. In sum, and absorbing the ordinary encoding bias into $C_{3,p,D}$, we obtain the bias

$$\mathbb{E}[\widehat{\theta}] - \theta = \underbrace{-\frac{C_{1,p,D,\theta,F,\sigma}}{\sqrt{\mathcal{J}}}}_{\textit{Regression}} + C_{2,p,D}\underbrace{\frac{1}{\mathcal{J}}(\log p_{prior})'}_{\textit{Prior Attraction}} + C_{3,p,D}\underbrace{\frac{p+2}{4}\left(\frac{1}{\mathcal{J}}\right)'}_{\textit{Likelihood Repulsion}} + O(\sigma^3) \tag{115}$$

where all quantities are evaluated at $\theta$. This concludes the proof of the theorem.

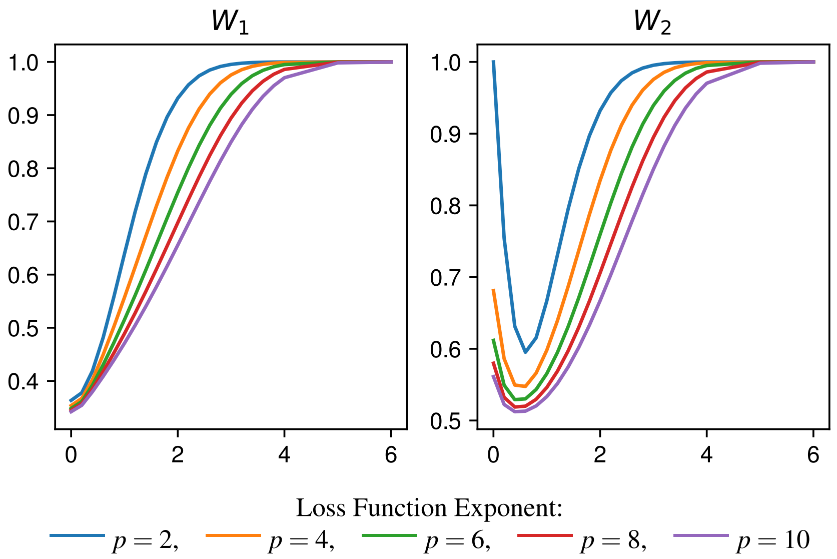

Figure S47: Behavior of the boundary-induced factors $W_1$ and $W_2$ in Equation 108, as a function of the effective distance $D := Q/\sigma$ (x-axis) and the loss function exponent (p=2,4,6,8,10). Both factors are bounded between $[0,1]$, and rapidly converge to 1 as the effective distance increases. As in Figure S46, the effect of the boundary is increased for higher exponents.

#### S3.4 Relationship between Threshold and Bias

Here, we show that the lawful relation between discrimination threshold and perceptual bias found by Wei and Stocker [32] holds if and only if prior and encoding are linked by a power law. Assuming based on the Cramer-Rao bound that the discrimination threshold is proportional to the inverse Fisher information, Wei and Stocker [32] equivalently formalize the law as

$$\mathbb{E}[\widehat{\theta}] - \theta \propto \left(\frac{1}{\mathcal{J}}\right)' \tag{116}$$

where the constant of proportionality may be positive (repulsive) or negative (attractive). Recall that by Theorem 1 ($p \geq 1$, analogous argument for $p = 0$)

$$\mathbb{E}[\widehat{\theta}] - \theta = \underbrace{\frac{1}{\mathcal{J}}(\log p_{prior})' +}_{Prior\ Attraction} \underbrace{\frac{p+2}{4}\left(\frac{1}{\mathcal{J}}\right)'}_{Likelihood\ Repulsion} ; \tag{117}$$

up to higher-order approximation error. Then the law holds if and only if

$$\frac{1}{\mathcal{J}}(\log p_{prior})' \propto \left(\frac{1}{\mathcal{J}}\right)'$$

Multiplying both sideas by $\mathcal{J}$, this holds if and only if

$$(\log p_{prior})' \propto (\log \mathcal{J})'$$

which holds if and only if, for some $a, b \in \mathbb{R}$,

$$a + b \log p_{prior} = \log \mathcal{J}$$

which is equivalent to

$$\mathcal{J}(\theta) = \exp(a) \cdot (p_{prior}(\theta))^b$$

i.e., prior and encoding are linked by a power law.

#### S3.5 Results under Gaussian model

Here, we derive the estimation bias under an alternative model sometimes used in the literature, showing that qualitative conclusions from Theorem 1 carry over to that model. Our model (Equation 1 in the main paper) accounts for the neural encoding process using a nonlinear map $F$ noisily mapping the stimulus into a sensory space, an abstraction of the neural encoding in the brain [e.g. 31, 32, 22, 10, 18, 21, 9, 34]. An alternative modeling approach sometimes employed [e.g. 7, 28, 15, 12, 19] forgoes such a nonlinear encoding, and instead assumes that Gaussian noise directly applies to the stimulus, but with a stimulus-dependent variance. Here, we show that a result analogous to Theorem 1 also holds in this model.[24] With a noise parameter $t > 0$, this Gaussian model can be written as:

$$m = \theta + \sqrt{t}\sigma_\theta \delta \tag{118}$$

where $\delta \sim N(0,1)$, and $\sigma_\theta = \frac{1}{\sqrt{t\mathcal{J}(\theta)}}$ is the inverse resource allocation at the stimulus $\theta$. For this model, the Fisher information is

$$\mathcal{J}(\theta) = \frac{1}{t\sigma_\theta^2} + \left(\frac{\mathrm{d}}{\mathrm{d}\theta}\log\sigma_\theta\right)^2$$

If noise $t$ is small, this is dominated by the first term (known as the "linear Fisher information"), and

$$\frac{1}{\mathcal{J}(\theta)} = t\sigma_\theta^2 + O(t^2) \tag{119}$$

We focus on the case of an $L^{2q}$ ($q > 0$ integer) loss function, showing the following decomposition analogous to Theorem 1:

**Theorem S6.** *For the $L^{2q}$ loss ($q > 0$ an integer), the bias is ($p := 2q$):*

$$t\sigma_\theta^2 \frac{\mathrm{d}}{\mathrm{d}\theta}\log\left[p_{prior}(\theta)\right] + t\frac{q}{2}\frac{\mathrm{d}}{\mathrm{d}\theta}\sigma_\theta^2 + O(t^2) \tag{120}$$

Considering

$$\mathcal{J}(\theta) = \frac{1}{t\sigma_\theta^2} + \left(\frac{\mathrm{d}}{\mathrm{d}\theta}\log\sigma_\theta\right)^2 \tag{121}$$

and hene $\frac{1}{\mathcal{J}} \approx t\sigma_\theta^2$ as $t \to 0$, this equals

$$\boxed{\underbrace{\frac{1}{\mathcal{J}(\theta)}\frac{\mathrm{d}}{\mathrm{d}\theta}\log\left[p_{prior}(\theta)\right]}_{\textit{Prior Attraction}} + \underbrace{\frac{p}{2}\left(\frac{\mathrm{d}}{\mathrm{d}\theta}\frac{1}{\mathcal{J}(\theta)}\right)}_{\textit{Likelihood Repulsion}} + O(t^2)}$$

Further, the bias at $p = 0$ was shown to be $\frac{1}{\mathcal{J}}(\log p_{prior})'$ by [28] using arguments equivalent to those in Section S3.1.4.4. The comparison between the biases under this model and our models is summarized in Table S2. Overall, the resulting biases differ only in the scaling of likelihood repulsion with the exponent. The key qualitative conclusions remain unchanged: first, biases decompose additively into prior attraction and likelihood repulsion; second, both are propotional to the noise variance; third, likelihood repulsion but not prior attraction increases monotonically with the loss function exponent.

We proceed to proving Theorem S6.

[24] Another variant is proposed by Prat-Carrabin and Woodford [24, 23], whereby stimuli $\theta$ are encoded by many independent random signals $X_1, \ldots, X_n$; this allows them to exploit the known asymptotic bias of the MLE to compute the bias at $p = 2$. The resulting bias equals that obtained under our model and the Gaussian model.

| Exponent | Ours | Gaussian |
|---|---|---|
| 0 | $\frac{1}{\mathcal{J}}(\log p_{prior})' + \frac{1}{4}\left(\frac{1}{\mathcal{J}}\right)'$ | $\frac{1}{\mathcal{J}}(\log p_{prior})'$ |
| 2 | $\frac{1}{\mathcal{J}}(\log p_{prior})' + \left(\frac{1}{\mathcal{J}}\right)'$ | $\frac{1}{\mathcal{J}}(\log p_{prior})' + \left(\frac{1}{\mathcal{J}}\right)'$ |
| 4 | $\frac{1}{\mathcal{J}}(\log p_{prior})' + \frac{3}{2}\left(\frac{1}{\mathcal{J}}\right)'$ | $\frac{1}{\mathcal{J}}(\log p_{prior})' + 2\left(\frac{1}{\mathcal{J}}\right)'$ |
| 6 | $\frac{1}{\mathcal{J}}(\log p_{prior})' + 2\left(\frac{1}{\mathcal{J}}\right)'$ | $\frac{1}{\mathcal{J}}(\log p_{prior})' + 3\left(\frac{1}{\mathcal{J}}\right)'$ |
| 8 | $\frac{1}{\mathcal{J}}(\log p_{prior})' + \frac{5}{2}\left(\frac{1}{\mathcal{J}}\right)'$ | $\frac{1}{\mathcal{J}}(\log p_{prior})' + 4\left(\frac{1}{\mathcal{J}}\right)'$ |

Table S2: Comparing bias in our model (Theorem 1) and in the Gaussian model assumed in some prior work (Section S3.5). At $p = 2$, the bias is equivalent in both models, recovering also the expression derived by Prat-Carrabin and Woodford [24]. The Gaussian model shows a steeper increase of repulsion with the exponent, but the overall pattern of a monotonic increase of repulsion with the exponent, whereas prior attraction is independent of the exponent, holds in both models.

*Proof.* The overall proof method for deriving the bias is analogous to the arguments in Section S3.1.4, but all integrals are now carried out in the stimulus space. We first note that encoding bias is zero in the Gaussian model. Now, to compute the decoding bias, as in Eq. 56, we want to solve the following for $\widehat{\theta}$ [WLOG $m = 0$]:

$$
\begin{aligned}
0 &= \int (\widehat{\theta} - \theta)^{2q-1} p_{prior}(\theta) N(\theta, t) \exp\left(-\frac{1}{t}\frac{(m-\theta)^2}{2\sigma_\theta^2}\right) d\theta \\
&= \int (\widehat{\theta} - \theta)^{2q-1} p_{prior}(\theta) N(\theta, t) \exp\left(-\frac{1}{t}\left[\frac{(m-\theta)^2}{2\sigma_\theta^2}\right]\right) d\theta \\
&= \int (\widehat{\theta} - \theta)^{2q-1} p_{prior}(\theta) N(\theta, t) \exp\left(-\frac{(m-\theta)^2}{t}\left[\frac{1}{2\sigma_\theta^2}\right]\right) d\theta \\
&= \int (\widehat{\theta} - \theta)^{2q-1} p_{prior}(\theta) N(\theta, t) \exp\left(-\frac{(m-\theta)^2}{t}\left[\frac{1}{2\sigma_\theta^2} + \theta\frac{\mathrm{d}}{\mathrm{d}\theta}\frac{1}{2\sigma_\theta^2} + \frac{1}{2}\theta^2\frac{\mathrm{d}^2}{\mathrm{d}\theta^2}\frac{1}{2\sigma_\theta^2} + \frac{1}{6}\theta^3\frac{\mathrm{d}^3}{\mathrm{d}\theta^3}\frac{1}{2\sigma_\theta^2} + \dots\right]\right) d\theta \\
&= \int (\widehat{\theta} - \theta)^{2q-1} p_{prior}(\theta) N(\theta, t) \exp\left(-\frac{(m-\theta)^2}{t}\left[\frac{1}{2\sigma_\theta^2}\right]\right) \exp\left(-\frac{(m-\theta)^2}{t}\left[\theta\frac{\mathrm{d}}{\mathrm{d}\theta}\frac{1}{2\sigma_\theta^2} + \frac{1}{2}\theta^2\frac{\mathrm{d}^2}{\mathrm{d}\theta^2}\frac{1}{2\sigma_\theta^2} + \theta^3\frac{1}{6}\frac{\mathrm{d}^3}{\mathrm{d}\theta^3}\frac{1}{2\sigma_\theta^2} + \dots\right]\right) d\theta \\
&= \int (\widehat{\theta} - \theta)^{2q-1} p_{prior}(\theta) \frac{1}{\sqrt{2\pi t\sigma_\theta^2}} \exp\left(-\frac{\theta^2}{t}\left[\frac{1}{2\sigma_\theta^2}\right]\right) \exp\left(-\frac{\theta^3}{t}\left[\frac{\mathrm{d}}{\mathrm{d}\theta}\frac{1}{2\sigma_\theta^2} + \dots\right]\right) d\theta \\
&= \int (\widehat{\theta} - \theta)^{2q-1} p_{prior}(\theta) \frac{1}{\sqrt{2\pi t\sigma_\theta^2}} \exp\left(-\frac{\theta^2}{t}\left[\frac{1}{2\sigma_\theta^2}\right]\right) (1 - \frac{\theta^3}{2t}\left[\frac{\mathrm{d}}{\mathrm{d}\theta}\frac{1}{\sigma_\theta^2} + \dots\right] + \dots) d\theta \\
&\propto \int (\widehat{\theta} - \theta)^{2q-1} p_{prior}(\theta) \frac{1}{\sqrt{\sigma_\theta^2}} \left(1 - \frac{\theta^3}{2t}\left[\frac{\mathrm{d}}{\mathrm{d}\theta}\frac{1}{\sigma_\theta^2}\right]\right) \exp\left(-\frac{\theta^2}{t}\left[\frac{1}{2\sigma_\theta^2}\right]\right) d\theta + h.o.t. \\
&\propto \int (\widehat{\theta} - \theta)^{2q-1} \left(1 - \frac{\theta^3}{2t}\left[\frac{\mathrm{d}}{\mathrm{d}\theta}\frac{1}{\sigma_\theta^2}\right]\right) \exp\left(-\frac{\theta^2}{t}\left[\frac{1}{2\sigma_\theta^2}\right]\right) d\theta \\
&\quad + \int (\widehat{\theta} - \theta)^{2q-1} \theta \frac{\mathrm{d}}{\mathrm{d}\theta} \log\left[\frac{p_{prior}(\theta)}{\sqrt{\sigma_\theta^2}}\right] \left(1 - \frac{\theta^3}{2t}\left[\frac{\mathrm{d}}{\mathrm{d}\theta}\frac{1}{\sigma_\theta^2}\right]\right) \exp\left(-\frac{\theta^2}{t}\left[\frac{1}{2\sigma_\theta^2}\right]\right) d\theta + h.o.t. \\
&= \sum_{i=0}^{2q-1} \int \binom{2q-1}{i} \widehat{\theta}^i (-1)^{2q-1-i} \theta^{2q-1-i} \left(1 - \frac{\theta^3}{2t}\left[\frac{\mathrm{d}}{\mathrm{d}\theta}\frac{1}{\sigma_\theta^2}\right]\right) \exp\left(-\frac{\theta^2}{t}\left[\frac{1}{2\sigma_\theta^2}\right]\right) d\theta
\end{aligned}
$$

$$+\sum_{i=0}^{2q-1}\int\binom{2q-1}{i}\widehat{\theta}^i(-1)^{2q-1-i}\theta^{2q-i}\frac{\mathrm{d}}{\mathrm{d}\theta}\log\left[\frac{p_{prior}(\theta)}{\sqrt{\sigma_\theta^2}}\right]\left(1-\frac{\theta^3}{2t}\left[\frac{\mathrm{d}}{\mathrm{d}\theta}\frac{1}{\sigma_\theta^2}\right]\right)\exp\left(-\frac{\theta^2}{t}\left[\frac{1}{2\sigma_\theta^2}\right]\right)d\theta+h.o.t.$$

As in Section S3.1.4.1, the lowest orders in $t$ are attained at $i=0,1$:

$$\begin{aligned}
&-\int\theta^{2q-1}\left(1-\frac{\theta^3}{2t}\left[\frac{\mathrm{d}}{\mathrm{d}\theta}\frac{1}{\sigma_\theta^2}\right]\right)\exp\left(-\frac{\theta^2}{t}\left[\frac{1}{2\sigma_\theta^2}\right]\right)d\theta\\
&-\int\theta^{2q}\frac{\mathrm{d}}{\mathrm{d}\theta}\log\left[\frac{p_{prior}(\theta)}{\sqrt{\sigma_\theta^2}}\right]\left(1-\frac{\theta^3}{2t}\left[\frac{\mathrm{d}}{\mathrm{d}\theta}\frac{1}{\sigma_\theta^2}\right]\right)\exp\left(-\frac{\theta^2}{t}\left[\frac{1}{2\sigma_\theta^2}\right]\right)d\theta\\
&+\int(2q-1)\widehat{\theta}\theta^{2q-2}\left(1-\frac{\theta^3}{2t}\left[\frac{\mathrm{d}}{\mathrm{d}\theta}\frac{1}{\sigma_\theta^2}\right]\right)\exp\left(-\frac{\theta^2}{t}\left[\frac{1}{2\sigma_\theta^2}\right]\right)d\theta\\
&+\int(2q-1)\widehat{\theta}\theta^{2q-1}\frac{\mathrm{d}}{\mathrm{d}\theta}\log\left[\frac{p_{prior}(\theta)}{\sqrt{\sigma_\theta^2}}\right]\left(1-\frac{\theta^3}{2t}\left[\frac{\mathrm{d}}{\mathrm{d}\theta}\frac{1}{\sigma_\theta^2}\right]\right)\exp\left(-\frac{\theta^2}{t}\left[\frac{1}{2\sigma_\theta^2}\right]\right)d\theta+h.o.t.
\end{aligned}$$

which evaluates to

$$\begin{aligned}
0=&\frac{1}{2t}\left[\frac{\mathrm{d}}{\mathrm{d}\theta}\frac{1}{\sigma_\theta^2}\right]\int\theta^{2q+2}\exp\left(-\frac{\theta^2}{t}\left[\frac{1}{2\sigma_\theta^2}\right]\right)d\theta\\
&-\frac{\mathrm{d}}{\mathrm{d}\theta}\log\left[\frac{p_{prior}(\theta)}{\sqrt{\sigma_\theta^2}}\right]\int\theta^{2q}\exp\left(-\frac{\theta^2}{t}\left[\frac{1}{2\sigma_\theta^2}\right]\right)d\theta\\
&+(2q-1)\widehat{\theta}\int\theta^{2q-2}\exp\left(-\frac{\theta^2}{t}\left[\frac{1}{2\sigma_\theta^2}\right]\right)d\theta\\
&-(2q-1)\widehat{\theta}\frac{1}{2t}\left[\frac{\mathrm{d}}{\mathrm{d}\theta}\frac{1}{\sigma_\theta^2}\right]\frac{\mathrm{d}}{\mathrm{d}\theta}\log\left[\frac{p_{prior}(\theta)}{\sqrt{\sigma_\theta^2}}\right]\int\theta^{2q+2}\exp\left(-\frac{\theta^2}{t}\left[\frac{1}{2\sigma_\theta^2}\right]\right)d\theta+h.o.t.
\end{aligned}$$

and hence

$$\begin{aligned}
0=&\frac{1}{2t}\left[\frac{\mathrm{d}}{\mathrm{d}\theta}\frac{1}{\sigma_\theta^2}\right]\left[t\sigma_\theta^2\right]^{q+1}(2q+1)!!\\
&-\frac{\mathrm{d}}{\mathrm{d}\theta}\log\left[\frac{p_{prior}(\theta)}{\sqrt{\sigma_\theta^2}}\right]\left[t\sigma_\theta^2\right]^{q}(2q-1)!!\\
&+(2q-1)\widehat{\theta}\left[t\sigma_\theta^2\right]^{q-1}(2q-3)!!\\
&-(2q-1)\widehat{\theta}\frac{1}{2t}\left[\frac{\mathrm{d}}{\mathrm{d}\theta}\frac{1}{\sigma_\theta^2}\right]\frac{\mathrm{d}}{\mathrm{d}\theta}\log\left[\frac{p_{prior}(\theta)}{\sqrt{\sigma_\theta^2}}\right]\left[t\sigma_\theta^2\right]^{q+1}(2q+1)!!+h.o.t.
\end{aligned}$$

The last term has higher order; the equation simplifies to

$$0 = \frac{2q+1}{2}\left[\frac{\mathrm{d}}{\mathrm{d}\theta}\frac{1}{\sigma_\theta^2}\right] t\left[\sigma_\theta^2\right]^2 - \frac{\mathrm{d}}{\mathrm{d}\theta}\log\left[\frac{p_{prior}(\theta)}{\sqrt{\sigma_\theta^2}}\right]\left[t\sigma_\theta^2\right] + \widehat{\theta} + h.o.t.$$

and thus

$$\begin{aligned}
\theta &= -\frac{2q+1}{2}\left[\frac{\mathrm{d}}{\mathrm{d}\theta}\frac{1}{\sigma_\theta^2}\right] t\left[\sigma_\theta^2\right]^2 + \frac{\mathrm{d}}{\mathrm{d}\theta}\log\left[\frac{p_{prior}(\theta)}{\sqrt{\sigma_\theta^2}}\right]\left[t\sigma_\theta^2\right] + h.o.t. \\
&= \frac{2q+1}{2}\left[\frac{\mathrm{d}}{\mathrm{d}\theta}\sigma_\theta^2\right] t - t\frac{1}{2}\frac{\mathrm{d}}{\mathrm{d}\theta}\sigma_\theta^2 + \frac{\mathrm{d}}{\mathrm{d}\theta}\log\left[p_{prior}(\theta)\right]\left[t\sigma_\theta^2\right] + h.o.t. \\
&= \frac{2q}{2} t\left[\frac{\mathrm{d}}{\mathrm{d}\theta}\sigma_\theta^2\right] + \frac{\mathrm{d}}{\mathrm{d}\theta}\log\left[p_{prior}(\theta)\right]\left[t\sigma_\theta^2\right] + h.o.t. \\
&= \frac{p}{2} t\left[\frac{\mathrm{d}}{\mathrm{d}\theta}\sigma_\theta^2\right] + \frac{\mathrm{d}}{\mathrm{d}\theta}\log\left[p_{prior}(\theta)\right]\left[t\sigma_\theta^2\right] + h.o.t.
\end{aligned}$$

This completes the proof. □

[10] Matthias Fritsche, Eelke Spaak, and Floris P. de Lange. A bayesian and efficient observer model explains concurrent attractive and repulsive history biases in visual perception. *bioRxiv*, 2020.

[18] Thomas Langlois, Nori Jacoby, Jordan W. Suchow, and T. Griffiths. Serial reproduction reveals the geometry of visuospatial representations. *Proceedings of the National Academy of Sciences of the United States of America*, 118, 2021.

[19] Michael J. Morais and Jonathan W. Pillow. Power-law efficient neural codes provide general link between perceptual bias and discriminability. In Samy Bengio, Hanna M. Wallach, Hugo Larochelle, Kristen Grauman, Nicolò Cesa-Bianchi, and Roman Garnett, editors, *Advances in Neural Information Processing Systems 31: Annual Conference on Neural Information Processing Systems 2018, NeurIPS 2018, December 3-8, 2018, Montréal, Canada*, pages 5076–5085, 2018. URL `https://proceedings.neurips.cc/paper/2018/hash/61d77652c97ef636343742fc3dcf3ba9-Abstract.html`.

[20] Adam Paszke, Sam Gross, Francisco Massa, Adam Lerer, James Bradbury, Gregory Chanan, Trevor Killeen, Zeming Lin, Natalia Gimelshein, Luca Antiga, Alban Desmaison, Andreas Köpf, Edward Z. Yang, Zachary DeVito, Martin Raison, Alykhan Tejani, Sasank Chilamkurthy, Benoit Steiner, Lu Fang, Junjie Bai, and Soumith Chintala. Pytorch: An imperative style, high-performance deep learning library. In Hanna M. Wallach, Hugo Larochelle, Alina Beygelzimer, Florence d'Alché-Buc, Emily B. Fox, and Roman Garnett, editors, *Advances in Neural Information Processing Systems 32: Annual Conference on Neural Information Processing Systems 2019, NeurIPS 2019, December 8-14, 2019, Vancouver, BC, Canada*, pages 8024–8035, 2019. URL `https://proceedings.neurips.cc/paper/2019/hash/bdbca288fee7f92f2bfa9f7012727740-Abstract.html`.

[21] Frederike H. Petzschner and Stefan Glasauer. Iterative bayesian estimation as an explanation for range and regression effects: A study on human path integration. *The Journal of Neuroscience*, 31:17220 – 17229, 2011.

[22] Rafael Polanía, Michael Woodford, and Christian C. Ruff. Efficient coding of subjective value. *Nature neuroscience*, 22:134 – 142, 2018.

[23] Arthur Prat-Carrabin and Michael Woodford. Efficient coding of numbers explains decision bias and noise. *bioRxiv*, 2020.

[24] Arthur Prat-Carrabin and Michael Woodford. Bias and variance of the bayesian-mean decoder, 2021.

[25] Evan D. Remington, Tiffany V Parks, and Mehrdad Jazayeri. Late bayesian inference in mental transformations. *Nature Communications*, 9, 2018.

[26] Walter Rudin. *Principles of mathematical analysis*. 1964.

[33] Yang Xiang, Thomas Graeber, Benjamin Enke, and Samuel J. Gershman. Confidence and central tendency in perceptual judgment. *Attention, perception & psychophysics*, 2021.

[34] Hang Zhang, Xiangjuan Ren, and Laurence T. Maloney. The bounded rationality of probability distortion. *Proceedings of the National Academy of Sciences of the United States of America*, 117:22024 – 22034, 2020.
